# Thermal adaptation in worldwide collections of a major fungal pathogen

**DOI:** 10.1101/2024.09.12.612681

**Authors:** Silvia Miñana-Posada, Cécile Lorrain, Bruce A. McDonald, Alice Feurtey

## Abstract

Adaptation to new climates poses a significant challenge for plant pathogens during range expansion, highlighting the importance of understanding their response to climate to accurately forecast future disease outbreaks. The wheat pathogen *Zymoseptoria tritici* is ubiquitous across most wheat production regions distributed across diverse climate zones. We explored the genetic architecture of thermal adaptation using a global collection of 411 *Z. tritici* strains that were phenotyped across a wide range of temperatures and then included in a genome-wide association study. Our analyses provided evidence for local thermal adaptation in *Z. tritici* populations worldwide, with a significant positive correlation between bioclimatic variables and optimal growth temperatures. We also found a high variability in thermal performance among *Z. tritici* strains coming from the same field populations, reflecting the high evolutionary potential of this pathogen at the field scale. We identified 69 genes putatively involved in thermal adaptation, including one high-confidence candidate potentially involved in cold adaptation. These results highlight the complex polygenic nature of thermal adaptation in *Z. tritici* and suggest that this pathogen is likely to adapt well when confronted with climate change.

## Introduction

Climate change and intensification of the global food trade significantly contribute to the spread and increased damage caused by endemic and emerging plant diseases (Ristaino et al. 2021). Since the 1960s, the global increase in temperature has caused a poleward shift in the distribution of most plant pathogens (Bebber, Ramotowski, and Gurr 2013).

Temperature increases are also associated with more severe and frequent epidemics, as well as the emergence of new pathogens (Bebber 2015; McDonald and Stukenbrock 2016). For instance, during the last twenty years, high-temperature-adapted and highly aggressive strains of the fungal wheat pathogen *Puccinia striiformis* f. sp. *tritici* have spread across wheat fields worldwide (de Vallavieille-Pope et al. 2018; Milus, Kristensen, and Hovmøller 2009; Walter et al. 2016). The rice pathogen *Pyricularia oryzae* showed increased virulence associated with high-temperature exposure in controlled experiments (Onaga et al. 2017). Investigating temperature adaptation in fungal pathogens will be crucial to understand how these organisms will respond to climate change. In agroecosystems, widely distributed crop pathogens encounter diverse environments, including spatial and temporal temperature patterns that are likely to be a major driving force for local adaptation. Local adaptation is determined by the interplay between natural selection, gene flow, and specific events such as founder effects or population extinctions. Evidence of local thermal adaptation has already been described for some broadly distributed plant pathogens (Boixel, Chelle, and Suffert 2022; Mariette et al. 2016; Mboup et al. 2012; Pereira et al. 2020; Stefansson, McDonald, and Willi 2013; Zhan and McDonald 2011).

Despite its importance, our understanding of the genetic basis of thermal adaptation in fungal plant pathogens remains limited. This can be partially attributed to the fact that the temperature response in fungi involves multiple genes affecting many morphological, cellular, and molecular processes (Blasi et al. 2015; Feurtey et al. 2023; Foulongne-Oriol et al. 2014; Maclean et al. 2017; Lendenmann et al. 2016; Tralamazza et al. 2024; Weiss et al. 2018). Fungi respond to temperature stress through morphological changes (Berman 2006; Keuenhof et al. 2021; van den Brule et al. 2020), alterations in membrane composition (Covino et al. 2016; Leach and Cowen 2014), the production of specific metabolites such as pyruvate (Zhang et al., 2018) and stabilization of proteins through the expression of chaperone proteins like heat shock proteins (Masser et al. 2019; Wu et al. 2016). Additionally, the thermal stress response intertwines with other stress response pathways, such as oxidative and osmotic stress response pathways, which involve the production of antioxidants and activation of the high osmolarity glycerol (HOG) pathway (Kubicek and Druzhinina 2007; Yaakoub et al. 2022). The HOG pathway plays a crucial role in cold stress response in Antarctic fungi (Kostadinova et al. 2011) and in heat stress response in *Saccharomyces cerevisiae* and *Aspergillus nidulans* (Panadero et al. 2006). Transcription factors are also key regulators of temperature responses, particularly heat shock transcription factors (Hsf) that govern the expression of heat shock proteins, as demonstrated for Hsf1 in *S. cerevisiae* and *Candida albicans* (Veri, Robbins, and Cowen 2018). Other transcription factors, such as MoSfl1 in *P. oryzae*, can regulate pathogenicity-related genes and responses to elevated temperatures and melanization (G. Li et al. 2011). Thermal adaptation mechanisms have barely been explored in fungal plant pathogens. Little is known about the molecular mechanisms underlying responses to stressful temperature conditions or potential differences in the genetic architecture of temperature adaptation among populations of fungal plant pathogens from diverse origins.

The fungal pathogen *Zymoseptoria tritici* is the causal agent of Septoria tritici blotch on wheat, a disease capable of causing yield losses of up to 50% in susceptible wheat cultivars without fungicide treatment (Jalli et al. 2020) and up to 10% in fungicide-treated fields across Europe (Fones and Gurr 2015). This pathogen emerged through host-tracking during the domestication of wheat in the Middle East (Stukenbrock et al. 2007). As wheat cultivation spread globally, *Z. tritici* became widely distributed, particularly in temperate climates (Petit-Houdenot, Lebrun, and Scalliet 2021; Savary et al. 2019). Previous studies on this pathogen’s thermal responses revealed high variation between and within populations (Boixel, Chelle, and Suffert 2022; Zhan and McDonald 2011). Quantitative Trait Loci (QTL) mapping in Swiss strains identified 18 QTLs related to growth at two temperatures and revealed pleiotropic effects with morphology and melanization (Lendenmann et al. 2016). Recent genome-environment association studies using global collections of *Z. tritici* revealed nearly nine hundred variants associated with bioclimatic variables, including one locus on chromosome 7 harboring a massive transposable element associated with the annual temperature (Feurtey et al. 2023; Tralamazza et al. 2024). Despite these promising findings, deeper exploration using extensive phenotype datasets is needed to reveal candidate genes responsible for temperature adaptation.

Here, we gathered a global collection of 411 strains of *Z. tritici,* measured their growth across a wide range of temperatures, and analyzed their genomes to investigate the patterns of local thermal adaptation by (i) exploring the population structure and sources of genetic variance between populations and genetic clusters, (ii) characterizing the thermal response of the pathogen through a series of growth variables estimated from growth phenotypes under five temperature conditions, and (iii) identifying genetic variants and candidate genes associated with temperature adaptation using genome-wide association studies (GWAS).

## Results

### Population structure and genetic diversity in a worldwide collection of *Zymoseptoria tritici*

We obtained 1,029,154 high-confidence single nucleotide polymorphisms (SNPs) and small indels including 982,820 biallelic SNPs from a collection of 411 *Z. tritici* strains. Among these, 937,532 SNPs were located on the 13 core chromosomes. Admixture analysis revealed eight distinct genetic clusters, which broadly corresponded to the geographical origins of the strains (Figures 1A & 1B; Supplementary Figures S2B-D). Only 10 of the 411 strains (2.4%) could not be assigned to a specific cluster (Supplementary Figure S2F), most of them originating from Turkish and Ukrainian populations. We observed population differentiation at the continental level, with some notable exceptions (Figure 1A & 1B). For instance, Canadian strains formed a distinct cluster separate from other North American strains. Similarly, in South American populations, Argentinian and Uruguayan strains clustered together while Chilean strains formed a separate cluster (Figures 1A & 1B).

**Figure 1.**
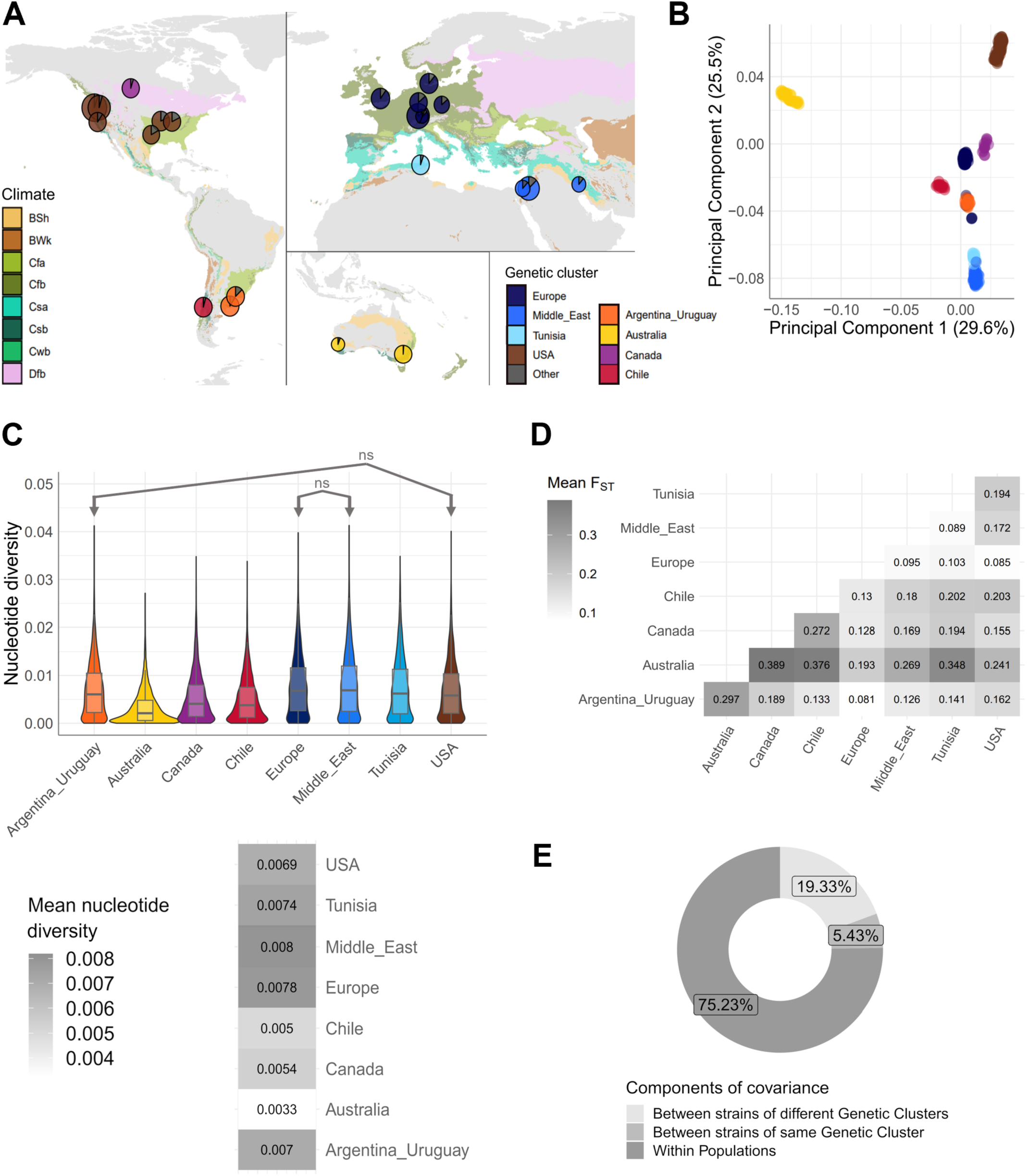
**A**) Geographic distribution of the genetic clusters identified among 411 *Zymoseptoria tritici* strains. Each pie represents the average cluster assignment coefficient for each population. Only populations with more than five strains are displayed. Each color represents a different genetic cluster but any fraction smaller than 20% was colored in grey to improve clarity. The map background is colored according to the Köppen-Geiger climate classes (BSh: dry, semi-arid or steppe, hot; BWk: dry, arid desert, cold; Cfa: temperate, no dry season, hot summer; Cfb: temperate, no dry season, warm summer; Csa: temperate, dry summer, hot summer; Csb: temperate, dry summer, warm summer; Cwb: temperate, dry winter, warm summer; Dfb: continental, no dry season, warm summer). **B**) Principal component analysis, showing the first and second components (PCs) based on a thinned genome-wide bi-allelic SNP dataset. Colors indicate the genomic clusters identified with the sNMF method. **C**) Nei and Li’s *π* estimating whole genome nucleotide diversity in 10 kb windows for each genetic cluster. Non-significant differences are identified with “ns” lines (Wilcoxon signed-rank tests Bonf. corr. *p*-value > 0.05). All other differences are significant. **D**) Mean genetic differentiation (F_ST_) between pairs of the different genetic clusters. **E**) Analysis of Molecular Variance using genetic clusters and geographical populations as the two levels of hierarchical subdivision.

We analyzed the genetic diversity within clusters and the differentiation between clusters. The highest mean genetic diversity (Nei and Li’s *π*) was observed in the Middle Eastern, European, and Tunisian clusters (0.007-0.008, Figure 1C). The Middle Eastern cluster exhibited the lowest genetic differentiation (Weir and Cockerham fixation index, F_ST_) with the Tunisian and European clusters (0.089 and 0.095, respectively; Figure 1D). The lowest pairwise genetic differentiation was found between the European cluster and both the North American (USA populations) and South American (Argentina and Uruguay populations) clusters (Figure 1D). In addition, the majority of molecular variance (75.2%) was attributed to within-population variation. However, a substantial portion of the variance (19.3%) was due to differences between populations from different genetic clusters (Figure 1E). These findings add further support to the high genetic diversity found within populations of *Z. tritici*.

### Local thermal adaptation in *Z. tritici* populations

To investigate whether local adaptation to temperature could be detected in the worldwide collection of *Z.tritici*, we compared 11 bioclimatic variables of the sampling locations to the 12 growth variables estimated based on the *in vitro* phenotyping of five temperature conditions. First, we examined the correlations among the 11 bioclimatic variables from the different sampling locations. We observed that 51 out of the 55 comparisons (93%) among the 11 bioclimatic variables are significant, with absolute correlation coefficients ranging from 0.17 to 0.97 (Pearson’s correlation, p-value < 0.05; see Supplementary Tables S7 and S8). The five most representative bioclimatic variables are shown in Figure 2A, including the annual mean temperature, the mean temperature of the warmest quarter, of the coldest quarter, of the wettest quarter, and of the driest quarter. The rest of the bioclimatic variables are shown in Supplementary Figure S3.

**Figure 2.**
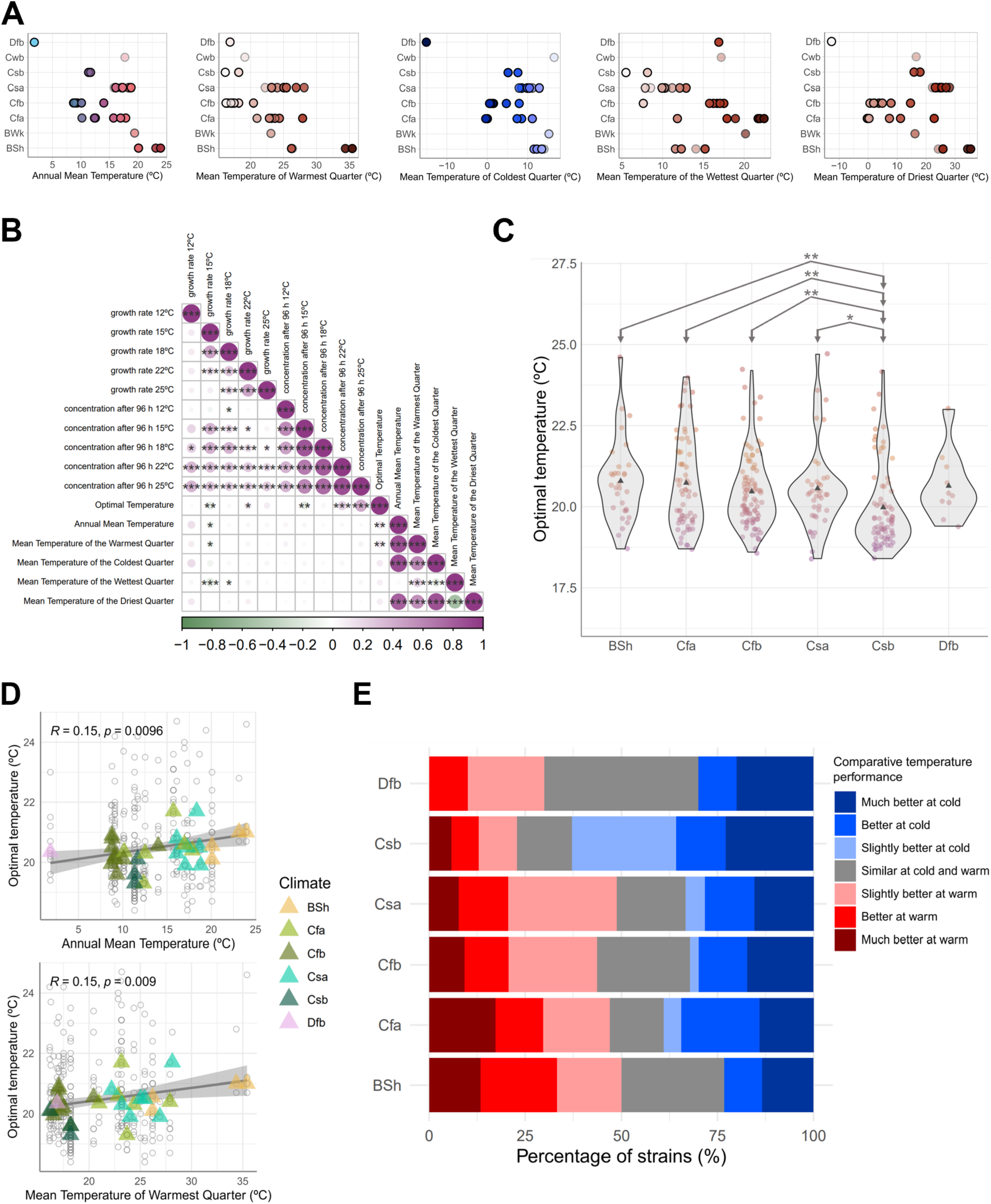
**A**) Variation of five bioclimatic variables for the 35 sampling locations (dots) classified by the Köppen-Geiger (KG) climate index (*y*-axis; BSh: dry, semi-arid or steppe, hot; BWk: dry, arid desert, cold; Cfa: temperate, no dry season, hot summer; Cfb: temperate, no dry season, warm summer; Csa: temperate, dry summer, hot summer; Csb: temperate, dry summer, warm summer; Cwb: temperate, dry winter, warm summer; Dfb: continental, no dry season, warm summer). **B**) Pearson’s correlations between a subset of the phenotypes measured and the bioclimatic variables. The color scale indicates the correlation coefficient, and the significance levels are represented by asterisks: * for *p*-value < 0.05, ** for *p*-value < 0.01, and *** for *p*-value < 0.001. **C**) Estimates of the optimal temperature for each climate class. Each point represents the optimal temperature for a strain. The significance levels of the Wilcoxon signed-rank tests with Bonferroni correction are represented by asterisks: * for *p*-value < 0.05, ** for *p*-value < 0.01, and *** for *p*-value < 0.001. Only the climates with more than five strains were represented. **D**) Linear regression between the optimal growth temperature (*y*-axis) and the annual mean temperature (*x*-axis, top panel) or the mean temperature of the warmest quarter (*x*-axis, bottom panel). Grey circles represent the isolates while the triangles show the population mean and are colored according to the KG climate. Only climates with more than five strains are represented. The rho (*R*) and the *p*-value of the Pearson’s correlations are displayed at the top of each plot. **E**) Percentage of strains per climate class that perform better at warmer temperatures or at colder temperatures through the comparison of the temperature sensitivity at 15°C and the temperature sensitivity at 25°C. Only climate classes with more than five strains are shown.

Next, we focused on the relationships between the 12 growth variables, including the four variables that summarized the overall growth over time (e.g. eAUC and growth rate; Supplementary Table S3; Figure 2C, Supplementary Figures S6A, D, K) and the eight variables that represented static points of growth (e.g. spore concentration 96 hours after inoculation, inflection time; Supplementary Figures S6B, C, E, F, G, H, I, J). Across the 1540 comparisons, we found that 837 (54%) were significantly correlated with absolute correlation coefficients ranging between 0.10 and 0.99 (Pearson’s corr. *p*-value < 0.05; Supplementary Tables S7 and S8). Among the four overall growth variables, we found that growth rates in warmer temperatures (18°C - 25°C) are significantly and positively correlated with each other (Pearson’s corr. *p*-value < 0.05), while growth rates in colder temperatures are only mildly (and non-significantly) correlated (Figure 2B). We also found that the growth rate and spore concentration after 96 hours are positively correlated, except for the extremes of the temperature conditions measured (12°C and 25°C) (Figure 2B).

Finally, we assessed the 616 comparisons between the 12 growth variables per temperature condition and the 11 bioclimatic variables. From these comparisons, 35 (6%) are significant with absolute correlation coefficients between 0.10 and 0.28 (Pearson’s corr. *p*-value < 0.05; Supplementary Tables S7 and S8). In particular, we observed significant positive correlations between the optimal growth temperature and the annual mean temperature and between the optimal growth temperature and the mean temperature of the warmest quarter (Figure 2D). This indicates that the *in vitro* thermal response growth is a suitable proxy for investigating the local thermal adaptation of *Z. tritici*. In addition, this result provides strong evidence for local thermal adaptation in *Z. tritici* populations.

This global collection of strains displayed a wide range of optimal growth temperatures within individual populations as well as moderate but significant differences among populations and climate classes. The estimated optimal growth temperatures ranged from 18.4°C to 24.7°C among the global populations (Figure 2C, Supplementary Table S3). We observed considerable variation in optimal growth temperature among strains within populations. We found that the variance in optimal growth temperatures was not significantly different between populations or climates (Levene’s test corrected for differences in population/climate size: p-values 0.96 and 0.97, respectively), meaning that the population/climate variables display a similar degree of dispersion in their optimal growth temperature estimates. However, we still detected significant differences in mean optimal growth temperatures between populations and climates (Kruskal-Wallis p-values: 3×10⁻⁶ and 2.6×10⁻⁴, respectively; corrected for differences in population/climate sample size: p-values 0.016 and 0.047, respectively). Pairwise comparisons between populations revealed significantly higher optimal growth temperatures of one Australian population (Wagga Wagga) compared to the three USA populations of Daviess County (Indiana), Corvallis (Oregon) sampled in 1990, and Corvallis sampled in 2015 (Wilcoxon signed-rank test *p*-value < 0.05; Supplementary Figure S5; Supplementary Table S4). At the climate level, we found that the significant differences in optimal temperatures are due mainly to the Csb climate strains having the lowest mean optimal temperature (Figure 2C; Supplementary Table S5). Interestingly, the Csb climate has the coldest temperatures during the warmest quarter (Figure 2A; Supplementary Table S6).

### Thermal specialization of individual strains explains the high variance found within local populations

We evaluated the temperature specialization of each strain by comparing the temperature sensitivity at high (25°C) and low (15°C) temperatures. We assigned each strain into a seven-category scale that reflects a strain’s comparative performance at warmer and colder temperatures (Figure 2E; Supplementary Figures S7-S9; Supplementary Table S9). All populations contained strains that performed similarly at both warmer and colder temperatures (i.e. generalist strains), as well as strains that performed better at either cold or warm temperatures (i.e. specialist strains), indicating that a broad variation in temperature specificity exists within each population (Figure 2E; Supplementary Figure S9). The climate Dfb had the highest percentage of generalist strains (40%), followed by BSh (26.7%), Cfb (24.1%), Csa (17.9%), Csb (14.3%) and Cfa (14.1%) (Figure 2E).

The two climates with the highest annual mean temperatures, BSh and Csa (Figure 3A; Supplementary Table S6), included 50% and 49% of strains performing better at warmer temperatures, respectively (Figure 2E). More than 75% of the strains from the Australian populations (Cfa and Csa) performed at least slightly better at warmer temperatures than at colder temperatures (Supplementary Figure S9). The Iranian population (BSh) also showed a high proportion of strains that performed better at warmer temperatures (>50%), with no strains performing better at colder temperatures (Supplementary Figure S9). The strains from Israel (BSh) generally preferred warmer temperatures (41%), although several strains (35%) performed better at colder temperatures (Supplementary Figure S9). The cold specialists were found in higher proportion in the Csb climate which comprises >50% of strains performing at least slightly better at colder temperatures (Figure 2E). The Daviess County (Indiana, USA; Cfa) population had the highest proportion of strains that performed better at colder temperatures, with no strains performing better at warmer temperatures (Supplementary Figure S9). This was followed by the two Corvallis (Oregon, USA) and Chile populations (Csb), where more than 50% of the strains performed better at colder temperatures (Figure 2E).

**Figure 3.**
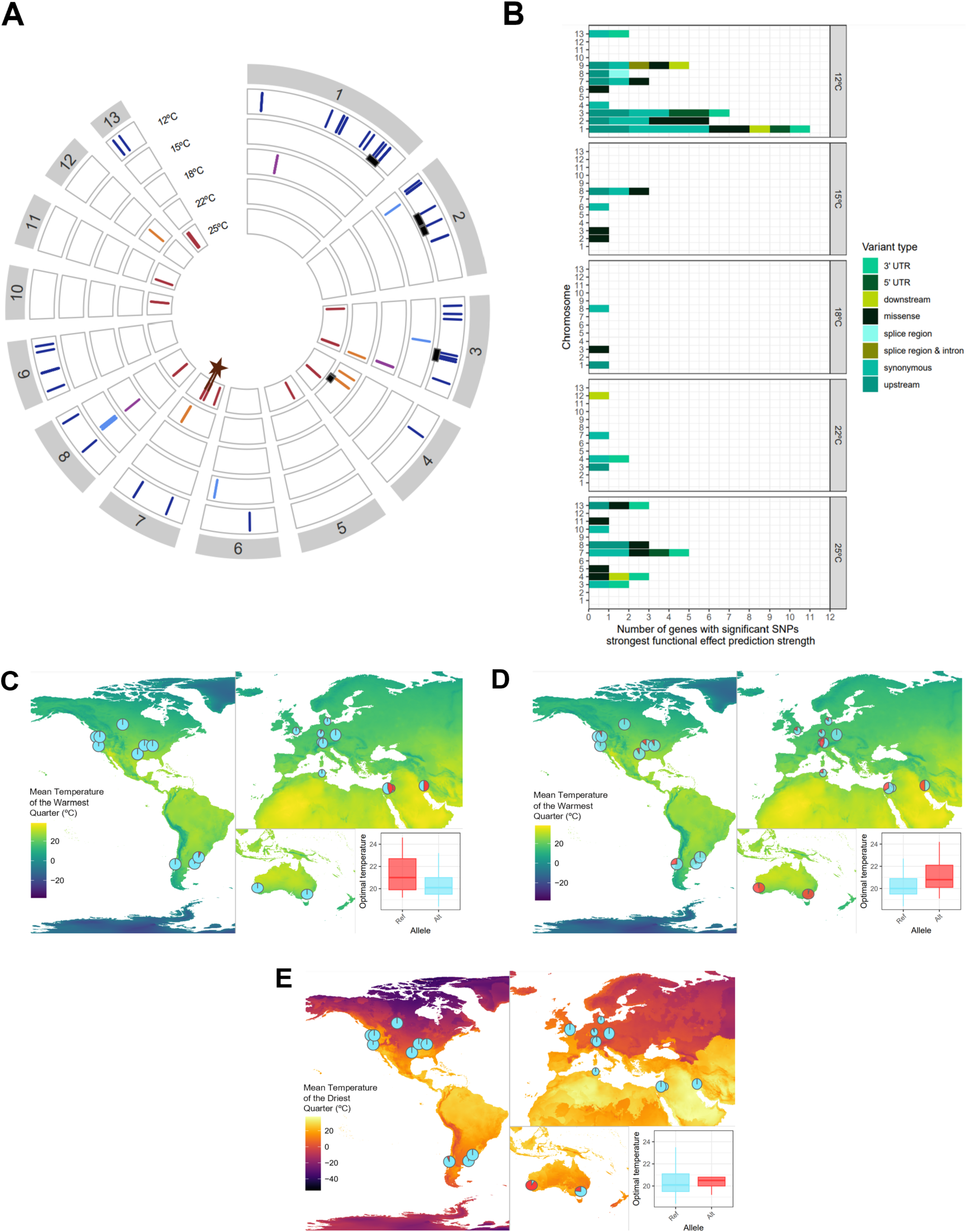
**A**) Location in the genome of variants significantly associated with any of the phenotypes measured at 12°C (dark blue), 15°C (light blue), 18°C (purple), 22°C (orange), and 25°C (red) on each of the 13 core chromosomes (grey boxes). The black rectangles represent previously identified quantitative trait loci linked to temperature adaptation that overlap with our significant variants (Lendenmann et al., 2016). The brown lines next to the star represent five genes, previously associated with bioclimatic variables (Feurtey et al., 2023). **B**) Number of genes (per chromosome and growth temperature) potentially impacted by significant variants. Each gene is classified according to the variant of the strongest effect as estimated by SnpEff. **C**) Geographic distribution of alleles for a variant associated with growth at cold temperature (12°C, chromosome 1, position 5569967). The associated box plot represents the distribution of the optimal growth temperature (°C) for strains carrying the reference allele (“Ref”) and the alternative allele (“Alt”). The alleles are colored according to the phenotype of the strains carrying them: blue for the cold-associated allele, and red for the warm-associated allele. The map background represents the mean temperature of the warmest quarter with more yellow for warmer areas, and more blue for colder areas. Only populations with more than five strains are shown. **D**) Same graph for a variant associated with growth at 22°C (chromosome 4, position 1074639). **E**) Same graph for a variant associated with growth at 25°C (chromosome 7, position 1512102). The map background represents the mean temperature of the driest quarter. The more yellow the area, the warmer, and the more purple the area, the colder.

We also explored the differences in temperature specialization between populations sampled in different years from the same field. Two Corvallis populations (Oregon, USA) were sampled from the same wheat cultivars in the same naturally infected field, once in 1990 and again in 2015 (Supplementary Table S1). We observed that the 2015 population had a slightly higher proportion of strains that performed better at colder temperatures (70%) compared to the 1990 population (62%) (Supplementary Figure S9). The proportion of generalists was stable over time with 14% generalists in 1990 and 17% in 2015. We did not find significant differences in estimates of optimal growth temperature between these populations (Supplementary Figure S5).

### Phenotypic responses to the coldest and the warmest temperatures exhibited the largest number of significantly associated genetic variants

To identify genetic determinants of local thermal adaptation, we conducted genome-wide association studies with each of the 12 phenotypic growth variables under each temperature condition (Supplementary Figures S10-S65). We found a total of 198 SNPs associated with at least one of the 12 growth variables that had significance values above the threshold for any of the temperature conditions (False Discovery Rate 10%; Supplementary Figures S10-S65; Supplementary Table S10). The 12°C temperature was associated with the highest number of significant variants (66% of the total), followed by 25°C (11%) (Figure 3A). The 22°C and 18°C temperatures had the lowest number of significantly associated variants (7% for both) (Figure 3A, Supplementary Table S10). The 198 significant variants overlap with 69 different genes, with a total of 38, 6, 3, 5, and 19 genes associated with 12°C, 15°C, 18°C, 22°C and 25°C, respectively (Supplementary Table S10). One gene (ZtIPO323_092450) was shared between the 15°C and 18°C conditions and another gene (ZtIPO323_038210) was shared between the 12°C and 25°C conditions. 36% of the genes have unknown functions and 64% have diverse predicted functions (Supplementary Table S10). A fraction (26%) of the associated genes have missense variants, including eight genes associated with 12°C, three genes with 15°C, one gene with 18°C and six genes with 25°C (Figure 3B; Supplementary Table S10).

To identify the most promising candidate genes for thermal adaptation, we compared our list of candidate genes with candidate genes identified in previous studies which used different approaches. One study used a QTL mapping approach with crosses between four Swiss strains (Lendenmann et al. (2016); hereafter “QTL”). Two other studies used a genome-environment association (GEA) approach, associating variants with the 11 temperature-associated bioclimatic variables and 8 additional bioclimatic variables deduced from the isolates’ geographical origin (Feurtey et al. 2023; Tralamazza et al. 2024; hereafter “GEA”). We identified 13 loci in common with previous QTL and GEA studies that included both global and local populations (Figure 3A) (Feurtey et al. 2023; Lendenmann et al. 2016; Tralamazza et al. 2024). Our GWAS identified four genes within the same QTL previously identified on chromosome 1. Three out of the four genes involved only a single significant SNP (Supplementary Table S10). The last gene (ZtIPO323_022050) codes for an ammonium transporter 1-like protein and was also found through GEA to be associated with one bioclimatic variable (isothermality) (Feurtey et al., 2023). ZtIPO323_022050 is impacted by seven significant SNPs associated with 12°C growth including one non-synonymous missense variant, while the others include three 5’ UTR variants and three synonymous variants (Supplementary Table S10, Figure 3C). The alternative allele of the missense variant is widely distributed except for strains from the arid climate (BSh) (Figure 3C), strongly suggesting an association with cold climate adaptation. Five other candidate genes for cold adaptation identified in our study were also found in common with the previous QTL study. Two of these genes were on chromosome 2, including ZtIPO323_027820 (protein Hook homolog1-like) and ZtIPO323_029900 (putative translation initiation factor). On chromosome 3, five significant SNPs associated with growth variables at 12°C were found in the ZtIPO323_044770 (unknown function), ZtIPO323_045330 (ketoacyl thiolase), and ZtIPO323_045710 (peptidase-like protein), that were also found in QTL for growth at lower temperatures (Figure 3A, Supplementary Table S10).

We also identified candidate genes for adaptation to warmer temperatures. We identified the gene ZtIPO323_054070 encoding the COMPASS/Set1C complex protein, involved in H3K4me3 methylation (Lukito et al. 2019), with five significant SNPs in our study and overlapping with a QTL for growth at 22°C (Figure 3A, Supplementary Table S10). The alternative allele of the most significant SNP is present in more than half of the genomes of strains from the arid climate (BSh) and it is the prevalent allele in Australia (Cfa, Csa), furthering its association with local adaptation to warmer growth conditions (Figure 3D). Additionally, we found overlaps between four variants significantly associated with growth at 25°C and four genes associated with the mean temperature of the driest quarter in one of the GEA studies (Feurtey et al., 2023). The first overlap includes the gene ZtIPO323_083310, which is described as a cell death-inducing p53 target 1 gene. The second overlap includes a cluster of three genes: ZtIPO323_084300, ZtIPO323_084310, and ZtIPO323_084320; described as a putative ATP-dependent helicase gene, an unknown function gene and a gene related to dienelactone hydrolase, respectively. The variant with the strongest association with growth at 25°C in this overlap is a non-synonymous missense variant (chromosome 7, position 1512102), co-located with the predicted dienelactone hydrolase (Supplementary Figure S9). The alternative allele for this variant is mostly distributed in Western and Eastern Australia and Chile (Figure 3D).

## Discussion

Adaptation to local temperature environments is one of the key challenges faced by plant pathogens as they expand into new niches. Here, we explored the genetic architecture of thermal adaptation to local environments in the widely distributed fungal pathogen *Z. tritici.* We compared *in vitro* growth variables under five temperature conditions for more than 400 global strains with bioclimatic variables of their associated sampling locations. We found strong evidence for local thermal adaptation, with a significant positive correlation observed between bioclimatic variables at the sampling sites and the optimal growth temperatures of the associated strains. The high variability in thermal performance found within global populations of *Z. tritici* is consistent with a high potential for this pathogen to adapt to any changes in local climate. Our GWAS across multiple thermal growth phenotypes reveals the complex polygenic nature of responses to global temperature variations. Several of the candidate genes revealed in the GWAS overlapped with loci from previous studies that used different methods and strain collections, reinforcing their potential significance as contributors to thermal adaptation. Among these, the GWAS-identified ammonium transporter-1-like protein emerged as a particularly strong candidate for future functional validation as a key contributor to thermal adaptation.

Correlations between the bioclimatic variables from the different sampling sites and the measured phenotypes support the existence of local thermal adaptation in *Z. tritici* populations around the world. In particular, we observed a positive significant correlation between the optimal growth temperature of the strains and the mean annual temperature of their sampling locations. This pattern aligns with findings from a previous study with a narrower geographical scope that reported a significant positive correlation between optimal growth temperature and average monthly temperatures (Boixel, Chelle, and Suffert 2022). Our analyses of thermal response in *Z. tritici* revealed distinct types of thermal adaptation. Specifically, we observed that strains from the Csb and Cfb climate classes had the lowest average optimal growth temperatures. These climates display summers with mean temperatures ranging from 15°C to 22°C (mean temperature of the warmest quarter), lower than the other temperate climates in our study, which have mean summer temperatures above 22°C. In addition, we observed some degree of specialization to either warmer or colder temperatures between populations. For warmer temperature specialization, we found that the Australian Wagga Wagga (Cfa climate) and Badgingarra populations (Csa climate) had more than 75% of strains performing better at warmer temperatures than at colder temperatures. We consider it likely that the founder bottlenecks and restricted gene flow into these populations contributed to the observed local thermal adaptation in these Australian populations, which remain largely isolated from other global populations. For colder temperature specialization, we found that over half of the strains in the populations from the Cfb climate were cold-specialized. These findings align with previous observations that populations from Oregon (Csb) and Switzerland (Cfb) appear to be more cold-adapted than those from Australia (Cfa) and Israel (BSh) (Zhan and McDonald 2011).

Individual *Z. tritici* field populations contained strains displaying a wide range of responses to temperature, including strains specialized to warmer or colder temperatures, as well as generalist strains. The coexistence of both specialists and generalists within the same population could be advantageous for ensuring the continued survival of *Z. tritici* in diverse wheat fields worldwide. The high variability in thermal growth response found within individual wheat fields from different climate classes reflects the high genetic variability found within *Z. tritici* field populations (McDonald et al. 2022). High phenotypic and genotypic variability should ensure the persistence of this pathogen in wheat fields globally, despite the predictions of increasingly severe temperature fluctuations (Cohen, Pfeiffer, and Francis 2018; Johnson et al. 2018). We found that the climates Dfb and BSh, which exhibit the highest seasonal temperature fluctuations, also contained the highest proportion of generalist strains. Thus, our results agree with earlier findings that thermal generalists are more common in more variable environments (Angilletta, Niewiarowski, and Navas 2002; Boixel, Chelle, and Suffert 2022; Huey and Hertz 1984; Kassen 2002; Pörtner and Farrell 2008; Ragland and Kingsolver 2008). The lack of a significant difference in optimal growth temperatures between pairs of populations sampled from the same location, 25 years apart, supports the resilience of *Z. tritici* populations. It is worth noting that local variations in temperature conditions during a single season (Boixel, Chelle, and Suffert 2022), in the wheat canopy (Chelle 2005), and in individual leaves (Bernard et al. 2013) can impact the pathogen. Thus, fluctuating temperatures at several scales can contribute to the maintenance of variability in thermal responses within field populations.

Our GWAS analyses revealed the complex genetic landscape of thermal adaptation in *Z. tritici*, with numerous SNP variants significantly associated with growth across five temperatures. These SNPs are located inside or near 69 putative genes, illustrating that thermal adaptation is a highly polygenic quantitative trait. The polygenic nature of temperature adaptation in *Z. tritici* was previously discovered using QTL mapping of growth at two temperatures in crosses between four Swiss strains (Lendenmann et al. 2016). Polygenic quantitative inheritance was also inferred in a GEA analysis of associations between 19 bioclimatic variables and a thousand-genome collection of *Z. tritici* that identified 187 genes co-located with significantly associated variants (Feurtey et al. 2023). Similar polygenic architectures have been observed for thermal response in other fungi including *S. cerevisiae* (Maclean et al. 2017; Weiss et al. 2018) and *Agaricus bisporus* (Foulongne-Oriol et al. 2014). However, our study represents the first comprehensive effort to investigate thermal adaptation in fungi using genome-wide association studies (GWAS) based on extensive phenotypic data collected from a global collection of more than 400 strains. Our findings suggest that GWAS and similar forward-genetics approaches are effective tools for identifying genetic regions associated with temperature adaptation in *Z. tritici*. We propose that similar approaches could be implemented for other fungi to better understand how they may adapt to climate change.

The identification of a putative ammonium transporter 1-like gene (ZtIPO323_022050) associated with adaptation to colder temperatures revealed a potential genetic mechanism underlying this pathogen’s adaptation to colder climates. This gene candidate was also identified by GEA studies with bioclimatic variables (Feurtey et al. 2023) and overlaps with a previously identified QTL region of 500 kb on chromosome 1 in a cross among Swiss strains (Lendenmann et al. 2016). We found that the alternative allele of the most significant non-synonymous variant for this gene is the predominant allele in the populations from most climates except the arid climate (BSh) found at the center of origin of *Z. tritici* (Stukenbrock et al. 2007). Adaptation to colder temperatures may have been a key attribute that enabled *Z. tritici* to become established in wheat production areas located in colder temperate climates. Class 1 ammonium transporters (AMT1) are mostly found in plants (McDonald, Dietrich, and Lutzoni 2012) but have not been characterized in fungi. However, class 2 ammonium transporters and methylammonium permeases have been identified in several fungal species (Berg, Lister, and Rutherford 2019; McDonald, Dietrich, and Lutzoni 2012). The roles of these transporters in ammonium sensing and transport are not yet well understood. Some ammonium transporters are involved in the yeast-hyphae morphological switch in species such as *S. cerevisiae* (Lorenz and Heitman 1998), *C. albicans* (Biswas and Morschhäuser 2005), *Ustilago maydis* (Smith et al. 2003), and *Cryptococcus neoformans* (Rutherford et al. 2008). Functional studies of ammonium transporters in plant pathogenic fungi show a reduced virulence or secondary metabolite production in deletion mutants, as observed in *Colletotrichum gloeosporioides* (Shnaiderman et al. 2013), *Fusarium fujikuroi* (Teichert et al. 2008), and *U. maydis* (Paul et al. 2018). The functions and effects on virulence of ammonium transporters in *Z. tritici* remain unknown. However, finding a gene with potential roles in thermal adaptation, yeast/hyphae dimorphism, and virulence is consistent with earlier findings in *Z. tritici* showing that quantitative trait loci can have pleiotropic effects. Pleiotropy in life history traits of *Z. tritici* has been widely reported (Dutta et al. 2021; Francisco, McDonald, and Palma-Guerrero 2023; Lendenmann et al. 2016; Lendenmann, Croll, and McDonald 2015; Lendenmann et al. 2014). Crucially, the relationship between thermal adaptation and virulence is likely to be complex, as growth rates at different temperatures and virulence can show synergistic or antagonistic pleiotropy depending on the host cultivar (Dutta et al. 2021).

## Materials and methods

### Sample collection

We used a worldwide collection of 420 *Z. tritici* strains sampled from wheat fields in 20 countries across different wheat-growing regions between 1981 and 2016. The collection encompasses 38 geographical populations and includes 28 field populations (strains sampled on the same day from the same naturally infected wheat field) from 15 countries (402 strains). The other 18 strains came from Algeria (3), Australia (2), Ethiopia (1), Syria (3), Turkey (5), Kenya (2), and Yemen (2). The field populations were located in six different climate zones. The Middle Eastern populations (52 strains) originated from an arid climate (Köppen-Geiger climate classification: B). The population from Canada (14 strains) originated from a continental climate (Köppen-Geiger: D). The remaining 336 strains originated from temperate climates in Northern Africa, Europe, the Americas, and Australia (Köppen-Geiger: C). However, these temperate climates differed according to precipitation pattern and heat intensity during the summer. The populations from Europe and Argentina (115 strains) originate from a temperate climate with warm and humid summers (Köppen-Geiger: Cfb). The Oregon (USA) populations and Chile (90 strains) originated from a temperate climate with warm and dry summers (Köppen-Geiger: Csb). The populations from Uruguay, East Australia, and most North American locations (90 strains) originated from a temperate climate with hot and humid summers (Köppen-Geiger: Cfa). The North African, Californian (USA), and Western Australian populations (41 strains) originated from a temperate climate with hot and dry summers (Köppen-Geiger: Csa). Detailed information for each strain can be found in Supplementary Table S1. The isolates were preserved in 50% glycerol and anhydrous silica at −80°C.

### Genomic datasets, sequencing, and variant calling

Publicly available Illumina whole genome sequences of 298 strains were retrieved (see Supplementary Table S1 for accession information) and added to newly generated sequencing data for 122 strains. We screened the 122 strains for clones using microsatellite markers as described in Lorrain et al. (2024) and no clones were identified. DNA extractions were performed and sequenced using Illumina paired-end sequencing with a read length of 150 bp (Novogene Europe, Germany), as described in Lorrain et al. (2024). The genomic data generated during this study are available from the NCBI Sequence Read Archive platform under the BioProject PRJNA1153813.

We performed single nucleotide polymorphism (SNP) and short indels variant calling by mapping onto the genome of the reference strain IPO323 (Goodwin et al. 2011). We first used Trimmomatic v.0.35 (Bolger, Lohse, and Usadel 2014) to remove adapters, low-quality bases (<28), and sequences shorter than 50 bp. We aligned the trimmed sequences to the reference genome using the software BWA v.0.7.17 (H. Li and Durbin 2010) and removed the duplicated sequences with the “MarkDuplicates” option of Picard v.2.25.7 (https://broadinstitute.github.io/picard/). For the short variant calling, we used the functions “HaplotypeCaller”, “CombineGVCFs”, and “GenotypeGVCFs” from GATK v.4.1.2.0 (Auwera and O’Connor 2020), setting the ploidy to one and maximum alternative alleles to three as described in Feurtey et al. (2023). To filter out erroneous variant callings, we used the “VariantFiltration” option with a per-genotype removal of depth lower than three and with the per-site filters FS > 10, MQ < 20, QD < 20, ReadPosRankSum - 2:2, MQRankSum −2:2, and BaseQRankSum −2:2. We assessed the individual levels of missing data of the strains through the option –missing-indv vcftools v.0.1.14 (Danecek et al. 2011) and removed four strains that had more than 80% missing data (Supplementary Table S1). We also filtered out positions with a calling-rate below 20% of the isolates with the –max-missing option of vcftools v.0.1.14 and filtered for a minor allele frequency of 5% (MAF > 0.05). After filtering, we retained 416 genomes from our genomic dataset.

### Population structure, genetic diversity, and differentiation

We assessed the population structure on a subset of biallelic SNPs pruned to remove physically linked variants (1 SNP per kb) using a principal component analysis (PCA) with PLINK 2.0 (Chang et al. 2015) and estimated individual admixture coefficients with the snmf method in the R package LEA (Frichot and François 2015). We considered K values between one and 15 with 10 repetitions for the genetic admixture analysis (Supplementary Figure S1). We assessed the most appropriate K value, based on the model fit to the data through the entropy method from snmf and the number of strains assigned to any cluster with a coefficient higher than 0.75 (non-admixed, Supplementary Figure S2B-D). After admixture analysis we filtered out five strains that did not correspond to the genetic cluster assigned to the rest of their corresponding populations, likely a result of mislabelling (Supplementary Figure S2A, Supplementary Table S1). The final dataset included a total of 411 genome sequences to use in further analyses.

To explore the differentiation between and within genetic groups we used an Analysis of Molecular Variance (AMOVA) implemented in the R package *poppr* and calculated the Weir and Cockerham fixation index (F_st_) between all pairs of genetic clusters with the vcftools v.0.1.17 function--weir-fst-pop with--haploid mode (https://github.com/jydu/vcftools). We calculated whole genome nucleotide diversity (Nei and Li’s *π*) for each genetic cluster, within 10 kb windows using the--window-pi function of vcftools v.0.1.17.

### *In vitro* phenotyping of thermal performance

Thermal performance of the 411 strains was measured using a modification of the approach described by Boixel et al. (2019). We pre-cultured each strain by adding 150 µL of its glycerol stock into 50 mL of yeast extract peptone dextrose media (YPD: yeast extract 10 g/L, bactopeptone 20 g/L, dextrose 20 g/L, kanamycin 50 mg/L) and incubated at 18°C for seven days in the dark at 120 rpm. The seven-day-old cultures were filtered through two layers of gauze to enrich blastospores (yeast-like asexual spores) and then centrifuged at 3273 g for 10 min at 4°C. The spores were re-suspended in approximately 20 mL of glucose peptone liquid media (GPL: dextrose 14.3 g/L, bactopeptone 7.1 g/L, yeast extract 1.4 g/L, kanamycin 50 mg/L). This spore solution was thoroughly mixed and diluted 1:10 and 1:100 for spore counting. The spore dilutions were loaded into a hemocytometer (KOVA® GLASSTIC® slide) and counted using a microscope-mounted camera and an ImageJ macro (https://github.com/jalassim/SporeCounter; Spore Counting v.9). We then adjusted the spore suspensions of each strain into 150 µL of GPL into four replicate wells of a sterile polystyrene 96-well microtiter plate with a flat-bottom (Greiner Bio-One, item num. 655161) to achieve a starting spore concentration of 2.5×10^5^ spores/mL. Sixteen wells per plate were filled with only GPL media that we used as absorbance blanks (control). The microtiter plates were sealed with breathable tape to enable gas exchange (Breathe-Easy®, Diversified Biotech). Duplicate plates were incubated in the dark for 144 hours at 70% humidity at five different temperatures: 12°C, 15°C, 18°C, 22°C and 25°C.

Fungal growth was measured by assessing the optical density at 405 nm (OD_405nm_) (Boixel et al. 2019) twice per day with a plate reader (Spark™ 10M, Tecan). We performed measurements using three technical replicates for each plate to account for instrument error. To convert optical density measures into spore concentrations, we produced calibration curves for each strain with a dilution series of 1×10^5^, 2.5×10^5^, 5×10^5^, 7.5×10^5^, 1×10^6^, 1.5×10^6^, 1.75×10^6^ and 2×10^6^ spores/mL. We used the *lm* function of the *stats* R package to calculate a linear regression of the spore concentration and optical density per strain (Supplementary Table S2).

To test for contamination and errors in the wells containing only GPL media, we applied a standard deviation-based filtering at each time point (Supplementary Table S2). To ensure accuracy and eliminate potential errors in the spore suspension measurements, we implemented two consecutive outlier filtering steps. First, we applied an upper-bound interquartile range (IQR) filter to the measurements at each timepoint, temperature, and replicate. This initial step removed 6,853 outliers from 308,416 measurements (see Supplementary Table S2). Next, we applied a second upper-bound IQR filter to the median concentration of each strain at each timepoint and condition, which resulted in the removal of 444 additional outliers from 301,563 measurements (Supplementary Table S2). No more than two replicates out of four per strain were removed, and we did not observe any bias related to specific populations or microtiter plates in the filtered measurements (Supplementary Table S2).

### Thermal response growth curve variables

We calculated the median spore concentration per strain, timepoint, and temperature to generate individual growth curves. We extracted 12 growth parameters from these curves to investigate various aspects of the *Z. tritici* thermal response (Supplementary Figure S4; Supplementary Table S3). We used logistic regressions to estimate the growth rate, the inflection time, and the asymptote of each curve using the *nls* function in R with a self-starting logistic model. We calculated the empirical area under the curve (eAUC) for each growth curve by averaging the right and left Riemann sums, summarizing the overall growth pattern of each strain in each temperature condition. Additionally, we used locally weighted polynomial regressions with the R function *loess* of each curve to estimate the time to the first, second, and third quantiles of the concentration of the curve. This allowed us to extract time measurements related to different moments of the growth curve that do not depend on the successful fit of a logistic model. With these regressions, we also estimated the spore concentrations for each curve every 48 hours, with concentrations estimated at 48 hours, 96 hours and 144 hours. All the R functions employed in this analysis belong to the built-in *stats* package.

We used a quadratic regression on the eAUC values across the five temperature conditions tested (12°C, 15°C, 18°C, 22°C, and 25°C) to estimate the optimal growth temperature for each strain. We excluded 50 strains that showed an optimal temperature of either 12°C or 25°C, as these could produce artifacts due to boundary effects. After determining the optimal growth temperature for each strain, we assessed the temperature sensitivity by dividing the eAUC at each tested temperature by the eAUC at the strain’s optimal temperature. We identified and removed outliers for all growth variables using a Local Outlier Factor (LOF) filter with 20 nearest neighbors, implemented with the *lof* function from the R package *dbscan* (Hahsler, Piekenbrock, and Doran 2019).

To summarize, we extracted i) four variables representing a general summary of growth: the eAUC, the optimal temperature, the temperature sensitivity, and the growth rate (Figure 2C, Supplementary Figures S6A, D, K); and ii) eight variables representing static points in the growth curve: the inflection time, the asymptote, the first, second, and third quantiles of the concentration of the curve, and the spore concentration after 48, 96 and 144 hours (Supplementary Figures S6B, C, E, F, G, H, I, J). An explanatory diagram of the different growth curve-derived variables can be found in Supplementary Figure S4. The estimates per growth variable for each strain can be found in Supplementary Table S3 and the summary statistics per population and climate in Supplementary Tables S4 and S5, respectively.

### Comparing local climatic conditions with thermal growth patterns

From the metadata of each strain, we obtained the approximate geographical coordinates for each population’s location (Supplementary Table S1). We retrieved 11 bioclimatic variables related to temperature values for each population’s coordinates from the database WorldClim2 at a 10-minute resolution mean from 1970 to 2000 (Fick & Hijmans, 2017). These bioclimatic variables consider different aspects of the temperature-related climatic variables in a location for different temporal scales, for example: the annual mean temperature, the temperature annual range, and the mean temperature of the driest quarter. The values of these bioclimatic variables can be found in Supplementary Table S6 and are represented in Figure 2A (5 variables) and Supplementary Figure S6 (the other 6 variables). We estimated Pearson’s correlation between these bioclimatic variables and the 12 growth variables with the *cor* function in the R built-in package *stats*. The Pearson’s correlation coefficients and the *p*-values are presented in Supplementary Tables S7 and S8, respectively.

To assess differences in temperature response between populations, we performed Kruskal-Wallis and pairwise Wilcoxon signed-rank tests of the 12 growth variables. The Kruskal-Wallis tests were performed using the estimates for each growth variable, except when the number of observations per population was less than five. Because there are differences in the number of growth variable estimates per population and climate, we performed additional Kruskal-Wallis tests by subsampling randomly 10 times across all populations based on the minimum number of strains in the comparison population and then calculated the mean *p*-value between the 10 subsampled tests. We employed the function *kruskal.test* from the R built-in package *stats* for the Kruskal-Wallis tests. Pairwise comparisons between populations were made using Wilcoxon signed-rank tests with a Bonferroni correction using the function *pairwise_wilcox_test* from the R package *rstatix*. We used the same procedure to assess differences in the temperature response between climate classes.

In addition, we evaluated differences between the variance in growth variable estimates from different populations and climate classes by performing Levene’s tests. We employed the function *leveneTest* from the R package *car*. As for the Kruskal-Wallis tests, we performed additional Levene’s tests in subsamples accounting for differences in sample sizes of each population/climate comparison.

### Thermal specialization profiles

To compare strain performance in cold temperatures versus warm temperatures, we classified the strains into seven categories. The scale was created as follows: from the distribution of strain temperature sensitivity at 15°C and 25°C, we calculated the quartile (0-4) of temperature sensitivity for each strain. We selected the temperature sensitivities at 15°C and 25°C as they produced comparable growth reduction patterns in the phenotyped strains (Supplementary Figure S7). Each strain was ranked as belonging to a specific quartile value for 15°C and to a specific quartile value for 25°C, with a value of 0 indicating that a strain is among the most sensitive strains at this temperature and a value of 4 assigned to the least sensitive strains. For each strain, a general index of temperature performance was estimated by subtracting the quartile value at 15°C from the quartile value at 25°C. The resulting performance scale was as follows: −3, growth much better at cold temperatures; −2, better at cold; −1, slightly better at cold; 0, similar at cold and warm; 1, slightly better at warm; 2, better at warm; 3, much better at warm. A diagram explaining this calculation can be found in Supplementary Figure S8 and the estimates of these calculations can be found in Supplementary Table S9.

### Genome-wide association studies

To identify genomic regions associated with temperature adaptation, we used all growth phenotypes to perform 56 GWAS with the GEMMA-based pipeline *vcf2gwas* (Vogt, Shirsekar, and Weigel 2022; Zhou and Stephens 2012). To control for population structure, the pipeline calculates a relatedness matrix by default, and we used the option-cf PCA-c 2 which uses the first two principal components of the PCA of all the variants. We used the option −lmm to perform association tests with univariate linear models. We set two significance thresholds with different levels of stringency: Bonferroni threshold (⍺=0.05), and false discovery rate (FDR) at 10% obtained with the Bioconductor package *qvalue* (Storey et al. 2023). To better understand the effects of significantly associated variants, we used SNPeff (Cingolani et al. 2012) with an upstream/downstream interval length of 2800 bp (mean inter-genic distance of *Z. tritici* from the most recent gene annotation (Lapalu et al. 2023) is 2858 bp). This allowed us to distinguish among variants with strong effects (e.g., frameshift variants), variants with small effects (e.g., synonymous mutations), and variants found outside of coding sequences but potentially involved in regulating gene expression as described in Feurtey et al. (2023). The significant associated variables can be found in Supplementary Table S10.

### Artificial intelligence disclosure

As we prepared this manuscript, the authors used CHATGPT to improve the wording and clarity of the draft in the abstract, introduction, and discussion sections. The authors reviewed the output of the tool and edited it accordingly. The authors take full responsibility for the content of the publication.

## Data availability

The sequencing data is publicly available from the NCBI Sequence Read Archive. Accession numbers can be retrieved from Supplementary Table S1. The newly sequenced data is under the BioProject PRJNA1153813. The bioclimatic variables for the location of the populations (coordinates in Supplementary Table S1) were obtained from the publicly available WorldClim database version 2 (Fick & Hijmans, 2017). The raw phenotypic data is publicly available in Zenodo (doi).

## Code availability

To ensure the reproducibility of the analyses presented in this manuscript, all custom scripts are available at https://github.com/SilviaMinana/Temperature_Adaptation_Z.tritici (https://doi.org/10.5281/zenodo.13750859). Data post-processing and visualization were done using R, bash, and Python, with the corresponding R Markdown reports available in the GitHub repository.

## Acknowledgments

We thank ETH Zurich student Katja Mühlecker for assistance with preparing phenotyping materials and each strain’s initial inoculum. We thank ETH Zurich student Daniel Osoko for preparing the preliminary R scripts used to process the phenotype data. Genotyping and growth phenotyping data were generated in collaboration with the Genetic Diversity Centre (GDC), ETH Zurich. This project was funded by ETH Zurich.

## Authors contributions

SMP, AF, CL, and BAM conceptualized the study. SMP, AF, and CL designed the experiments. SMP performed data acquisition. SMP and AF performed data analyses. SMP, AF, and CL contributed to writing the manuscript. BAM contributed to revising the manuscript.

---

**Supplementary Figure S1.** Admixture plots showing K values between 2 and 15 for the genetic clustering of 416 *Zymoseptoria tritici* strains, showing per-isolate bar plot cluster assignments. Each vertical bar represents a strain, and the clusters are represented by different colors. Strains are grouped by continent and country of origin.

**Supplementary Figure S2.** Genetic clusters of 416 *Z. tritici* strains. **A**) Admixture plot with K=8 that shows with arrows the strains that were incorrectly assigned to a cluster due to mislabeling. **B**) Cross-entropy for different numbers of clusters (K). **C**) Number of strains that were assigned to a cluster by the number of clusters (K). **D**) The number of strains in the smallest cluster according to K. **E**) The average ancestry coefficient per population assigned to a genetic cluster with K=8. Only populations with at least five strains are shown. **F**) Number of strains per population (out of 411 strains total) that belong to each genetic cluster (admixture coefficient > 0.75). The colors represent the eight genetic clusters, and the grey color represents the strains that did not have a high admixture coefficient for any of the clusters. The colors correspond to the colors of the genetic clusters shown in Figure 1.

**Supplementary Figure S3.** Values of six bioclimatic variables associated with temperature for each sampling location per climate class. These values are the averages for the years 1970-2000. **A**) Mean diurnal range, which is the mean of the mean monthly maximum temperature minus the mean monthly minimum temperature. **B**) Isothermality, which is the mean day-tonight temperature oscillation relative to the annual oscillations. **C**) Seasonality, which is the annual mean ratio of the standard deviation of the monthly mean temperatures to the mean monthly temperature. **D**) Maximum temperature of the warmest month. **E**) Minimum temperature of the coldest month. **F**) Temperature annual range, which is the difference between the minimum temperature of the coldest month and the maximum temperature of the warmest month.

**Supplementary Figure S4.** Diagrams showing how each of the 12 growth variables was derived.

**Supplementary Figure S5.** Estimates of the optimal temperature per population. Each point represents the optimal temperature for a strain. The significance levels of the Wilcoxon signed-rank tests with Bonferroni correction are represented by asterisks: * for *p*-value < 0.05, ** for *p*-value < 0.01, and *** for *p*-value < 0.001. Only the populations with more than five strains were represented.

**Supplementary Figure S6.** Estimates of the logistic growth rate (**A**), inflection time (**B**), asymptote (**C**), empirical area under the curve (**D**), concentration after 48 hours (**E**), concentration after 96 hours (**F**), concentration after 144 hours (**G**), time to 1^st^ quantile of the concentration (**H**), time to 2^nd^ quantile of the concentration (**I**), time to 3^rd^ quantile of the concentration (**J**), and the temperature sensitivity (**K**) for each climate class. Each point represents the value for each strain. The brown dots in the middle of the boxplots represent the mean values. The significance levels of the Wilcoxon signed-rank tests with Bonferroni correction are represented by asterisks: * for *p*-value < 0.05, ** for *p*-value < 0.01, and *** for *p*-value < 0.001. Only the climates with more than five strains were represented.

**Supplementary Figure S7.** Distribution of temperature sensitivity estimates per temperature condition tested.

**Supplementary Figure S8.** Diagram showing how calculations were made for the comparative temperature performance for each strain. “Q” represents the quartile and “n” represents the quartile number (0-4).

**Supplementary Figure S9**. Percentage of strains per population that perform better at warmer temperatures or at colder temperatures based on the comparison of the temperature sensitivity at 15°C and the temperature sensitivity at 25°C. Only populations with more than five strains are shown.

**Supplementary Figures S10-S66.** Manhattan plots of the genotype-phenotype associations for the 12 growth variables per temperature condition. Values obtained from the pipeline *vcf2gwas* using the software GEMMA with the default linear model. Darker colors were applied to dots if the variant was significant (FDR 10% threshold). The dotted horizontal lines indicate the Bonferroni thresholds.

**Supplementary Table S1.** Information for each strain, including the strain name, possible alternative names used in other publications, if it was previously publicly available (published), geographical location of the sampling site and inferred coordinates, sampling year, as well as the BioProject corresponding to the sequencing data.

**Supplementary Table S2.** Raw absorbance data with all the associated metadata and filtering status (kept or not for the analyses in the manuscript).

**Supplementary Table S3.** Estimates for all the growth variables for all the temperature conditions tested per strain.

**Supplementary Table S4.** Summary statistics of the 12 growth variables for all the temperature conditions per population.

**Supplementary Table S5.** Summary statistics of the 12 growth variables for all the temperature conditions per climate class.

**Supplementary Table S6.** The 11 bioclimatic variables estimated from the approximate coordinates per sampling location.

**Supplementary Table S7.** Pearson’s correlation coefficients of the 12 growth variables for all the temperature conditions and the 11 bioclimatic variables.

**Supplementary Table S8.** Pearson’s correlation significance *p*-value of the 12 growth variables for all the temperature conditions and the 11 bioclimatic variables.

**Supplementary Table S9.** Estimates of the comparative temperature performance derived from the temperature sensitivities at 15°C and 25°C per strain with population and climate information.

**Supplementary Table S10.** Description of the genetic variants significantly associated with the 12 growth variables per temperature condition tested, the allele distribution of these significant variants, the genes co-located with these variants, and the predicted effects of these variants on these genes as estimated using SnpEff.

**Supplementary File S1**: Detailed description of the supplementary tables.

