## Supplementary Figure for "Thermal adaptation in worldwide collections of a major fungal pathogen"

Silvia Miñana-Posada et al.

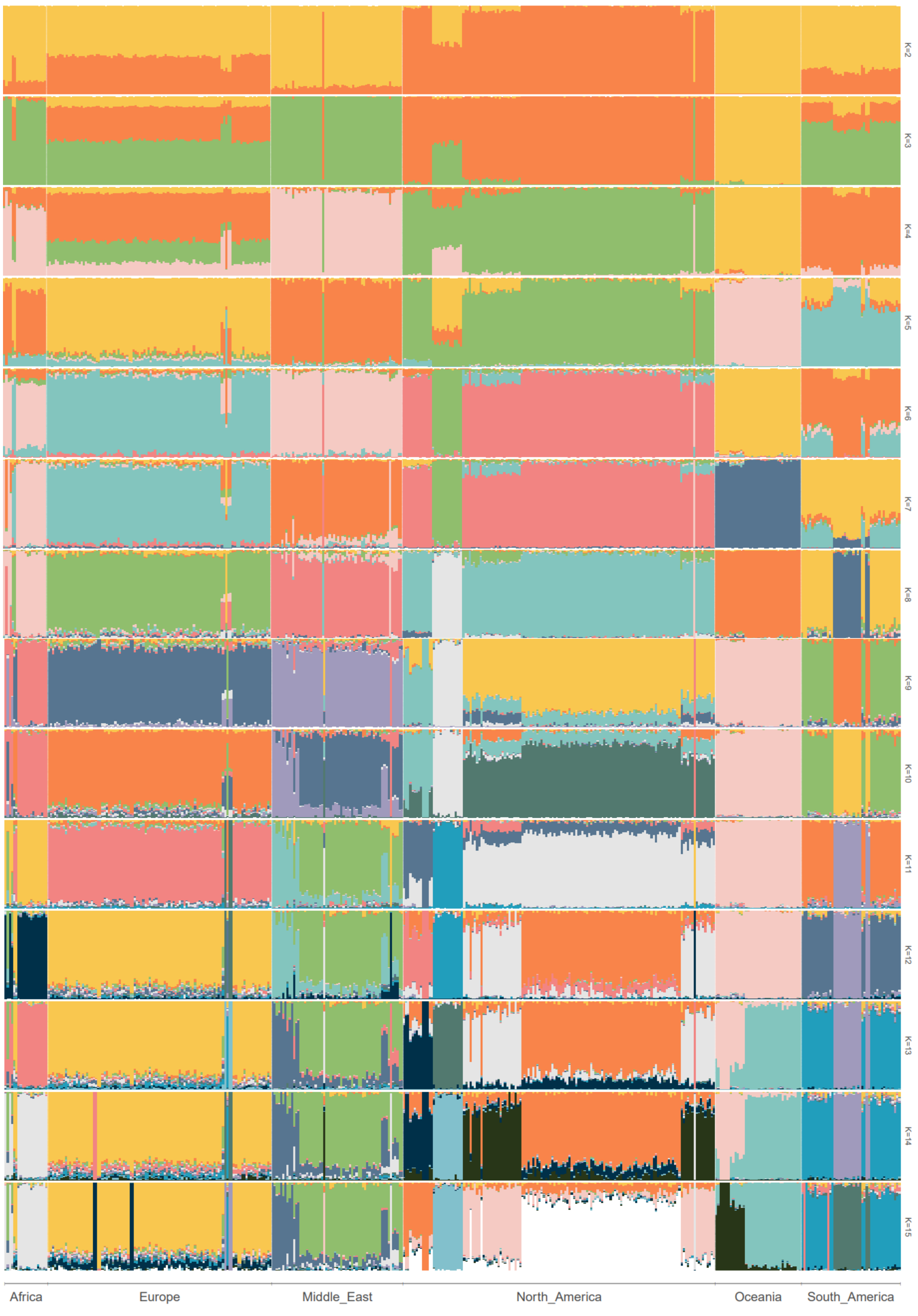


**Supplementary Figure S1.** Admixture plots showing K values between 2 and 15 for the genetic clustering of 416 *Zymoseptoria tritici* strains, showing per-isolate bar plot cluster assignments. Each vertical bar represents a strain, and the clusters are represented by different colors. Strains are grouped by continent and country of origin.

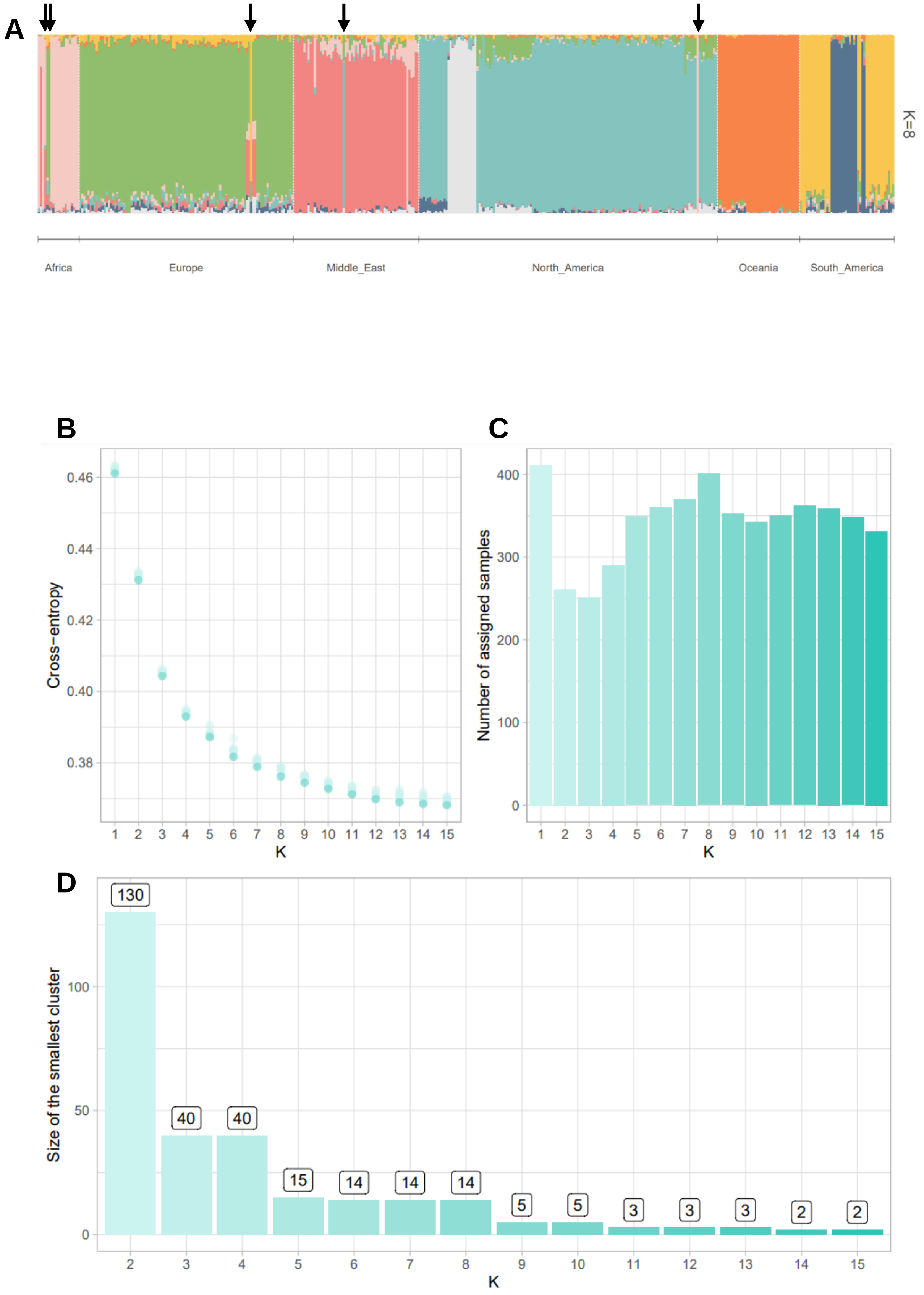
A
K=8
Africa
Europe
Middle_East
North_America
Oceania
South_America
B
Cross−entropy
0.46
0.44
0.42
0.40
0.38
1 2 3 4 5 6 7 8 9 10 11 12 13 14 15
K
C
Number of assigned samples
400
300
200
100
0
1 2 3 4 5 6 7 8 9 10 11 12 13 14 15
K
D
Size of the smallest cluster
100
50
0
130
40
40
15
14
14
14
5
5
3
3
3
2
2
2 3 4 5 6 7 8 9 10 11 12 13 14 15
K

**E**

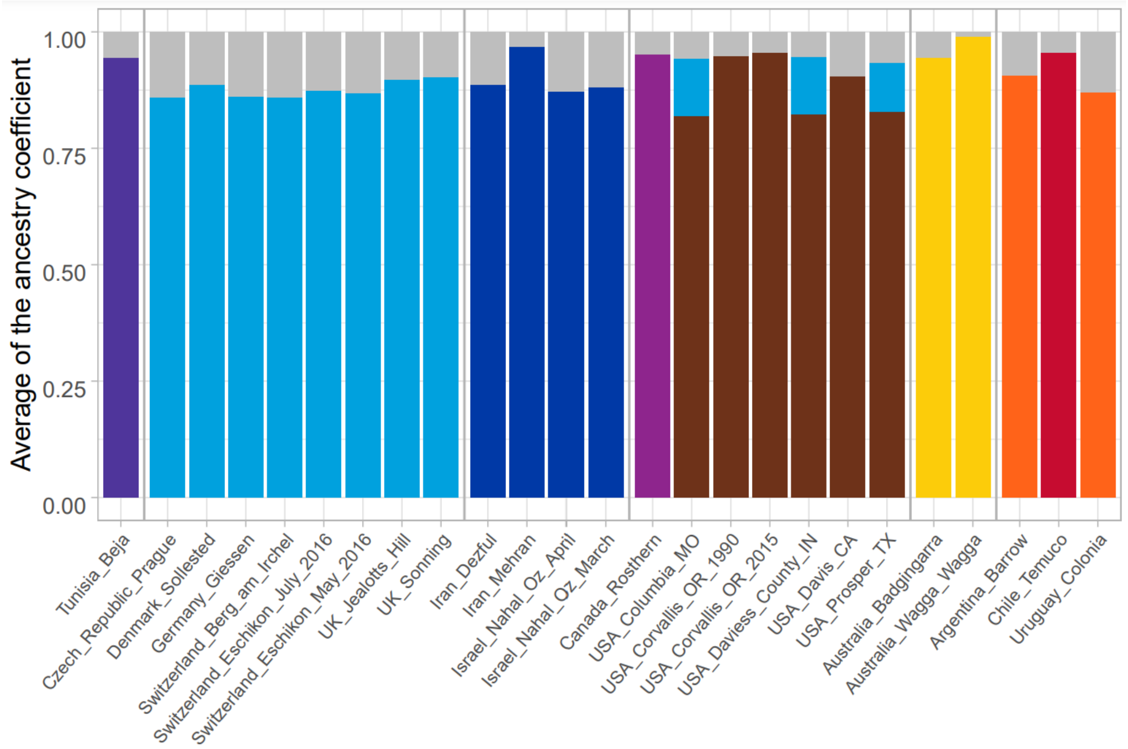


**F**

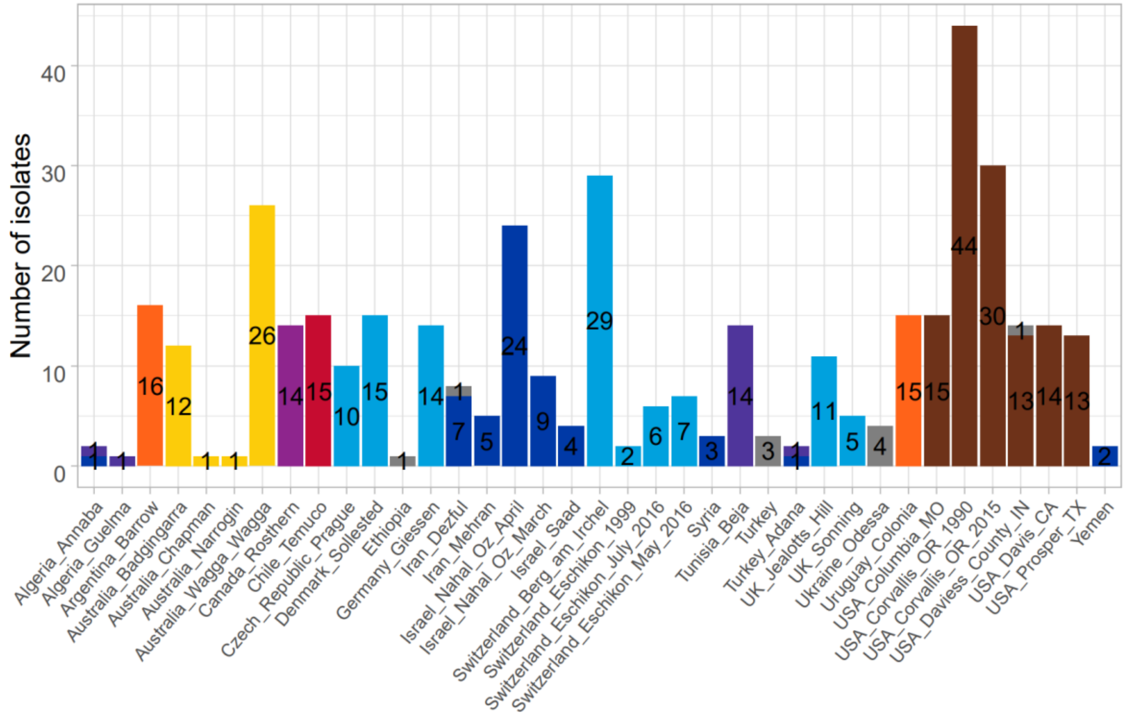


**Supplementary Figure S2.** Genetic clusters of 416 *Z. tritici* strains. **A**) Admixture plot with K=8 that shows with arrows the strains that were incorrectly assigned to a cluster due to mislabeling. **B**) Cross-entropy for different numbers of clusters (K). **C**) Number of strains that were assigned to a cluster by the number of clusters (K). **D**) The number of strains in the smallest cluster according to K. **E**) The average ancestry coefficient per population assigned to a genetic cluster with K=8. Only populations with at least five strains are shown. **F**) Number of strains per population (out of 411 strains total) that belong to each genetic cluster (admixture coefficient > 0.75). The colors represent the eight genetic clusters, and the grey color represents the strains that did not have a high admixture coefficient for any of the clusters. The colors correspond to the colors of the genetic clusters shown in Figure 1.

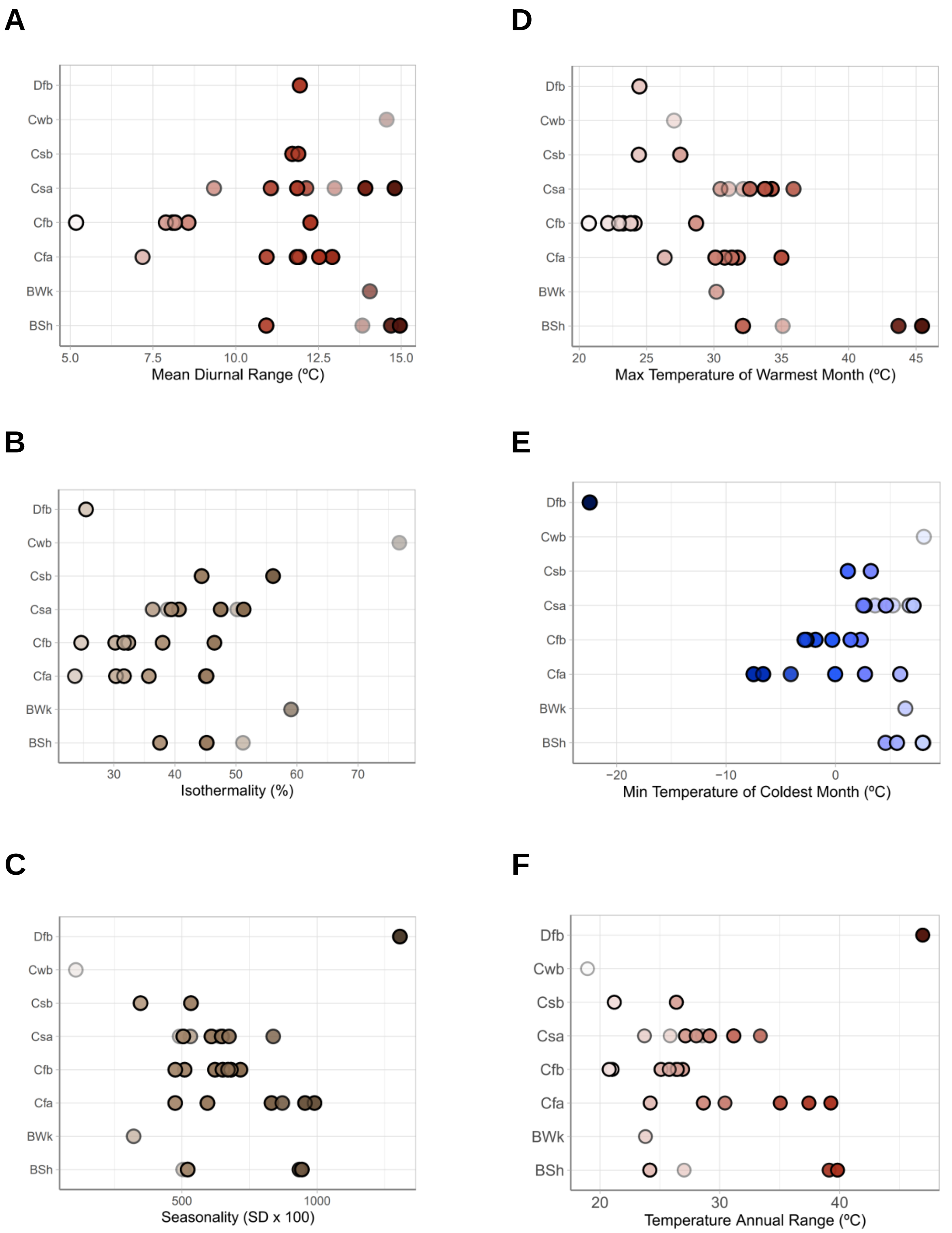


**Supplementary Figure S3.** Values of six bioclimatic variables associated with temperature for each sampling location per climate class. These values are the averages for the years 1970-2000. **A**) Mean diurnal range, which is the mean of the mean monthly maximum temperature minus the mean monthly minimum temperature. **B**) Isothermality, which is the mean day-tonight temperature oscillation relative to the annual oscillations. **C**) Seasonality, which is the annual mean ratio of the standard deviation of the monthly mean temperatures to the mean monthly temperature. **D**) Maximum temperature of the warmest month. **E**) Minimum temperature of the coldest month. **F**) Temperature annual range, which is the difference between the minimum temperature of the coldest month and the maximum temperature of the warmest month.

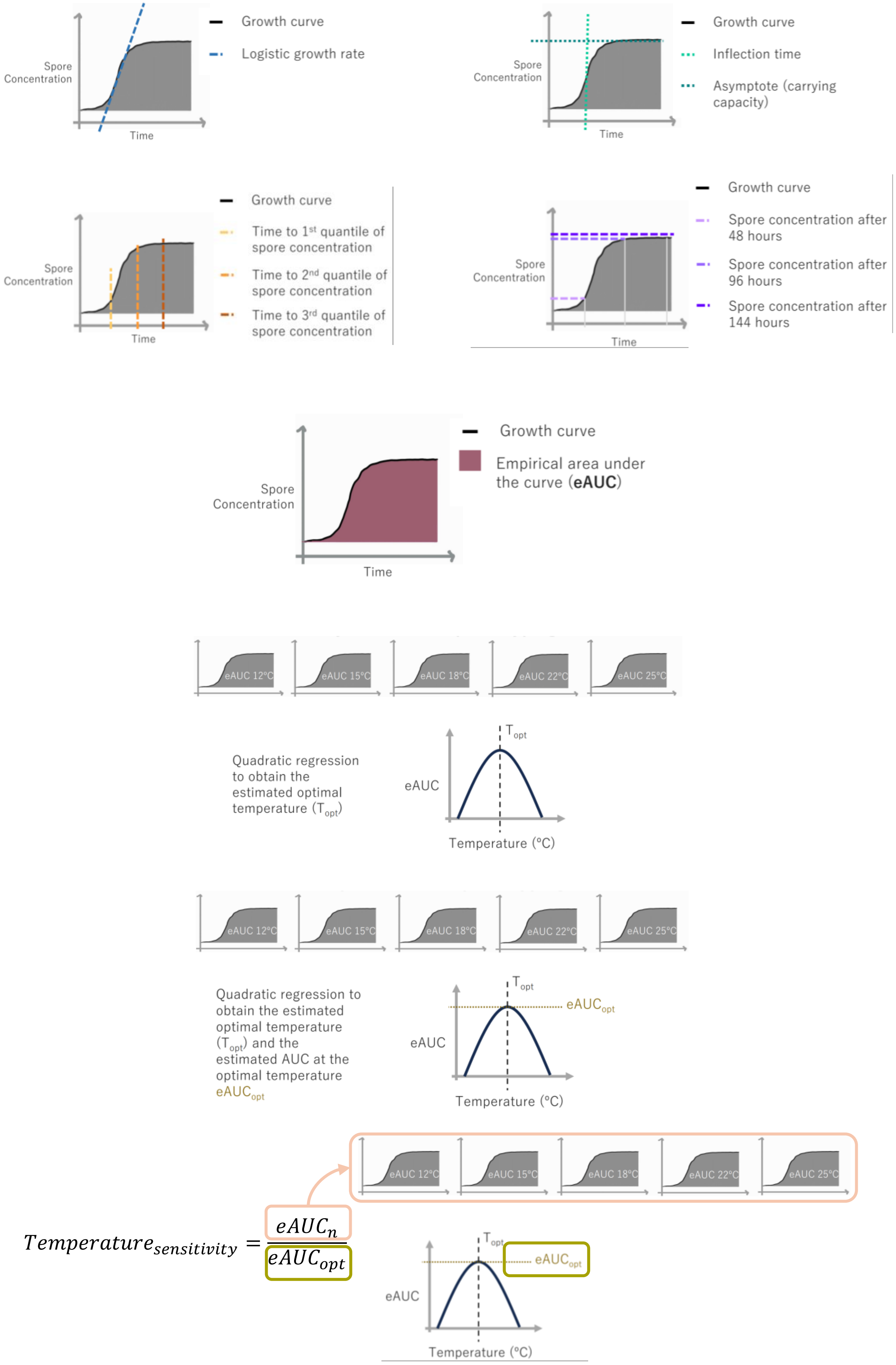


**Supplementary Figure S4.** Diagrams showing how each of the 12 growth variables was derived.

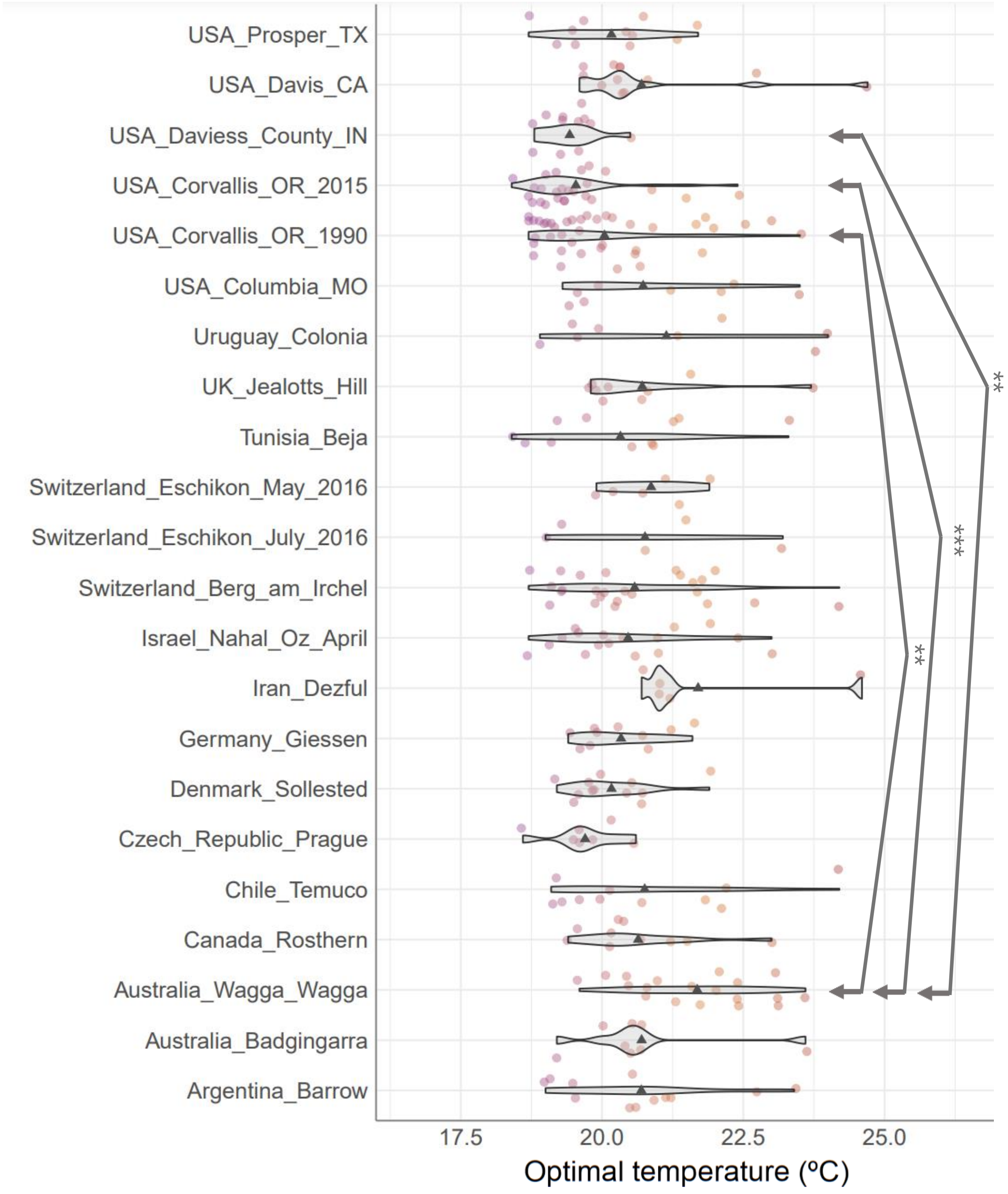


**Supplementary Figure S5.** Estimates of the optimal temperature per population. Each point represents the optimal temperature for a strain. The significance levels of the Wilcoxon signed-rank tests with Bonferroni correction are represented by asterisks: * for *p*-value < 0.05, ** for *p*-value < 0.01, and *** for *p*-value < 0.001. Only the populations with more than five strains were represented.

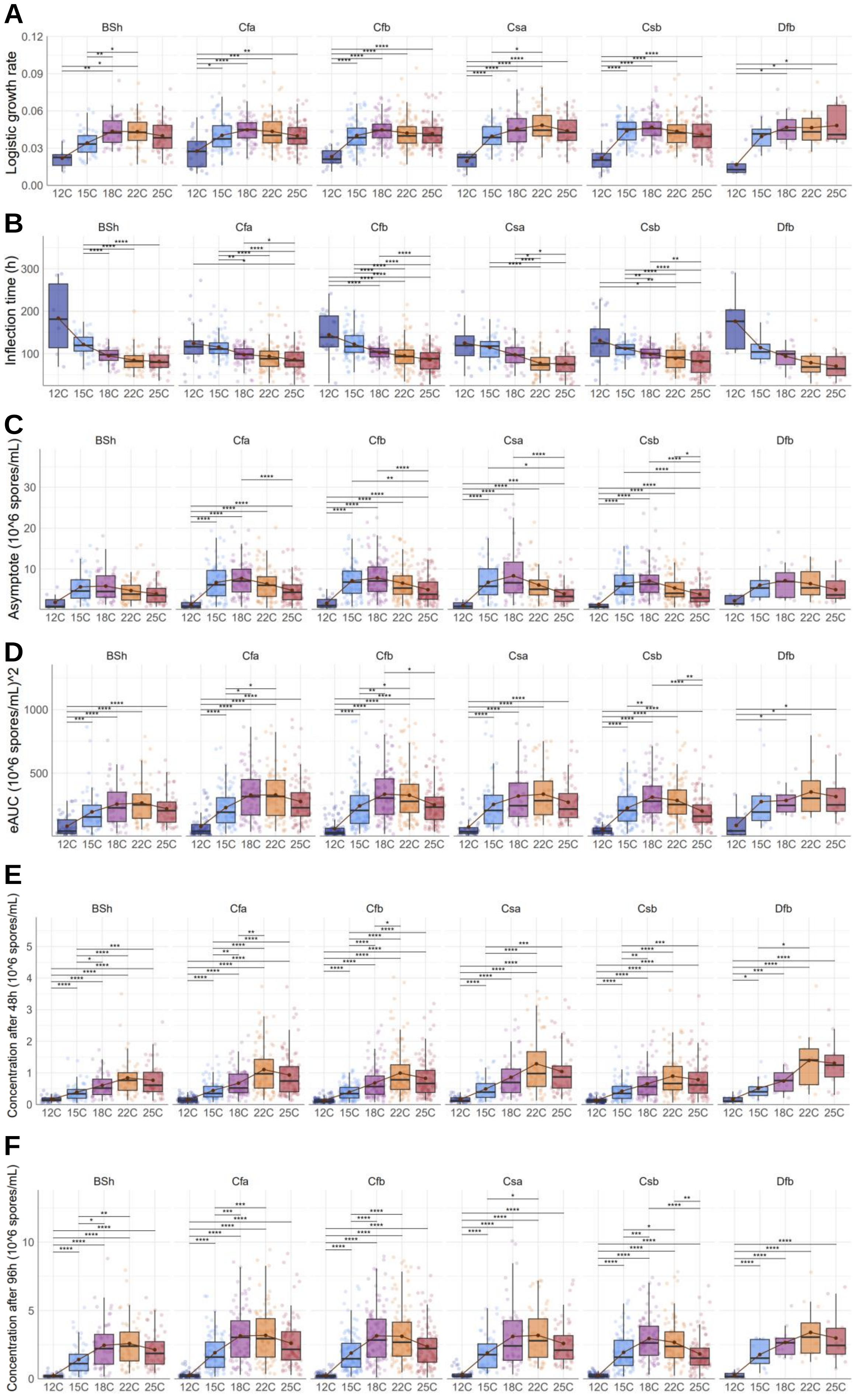
A
B
C
D
E
F
BSh
Cfa
Cfb
Csa
Csb
Dfb
Logistic growth rate
Inflection time (h)
Asymptote (10^6 spores/mL)
eAUC (10^6 spores/mL)^2
Concentration after 48h (10^6 spores/mL)
Concentration after 96h (10^6 spores/mL)
12C
15C
18C
22C
25C

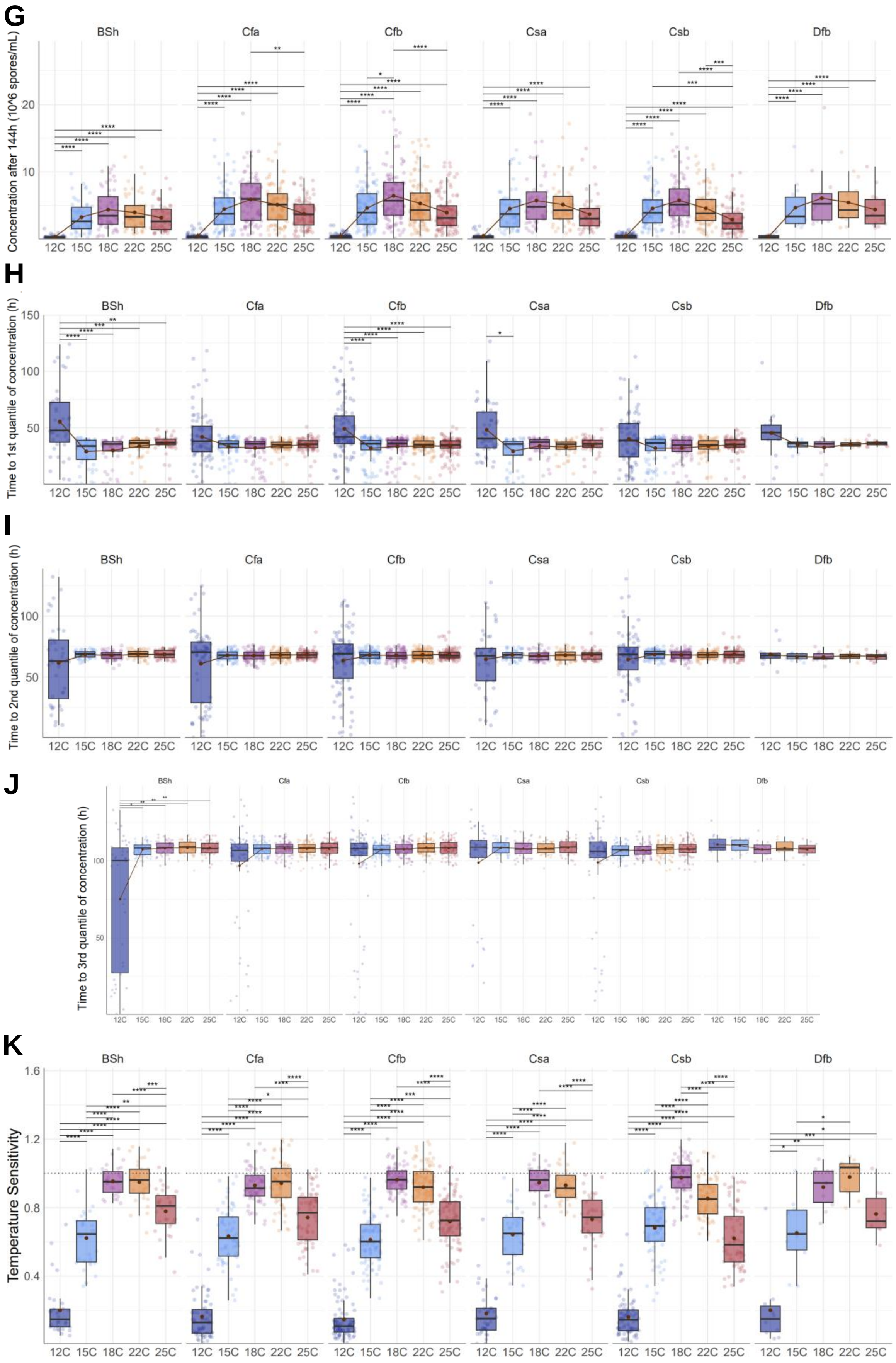


**Supplementary Figure S6.** Estimates of the logistic growth rate (**A**), inflection time (**B**), asymptote (**C**), empirical area under the curve (**D**), concentration after 48 hours (**E**), concentration after 96 hours (**F**), concentration after 144 hours (**G**), time to 1st quantile of the concentration (**H**), time to 2nd quantile of the concentration (**I**), time to 3rd quantile of the concentration (**J**), and the temperature sensitivity (**K**) for each climate class. Each point represents the value for each strain. The brown dots in the middle of the boxplots represent the mean values. The significance levels of the Wilcoxon signed-rank tests with Bonferroni correction are represented by asterisks: * for *p*-value < 0.05, ** for *p*-value < 0.01, and *** for *p*-value < 0.001. Only the climates with more than five strains were represented.

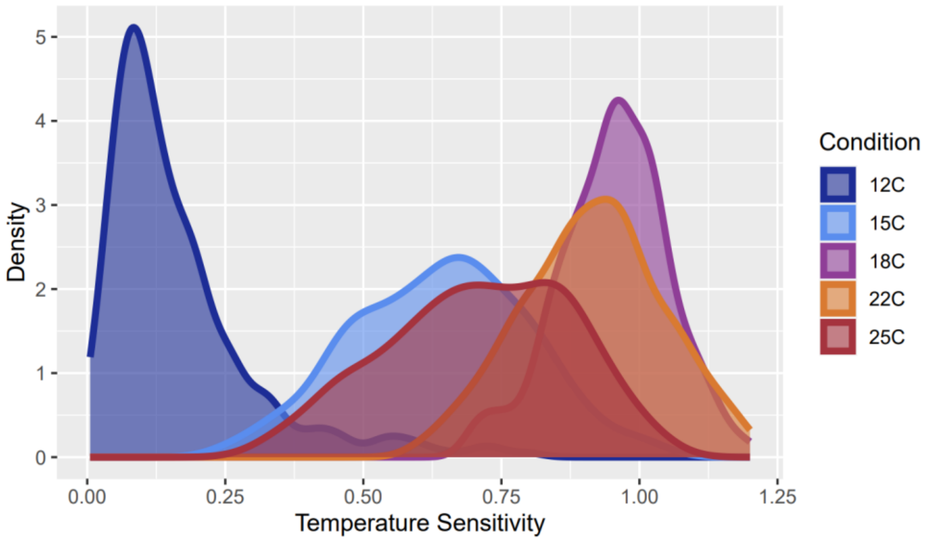


**Supplementary Figure S7.** Distribution of temperature sensitivity estimates per temperature condition tested.

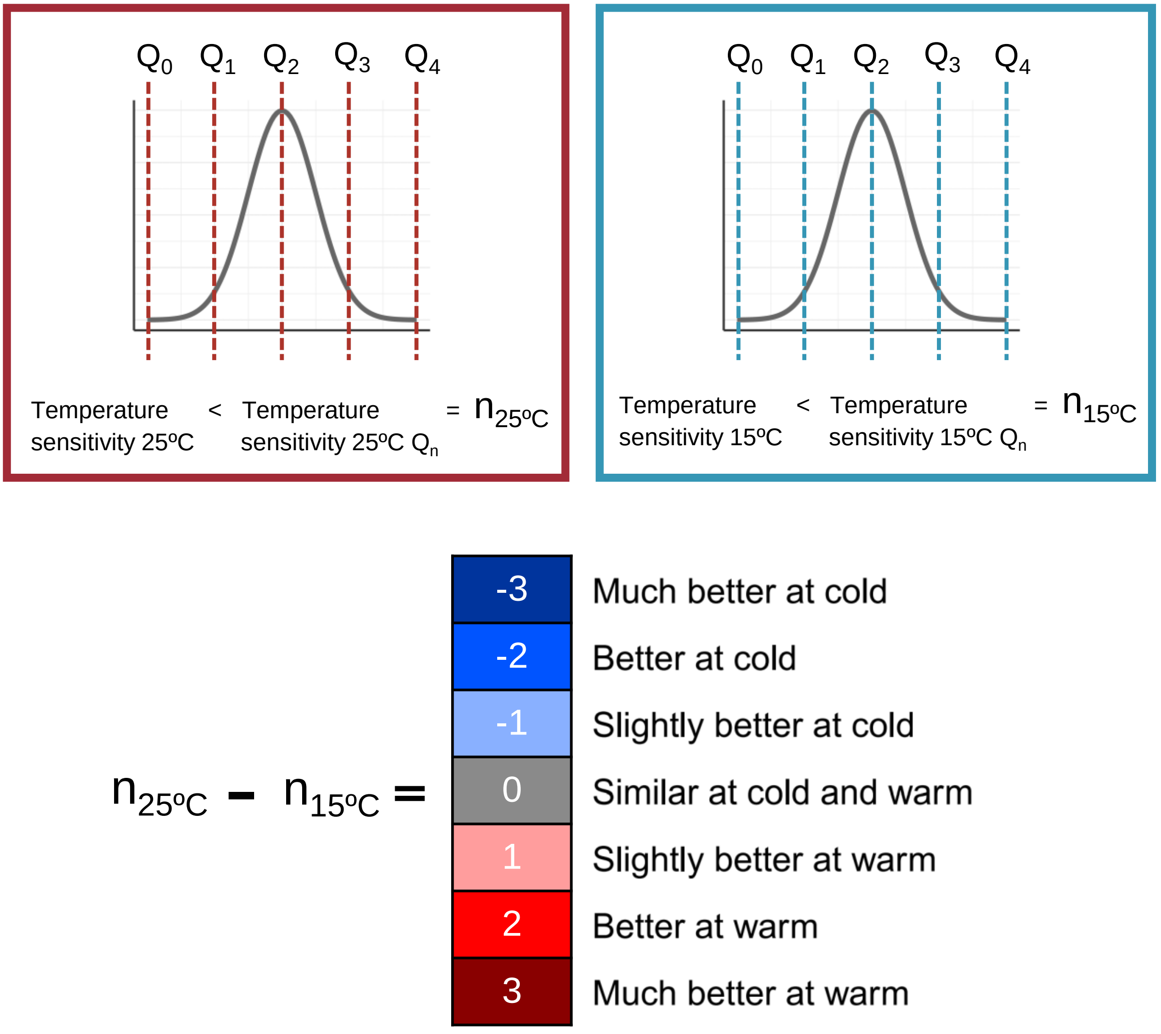


**Supplementary Figure S8.** Diagram of the calculation of the comparative temperature performance for each strain. “Q” represents quartile and “n” represents the quartile number (0-4).

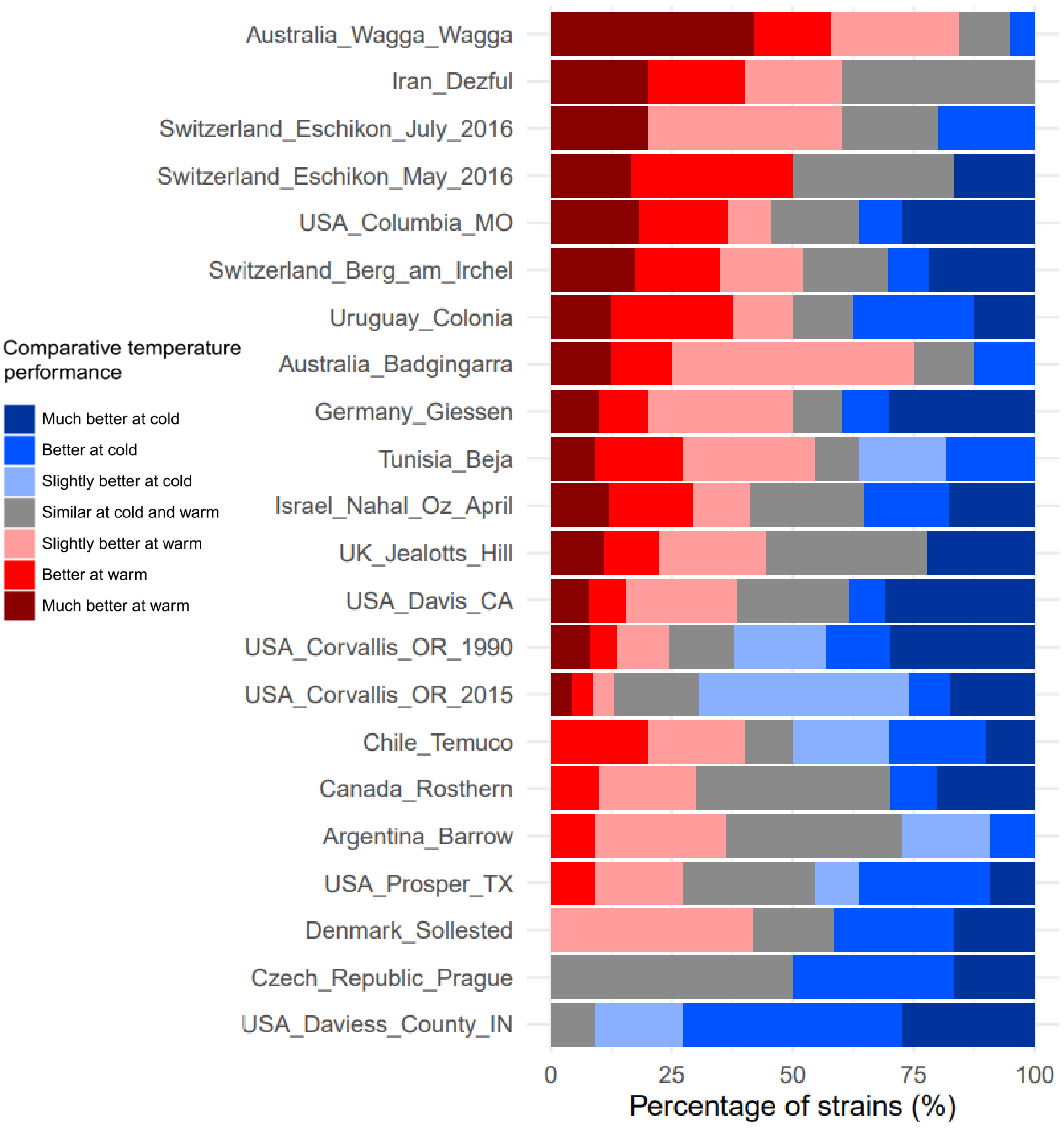


**Supplementary Figure S9.** Percentage of strains per population that perform better at warmer temperatures or at colder temperatures based on the comparison of the temperature sensitivity at 15ºC and the temperature sensitivity at 25ºC. Only populations with more than five strains are shown.

**Supplementary Figures S10-S66.** Manhattan plots of the genotype-phenotypes associations for the 12 growth variables per temperature condition. Values obtained from pipeline *vcf2gwas* through the software GEMMA using the default linear model. The darkest color of the dots represents if the variant is significant (FDR 10% threshold) and the dotted horizontal lines indicate the Bonferroni thresholds.

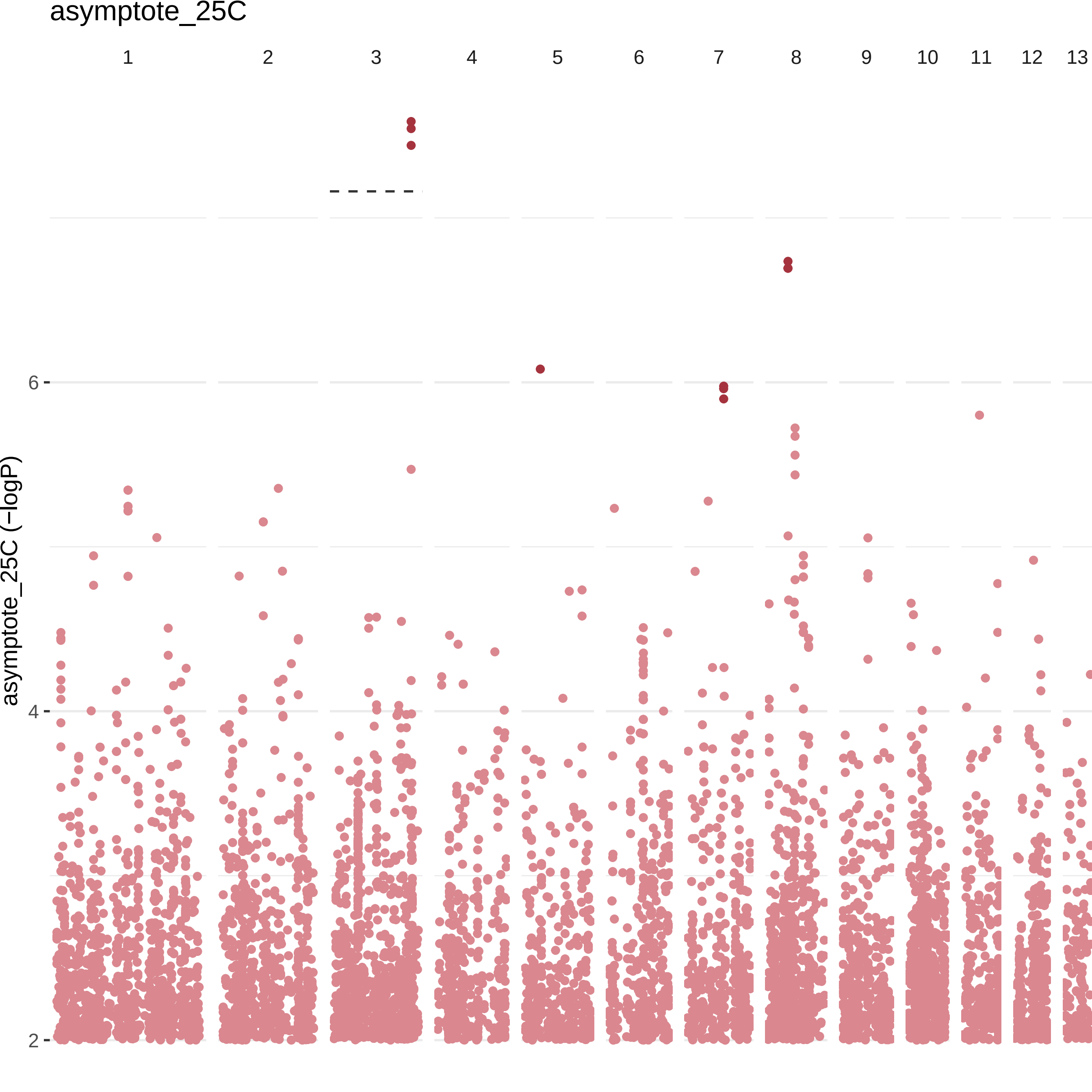

asymptote_25C
1
2
3
4
5
6
7
8
9
10
11
12
13
6
4
2
asymptote_25C (−logP)

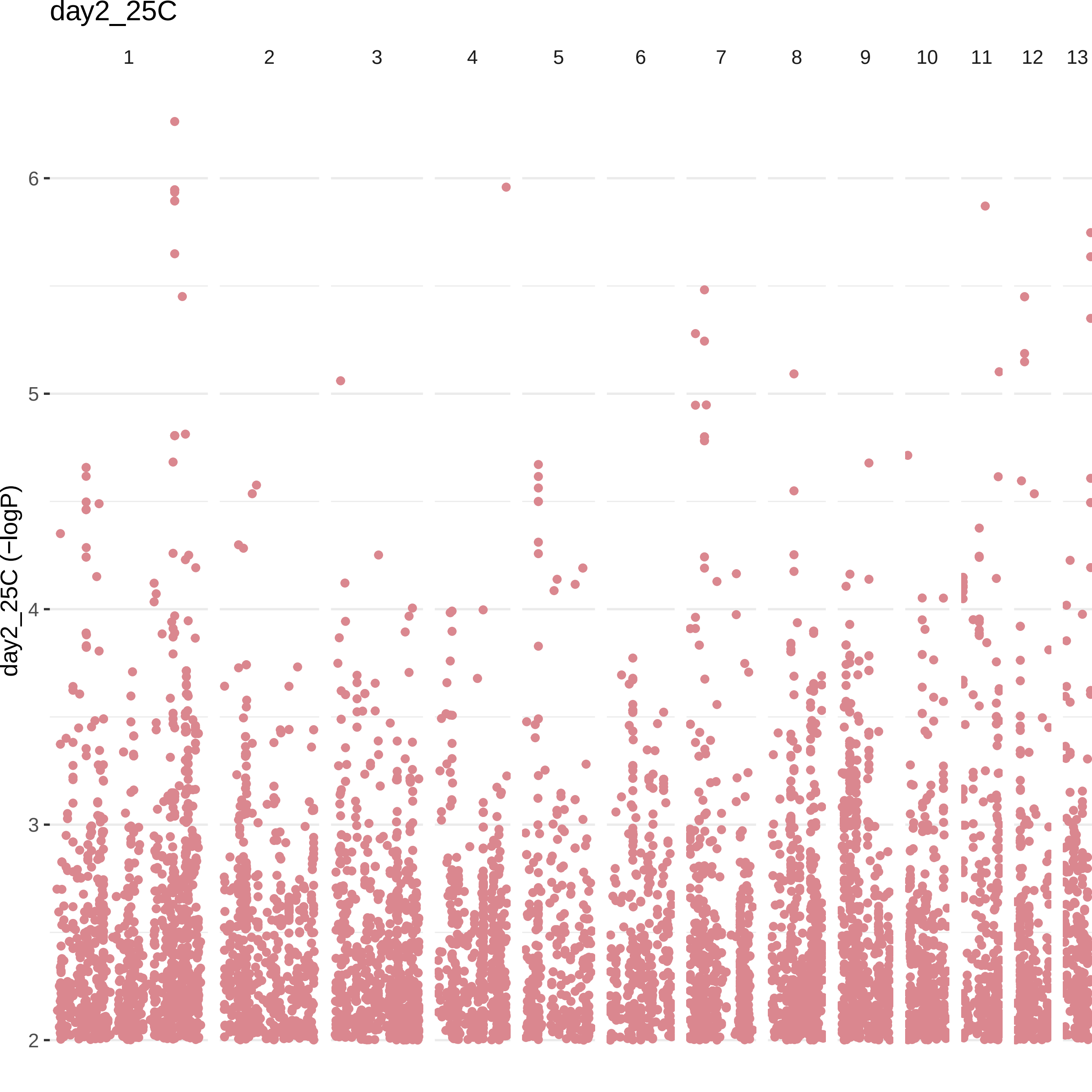
day2_25C
1
2
3
4
5
6
7
8
9
10
11
12
13
6
5
4
3
2
day2_25C (−logP)

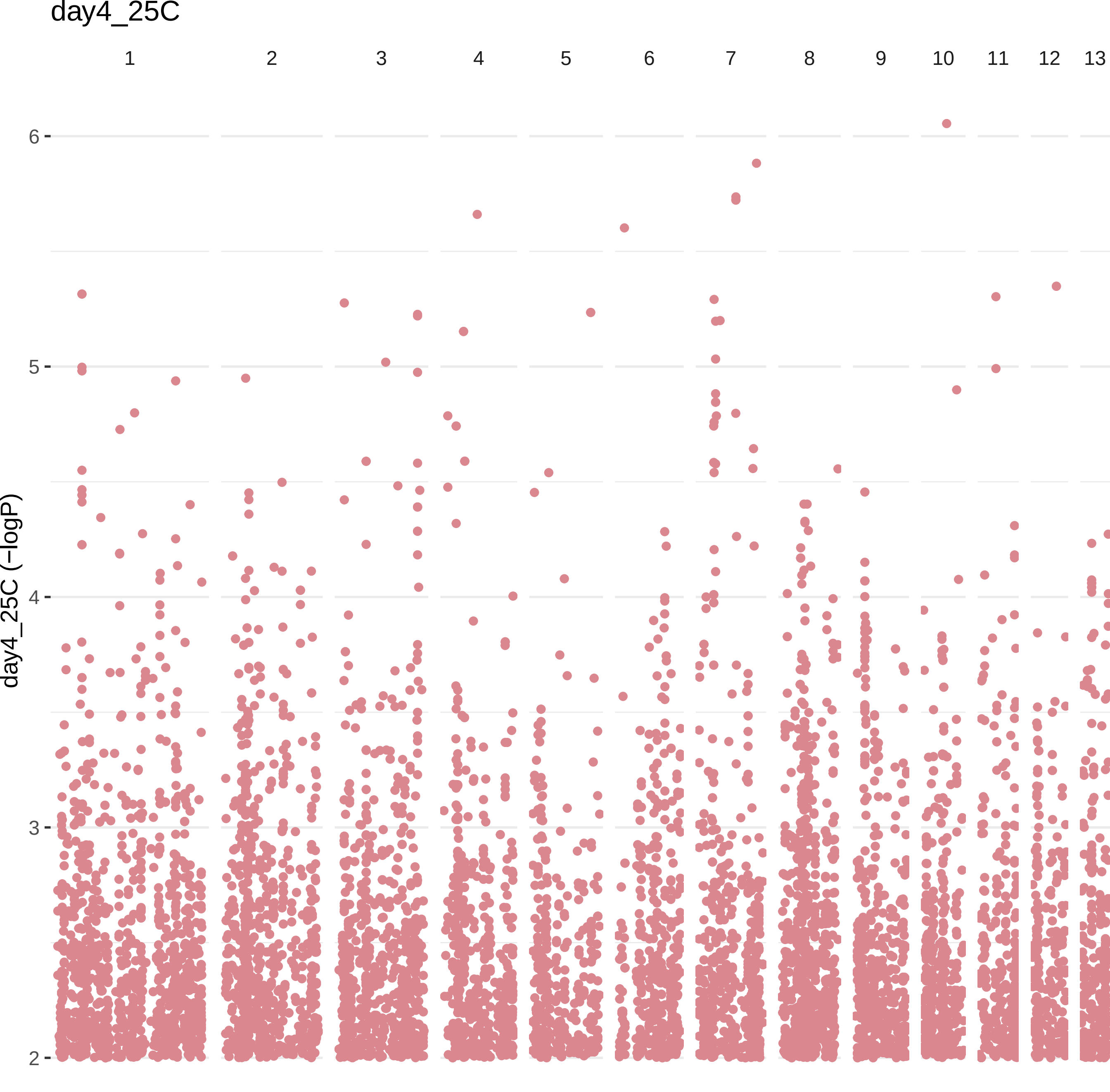
day4_25C
1
2
3
4
5
6
7
8
9
10
11
12
13
6
5
4
3
2
day4_25C (−logP)

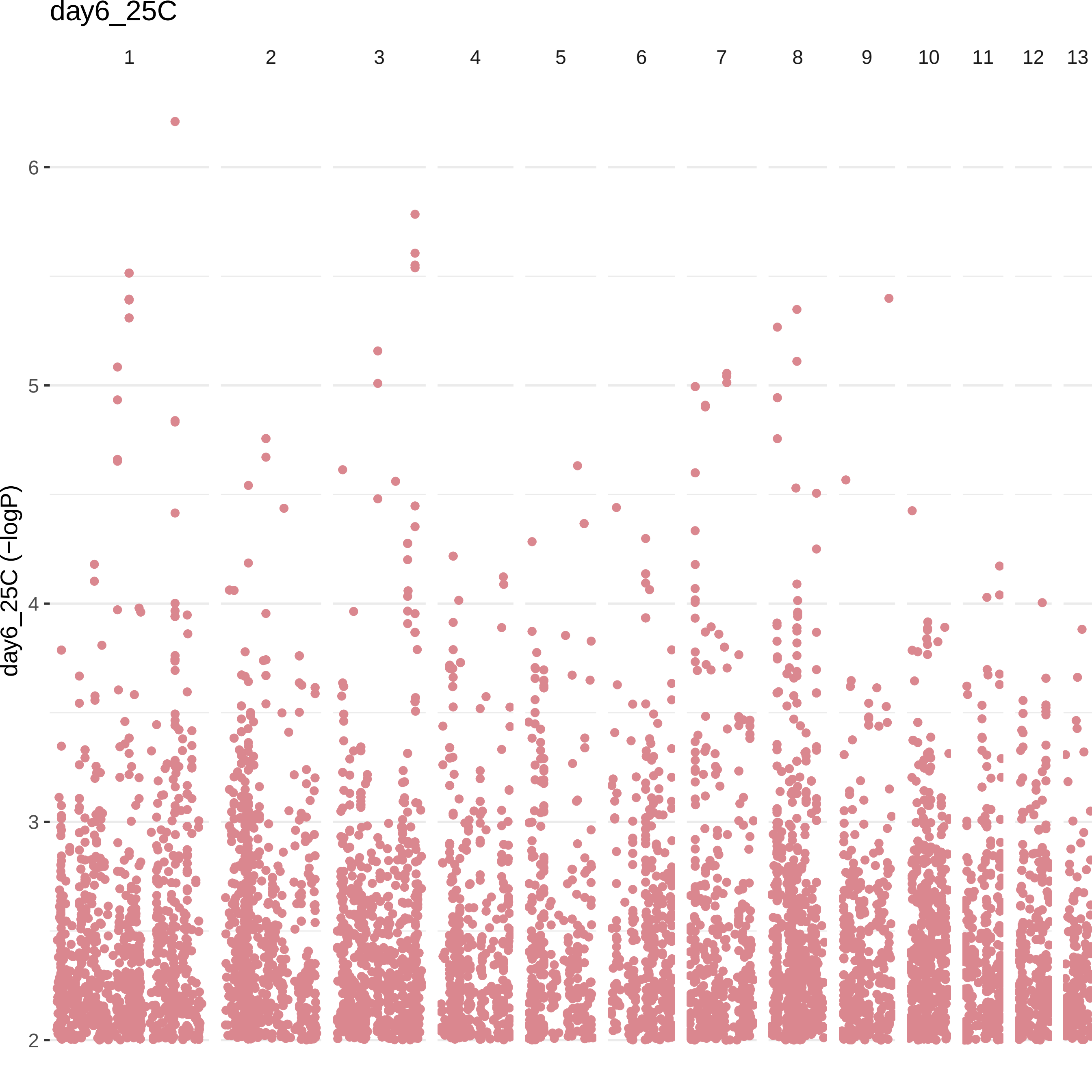
day6_25C
1
2
3
4
5
6
7
8
9
10
11
12
13
day6_25C (−logP)
6
5
4
3
2

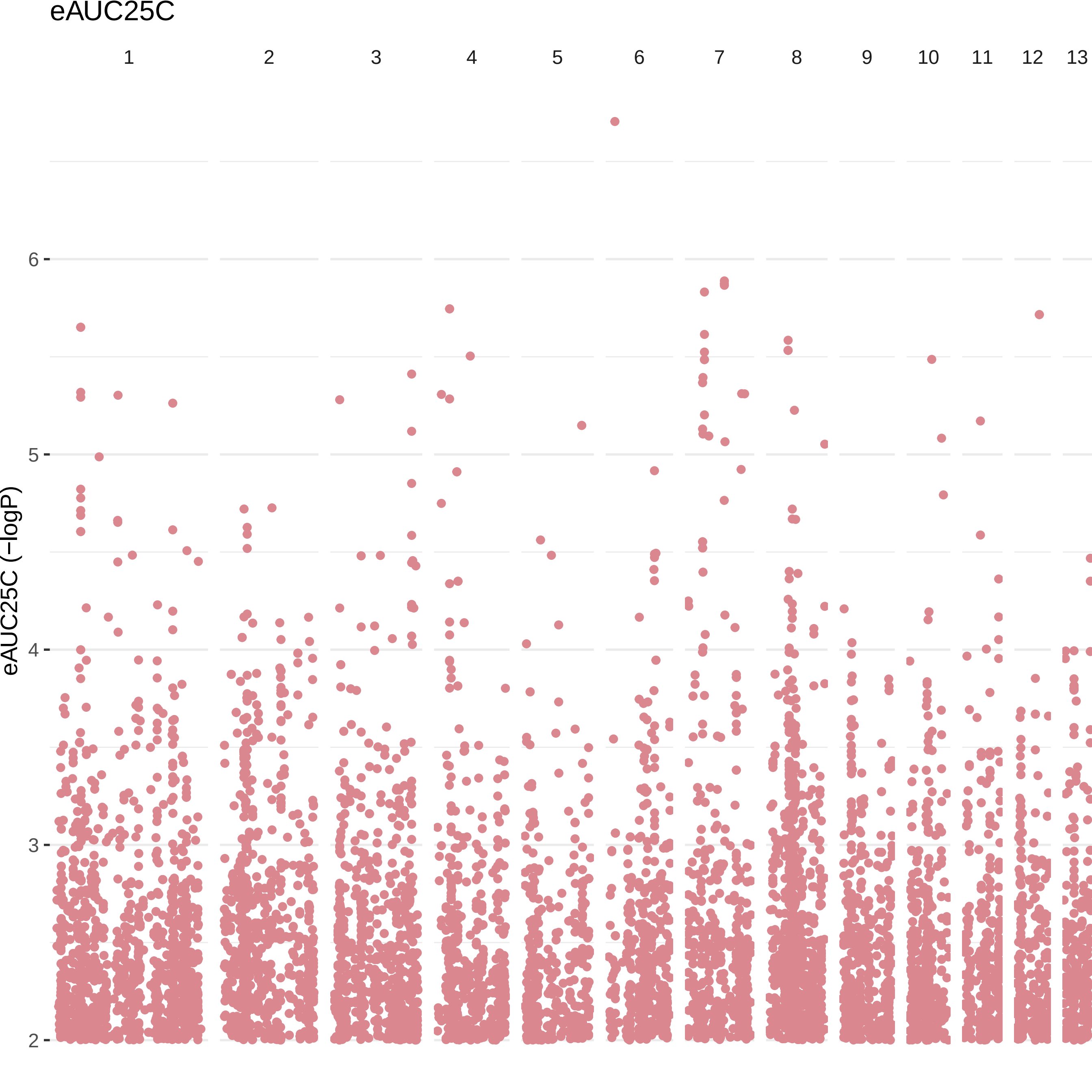
eAUC25C
1
2
3
4
5
6
7
8
9
10
11
12
13
eAUC25C (−logP)
6
5
4
3
2

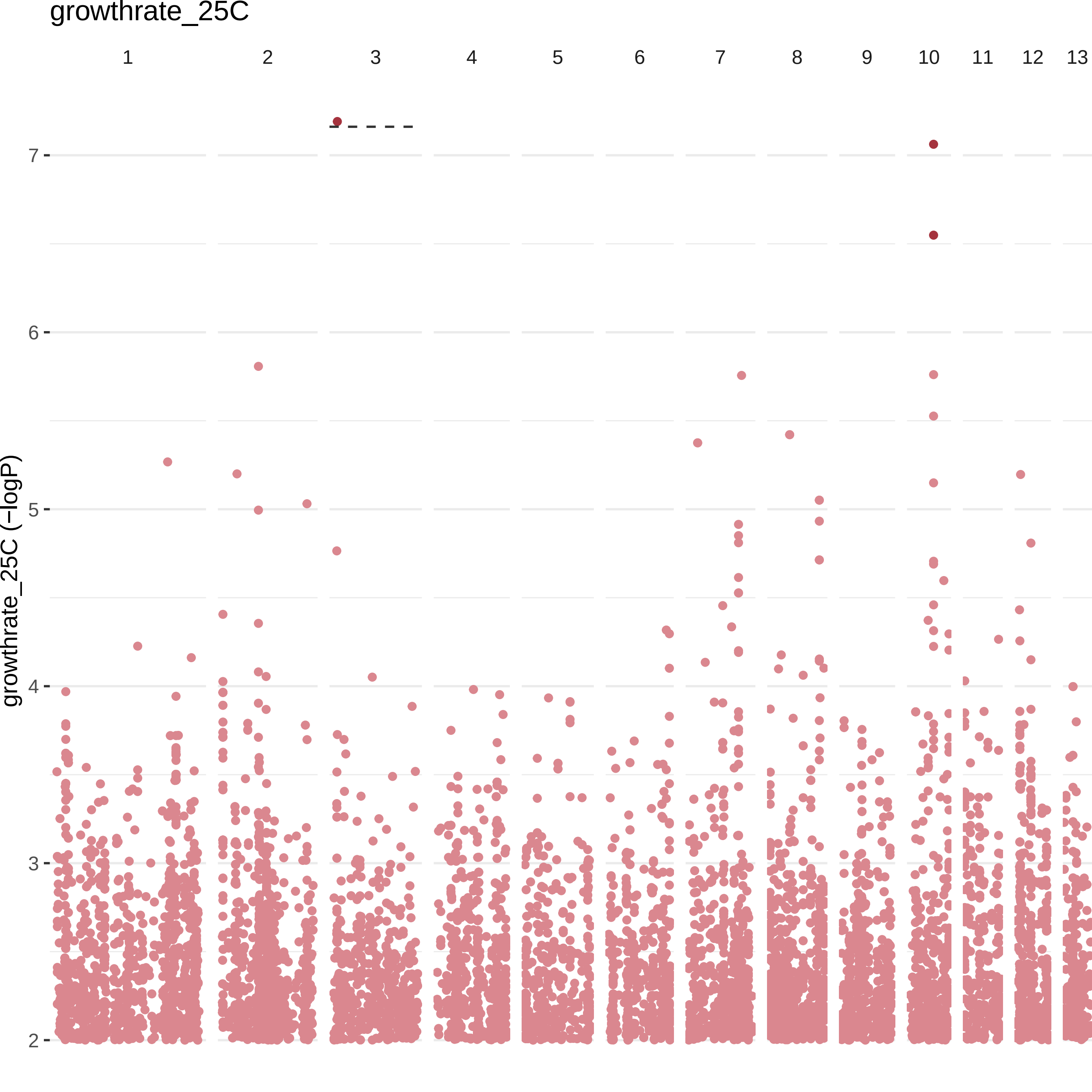

growthrate_25C
1
2
3
4
5
6
7
8
9
10
11
12
13
growthrate_25C (−logP)
7
6
5
4
3
2

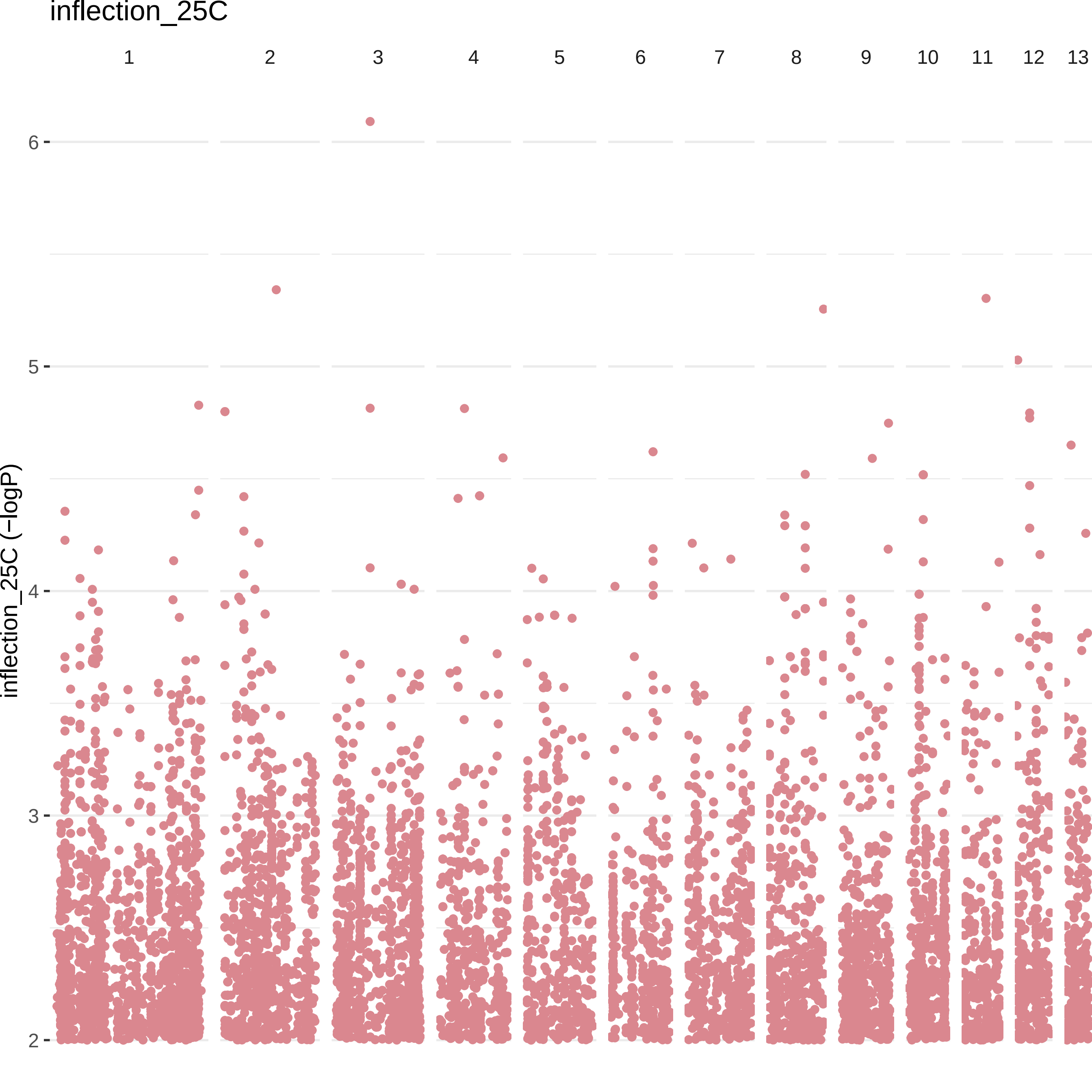

inflection_25C
1
2
3
4
5
6
7
8
9
10
11
12
13
6
5
4
3
2
inflection_25C (−logP)

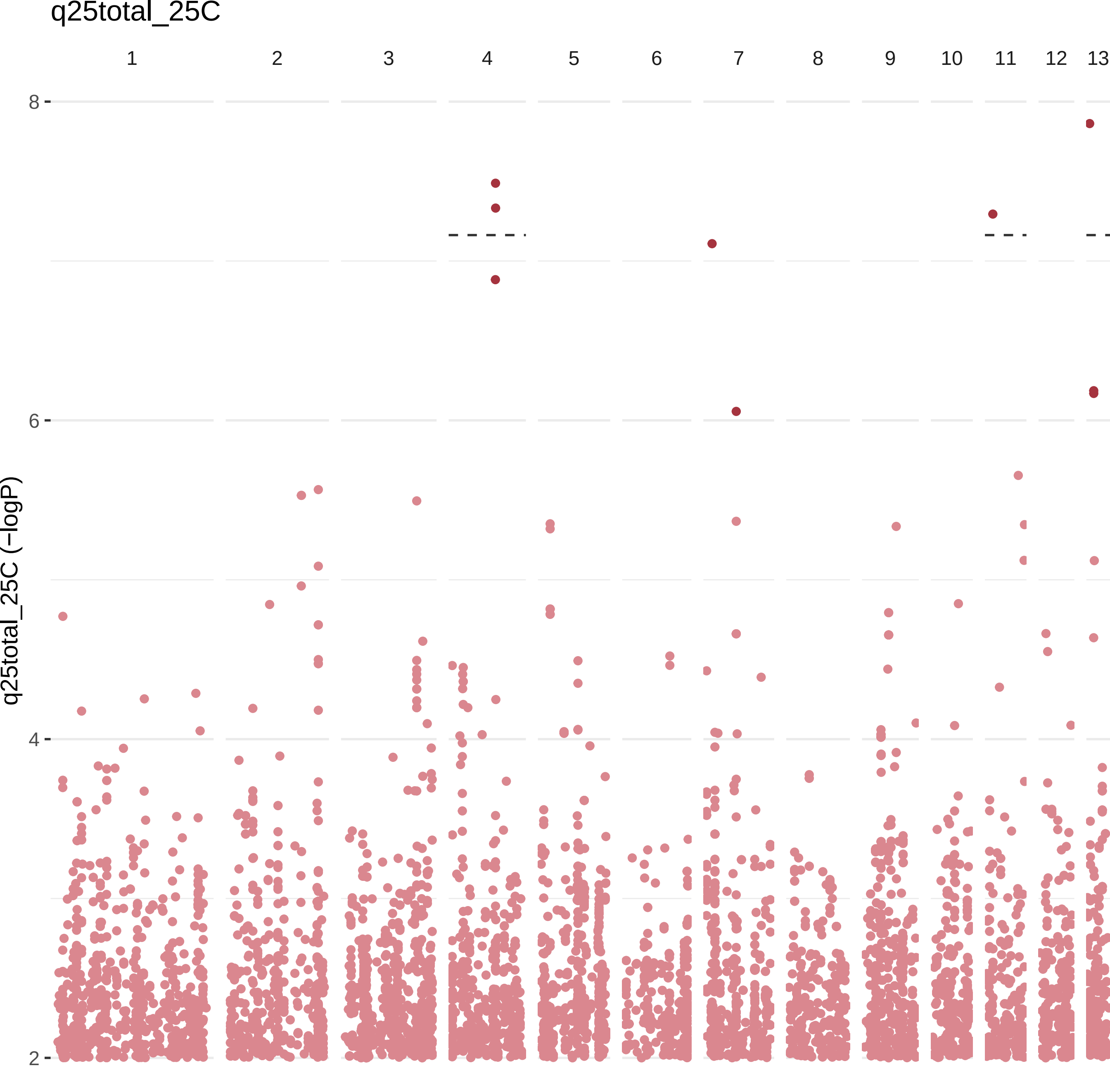
q25total_25C
1
2
3
4
5
6
7
8
9
10
11
12
13
8
6
4
2
q25total_25C (−logP)

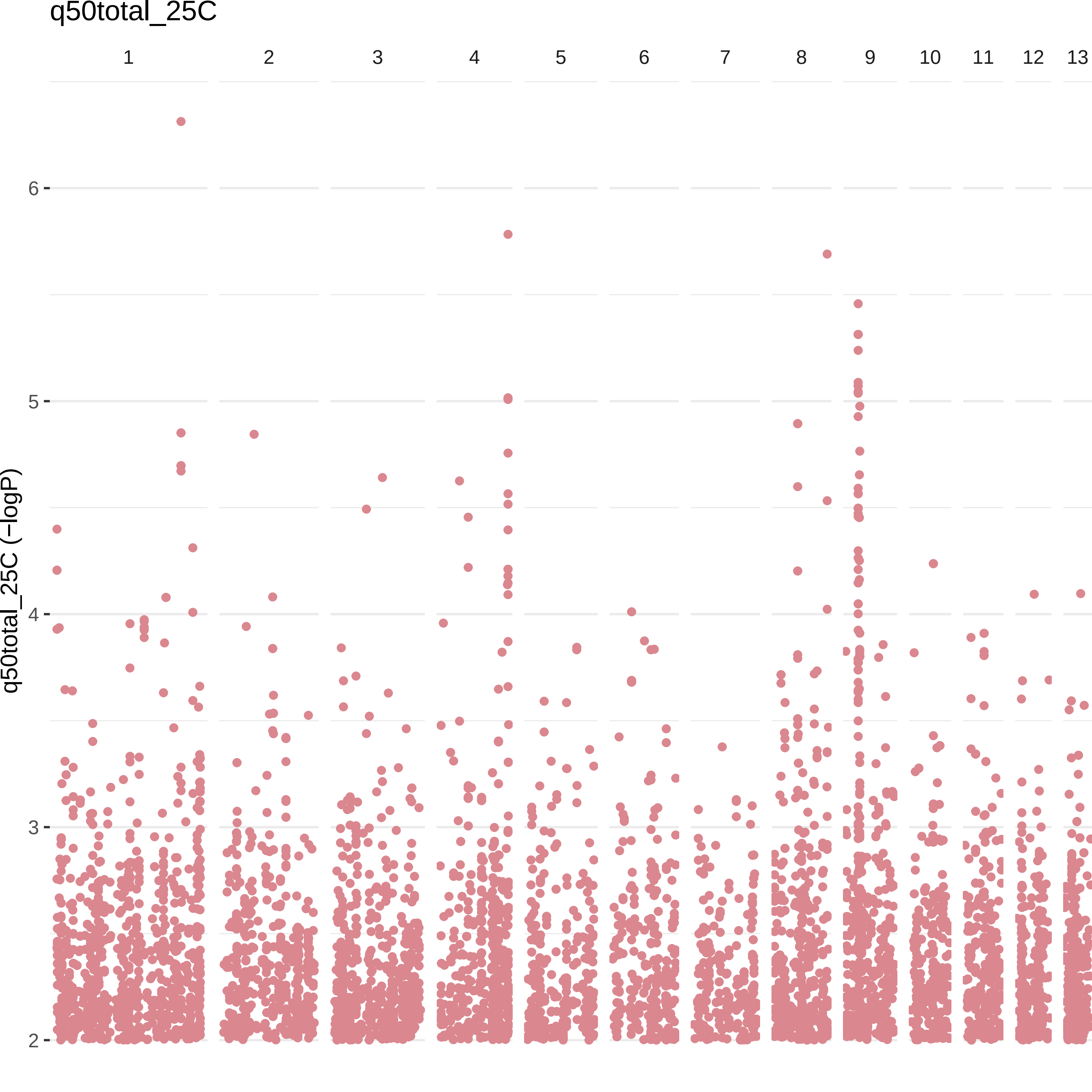

q50total_25C
1
2
3
4
5
6
7
8
9
10
11
12
13
6
5
4
3
2
q50total_25C (−logP)

q75total_25C

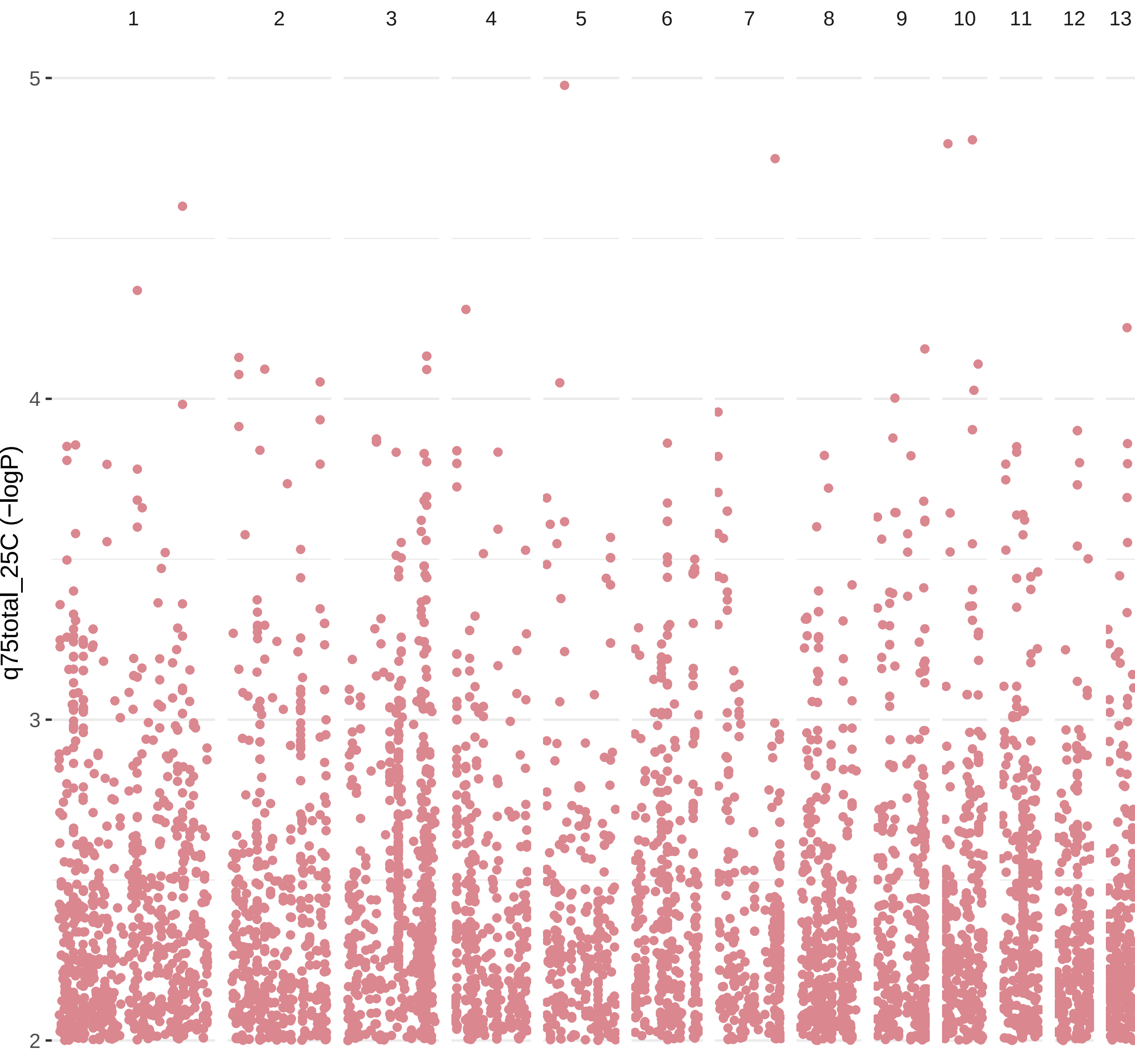

Tempsens25C

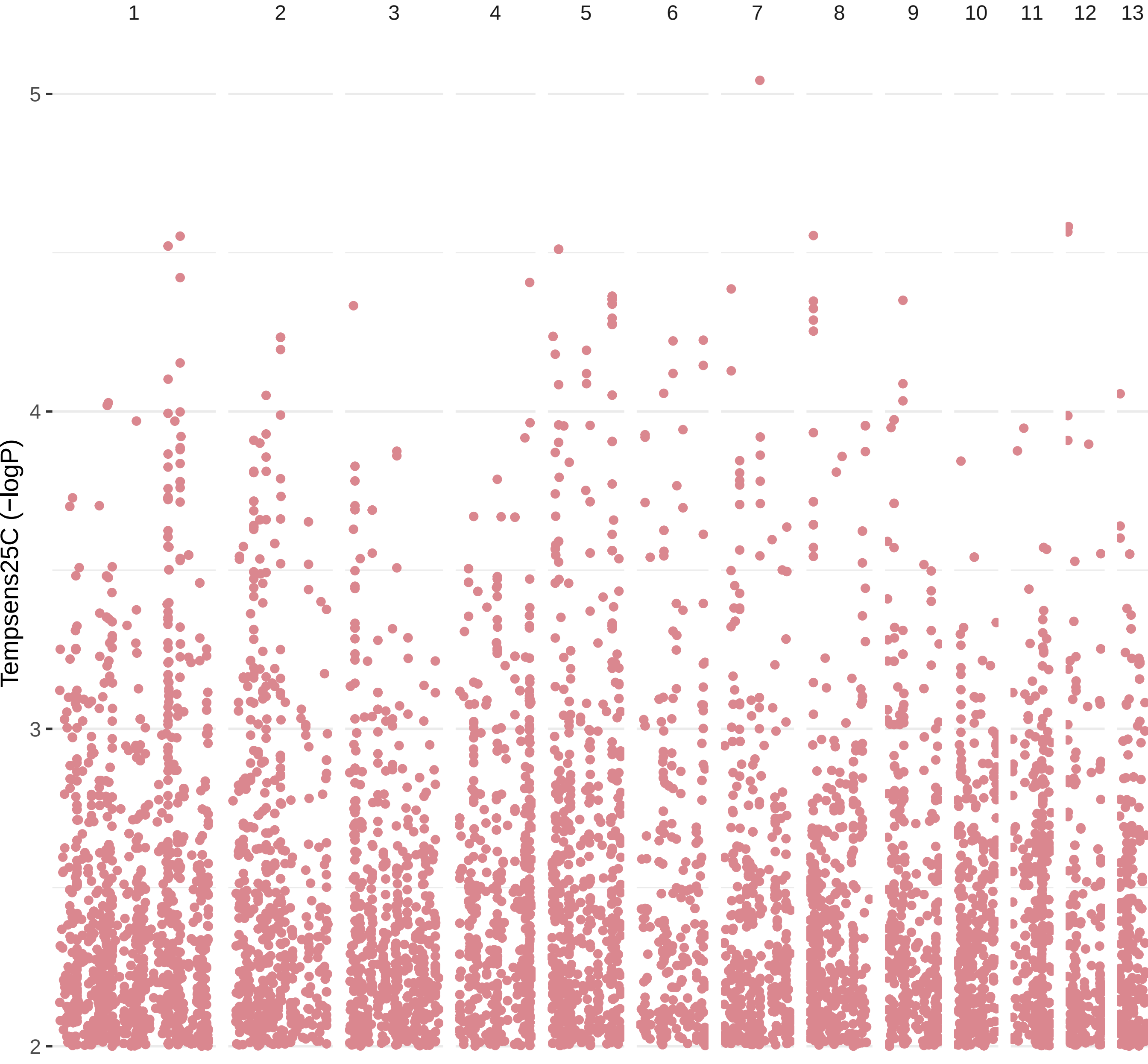

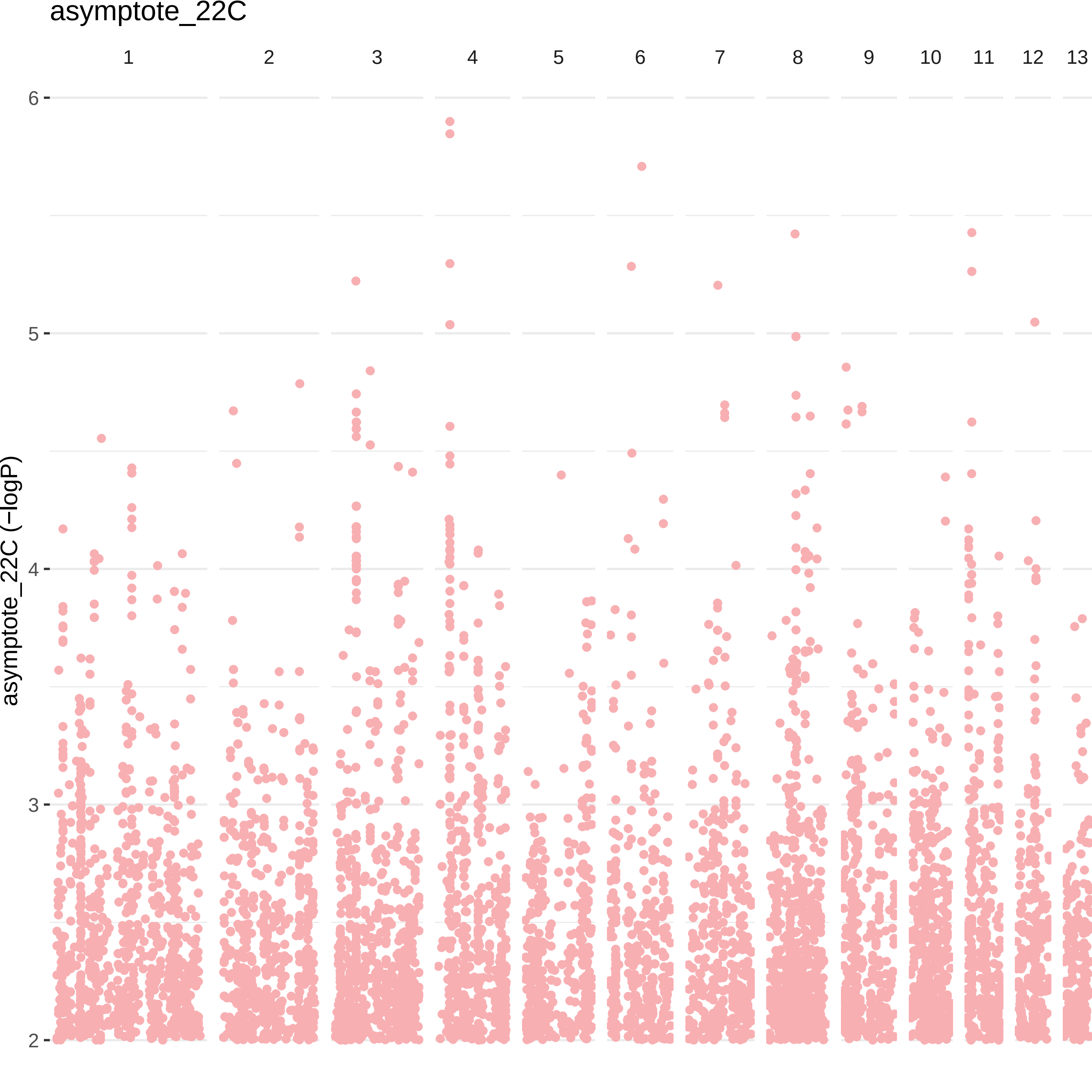

asymptote_22C
1
2
3
4
5
6
7
8
9
10
11
12
13
asymptote_22C (−logP)
6
5
4
3
2

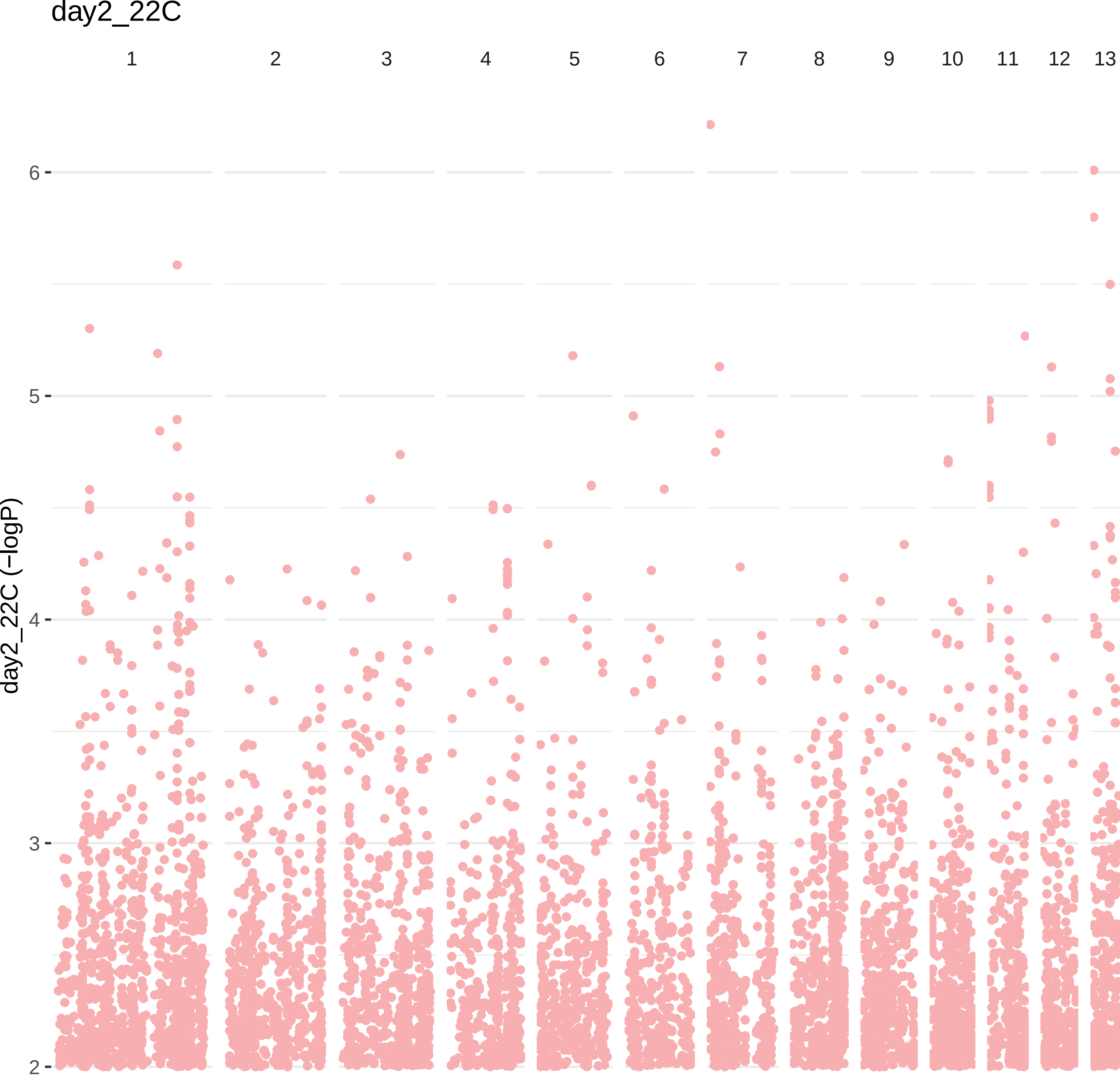

day2_22C
1
2
3
4
5
6
7
8
9
10
11
12
13
6
5
4
3
2
day2_22C (−logP)

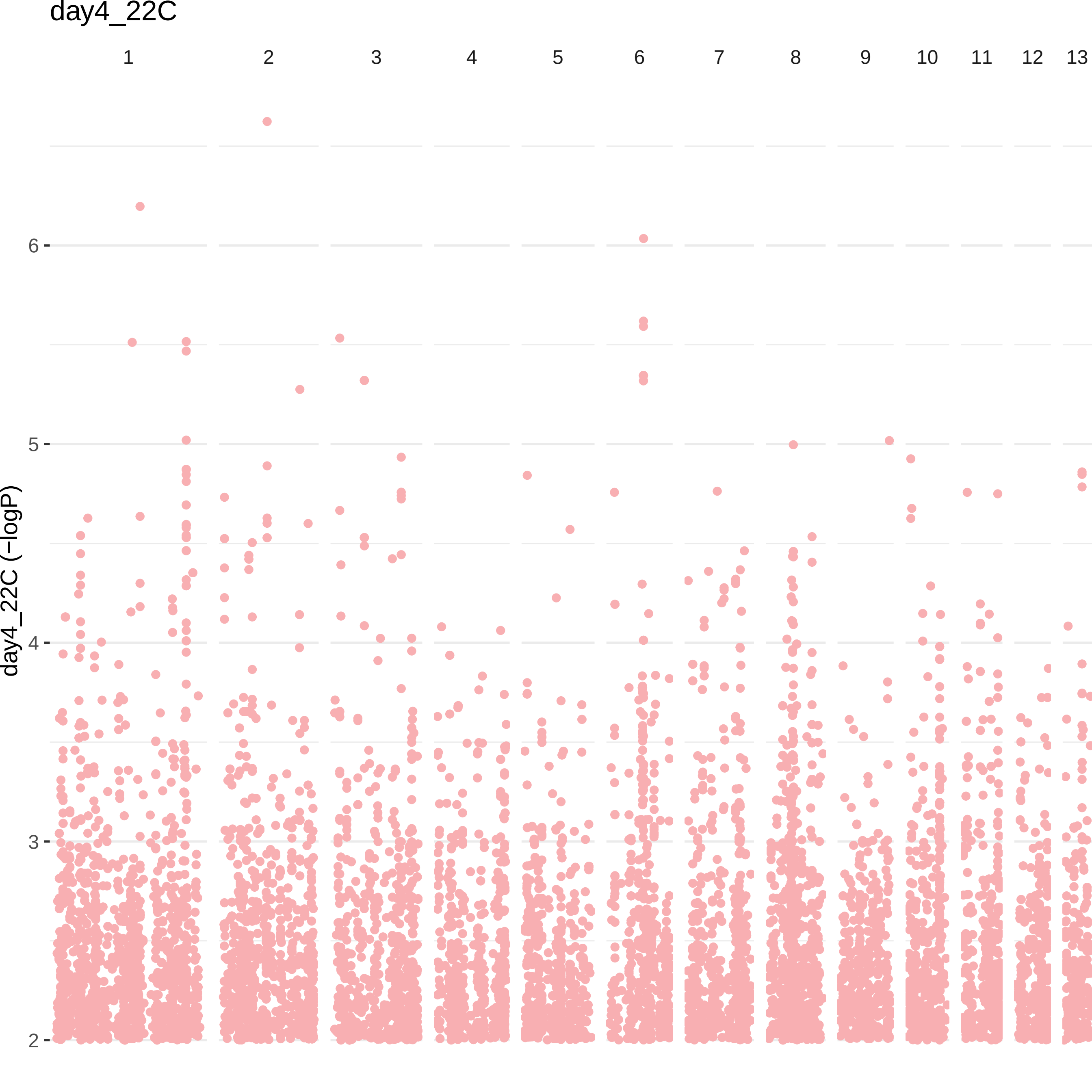
day4_22C
1
2
3
4
5
6
7
8
9
10
11
12
13
day4_22C (−logP)
6
5
4
3
2

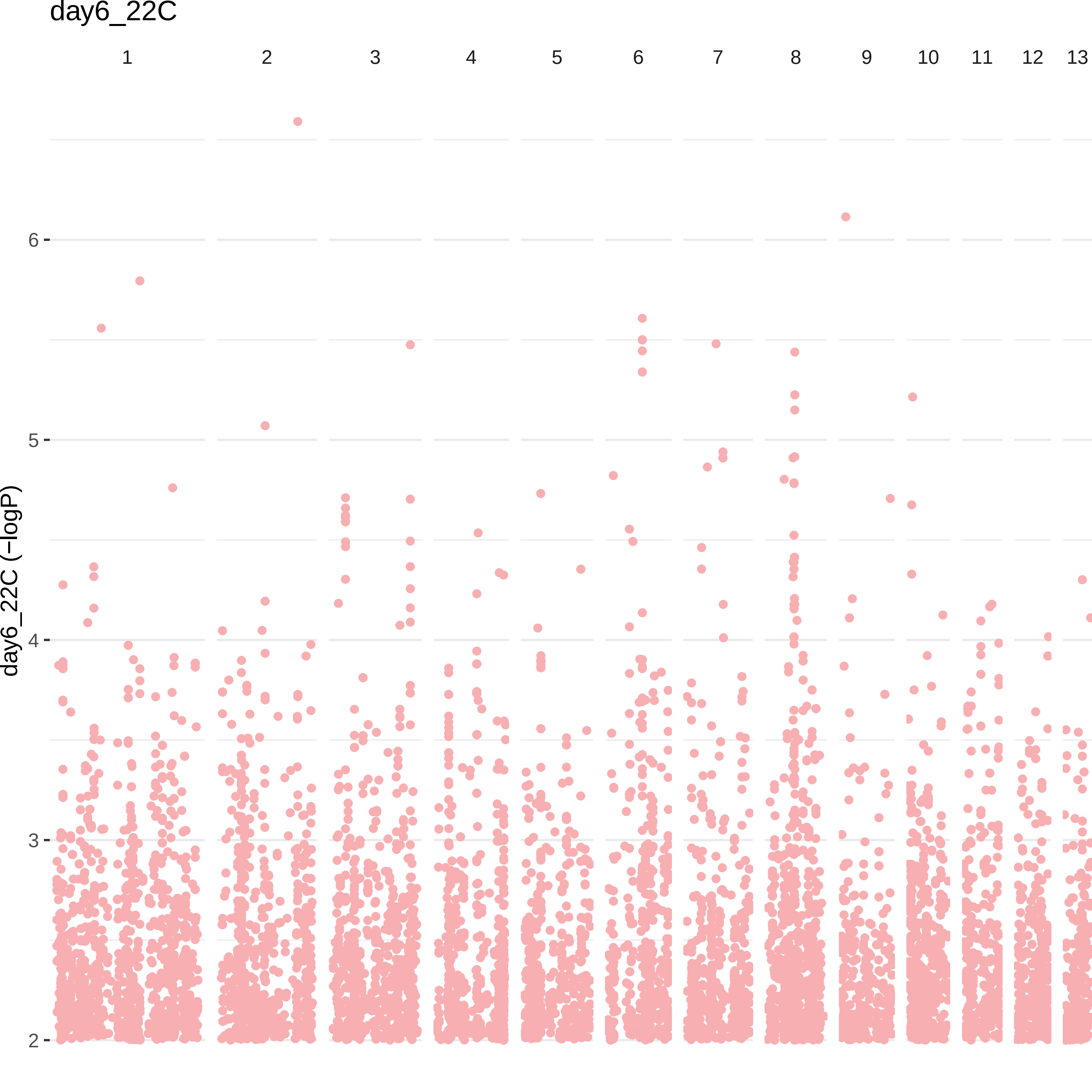

day6_22C
1
2
3
4
5
6
7
8
9
10
11
12
13
day6_22C (−logP)
6
5
4
3
2

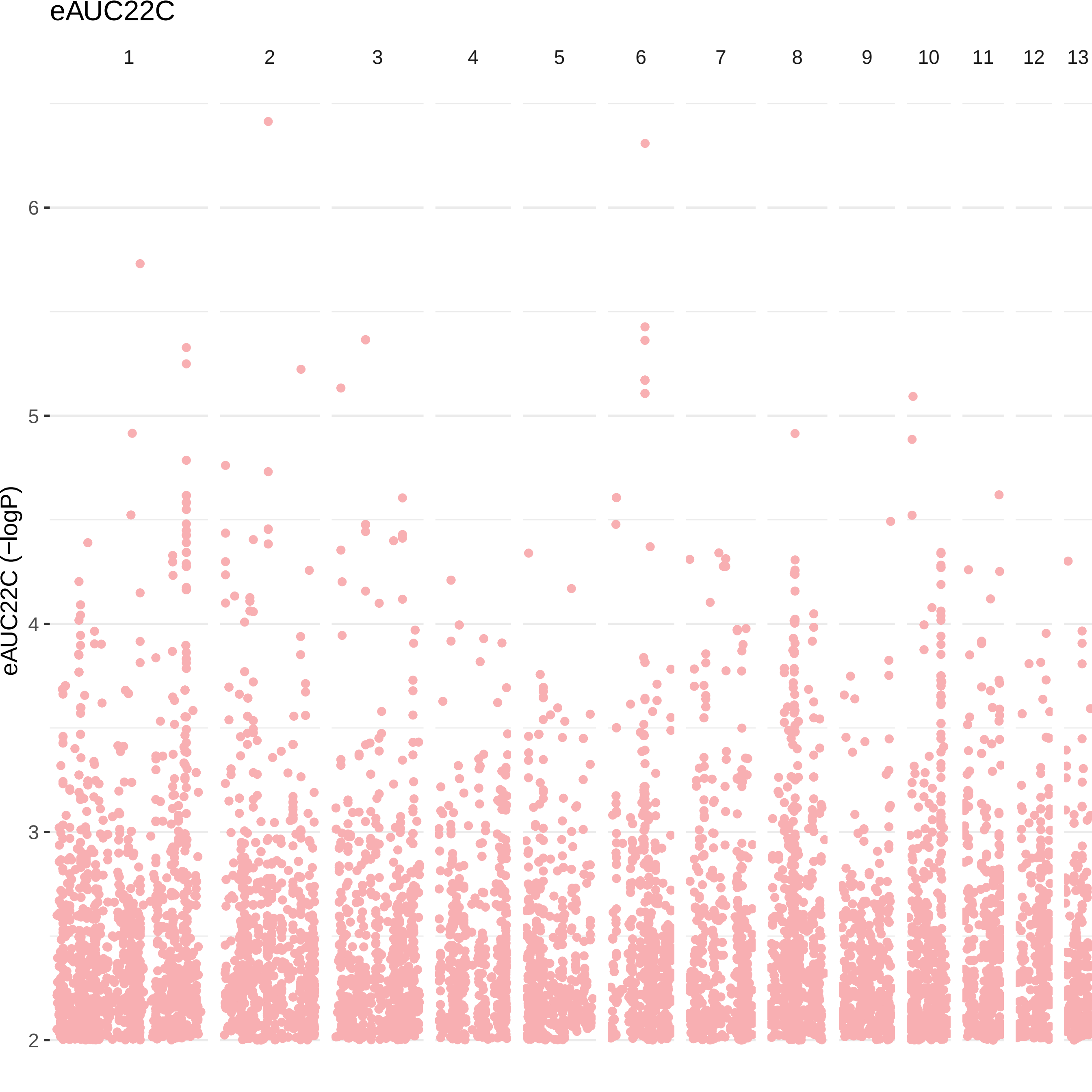
eAUC22C
1
2
3
4
5
6
7
8
9
10
11
12
13
eAUC22C (−logP)
6
5
4
3
2

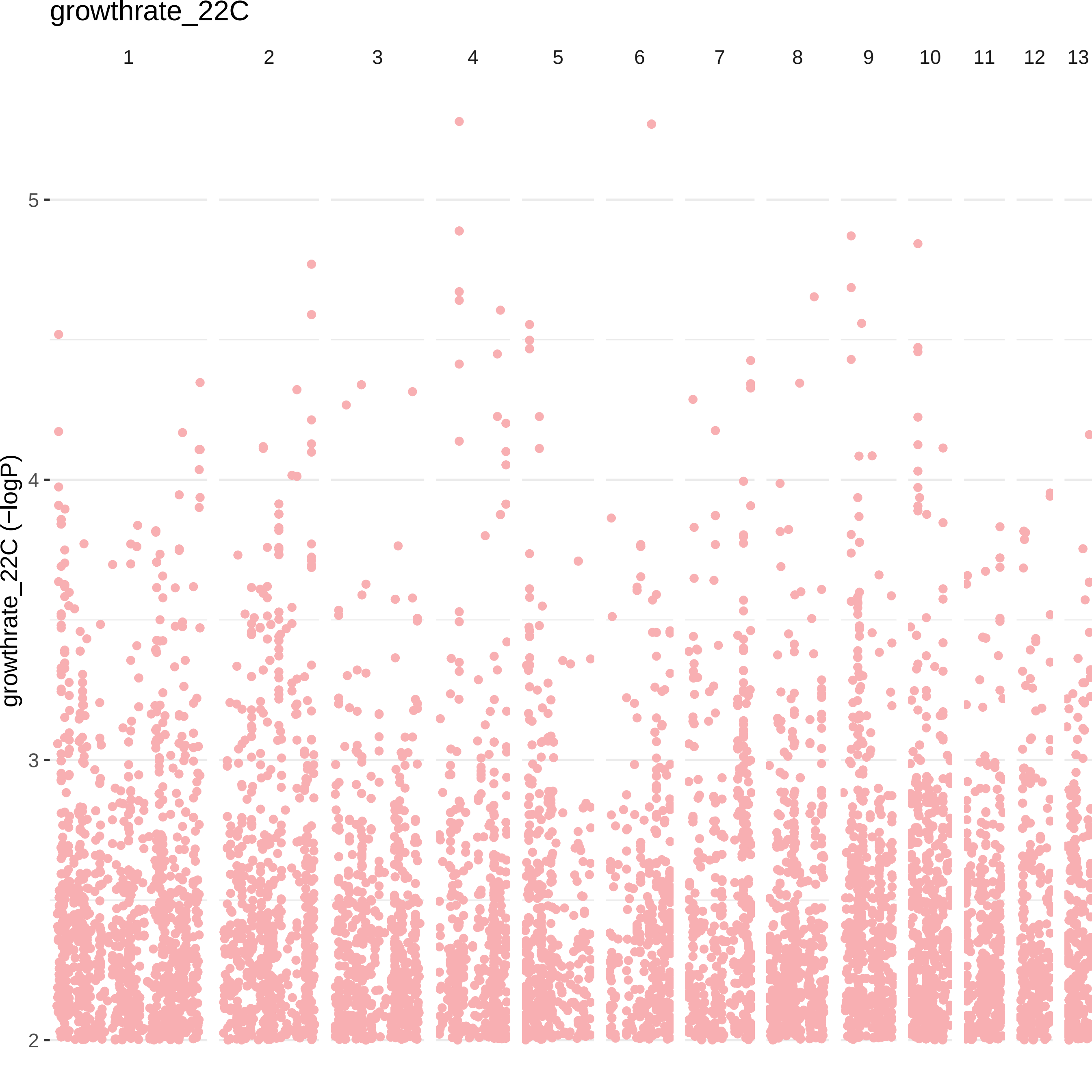
growthrate_22C
1
2
3
4
5
6
7
8
9
10
11
12
13
growthrate_22C (−logP)
5
4
3
2

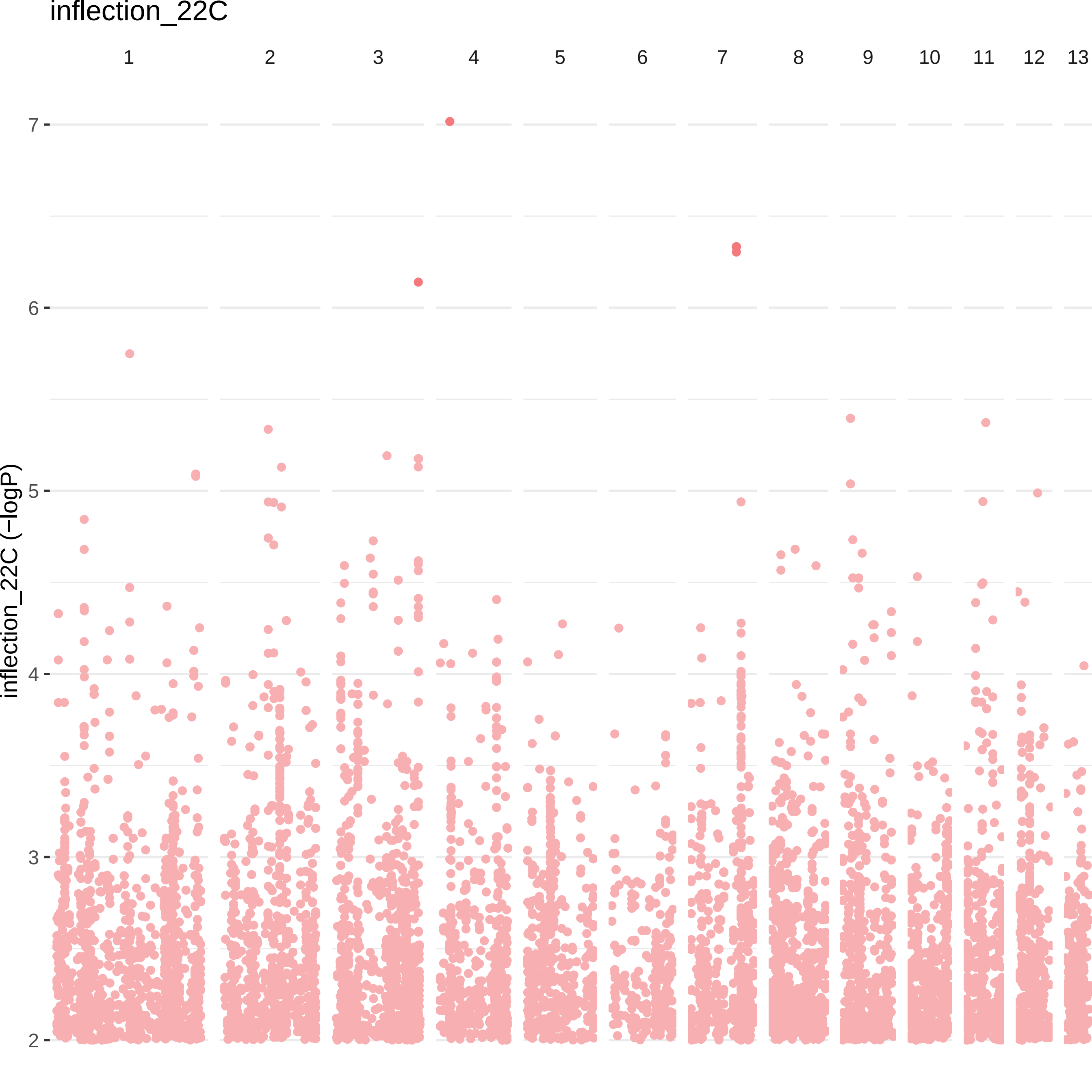

inflection_22C
1
2
3
4
5
6
7
8
9
10
11
12
13
inflection_22C (−logP)
7
6
5
4
3
2

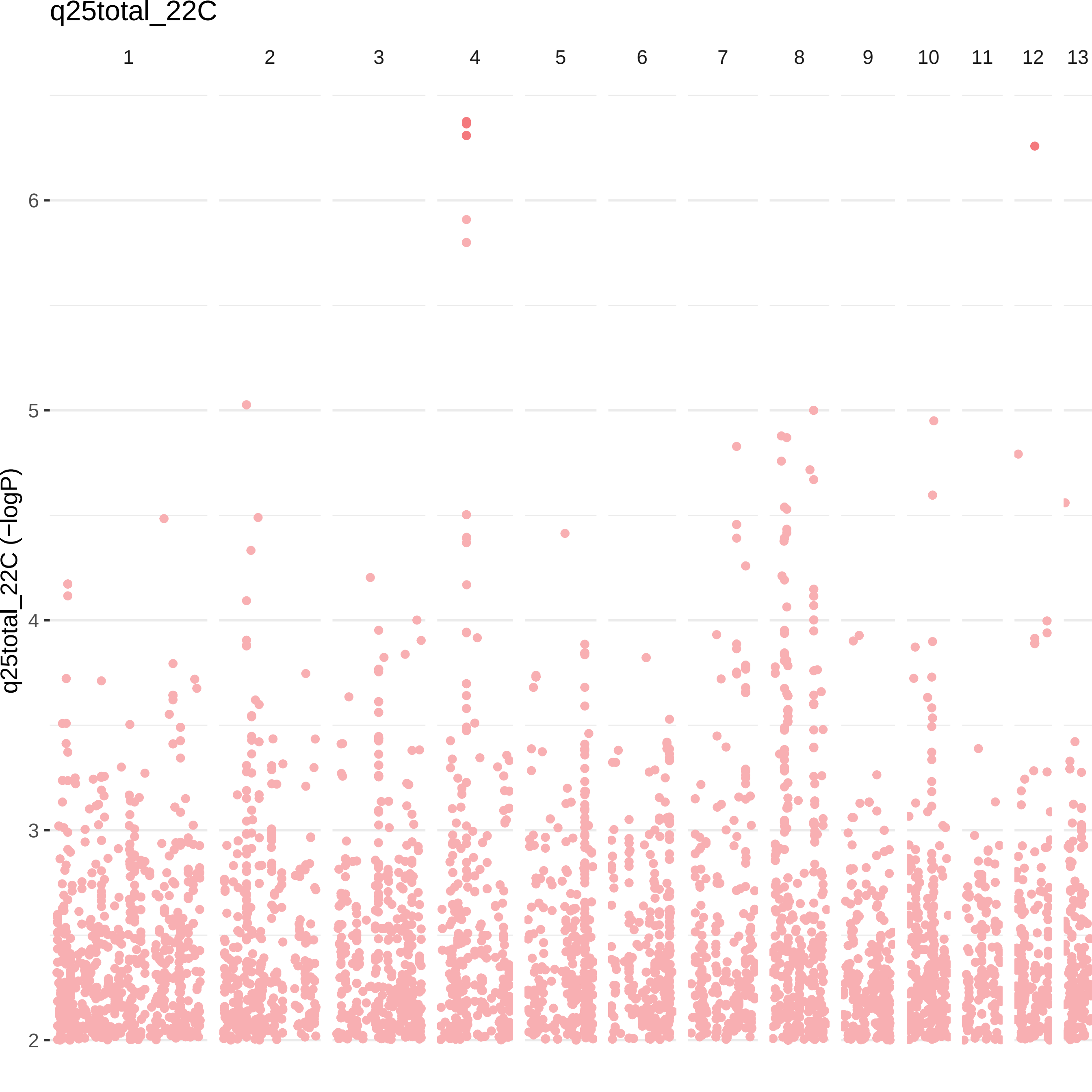

q25total_22C
1
2
3
4
5
6
7
8
9
10
11
12
13
q25total_22C (−logP)
6
5
4
3
2

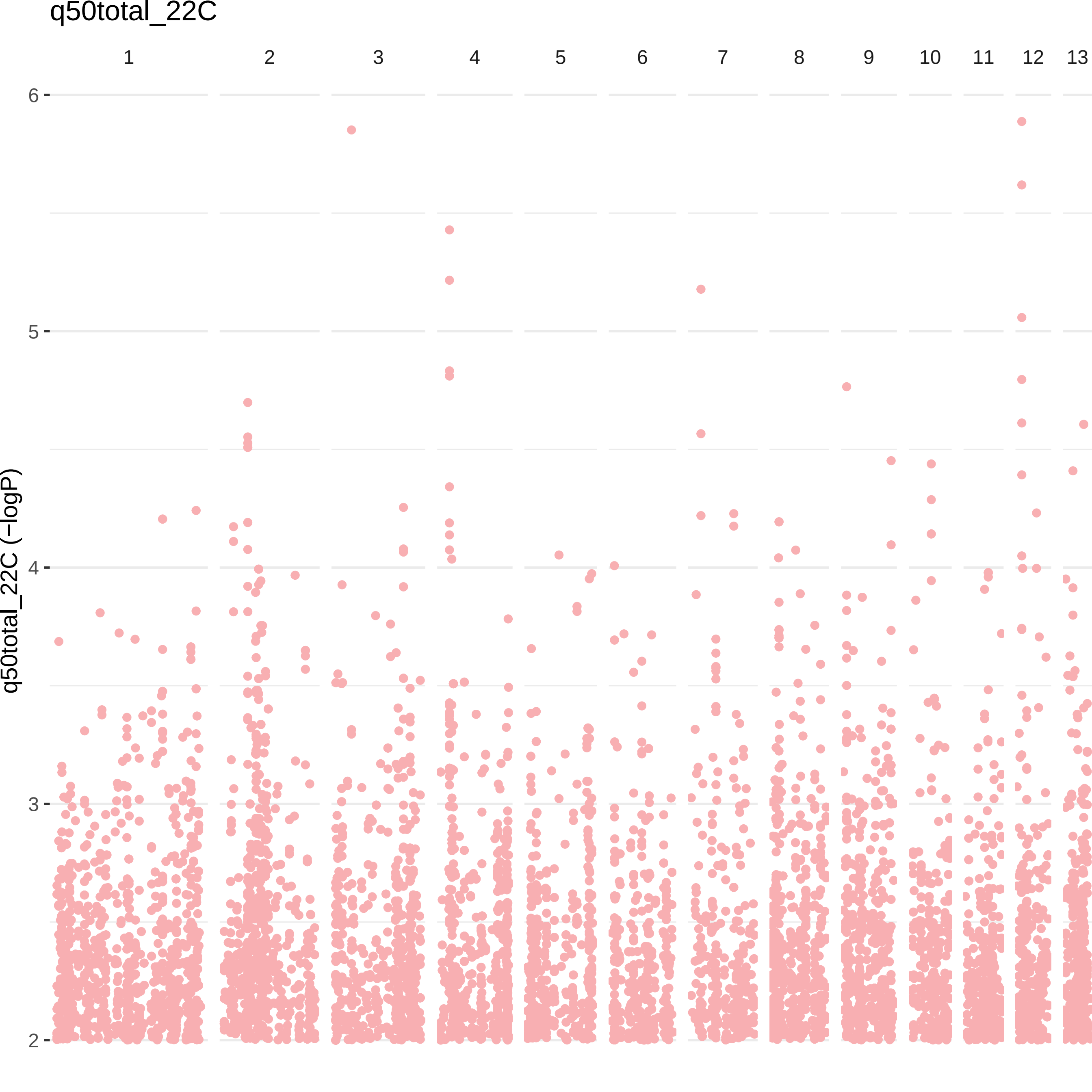

q50total_22C
1
2
3
4
5
6
7
8
9
10
11
12
13
6
5
4
3
2
q50total_22C (−logP)

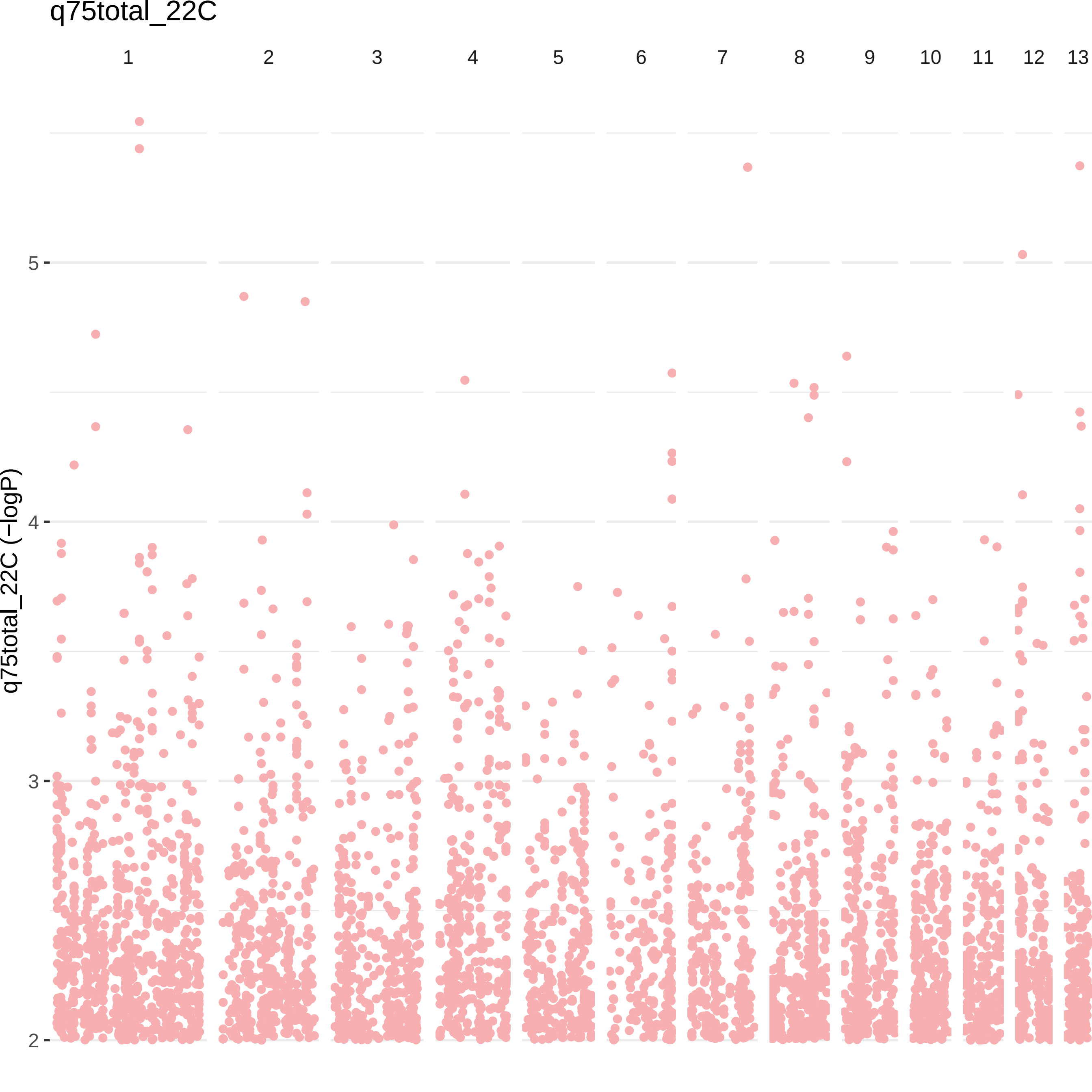

q75total_22C
1
2
3
4
5
6
7
8
9
10
11
12
13
q75total_22C (−logP)
5
4
3
2

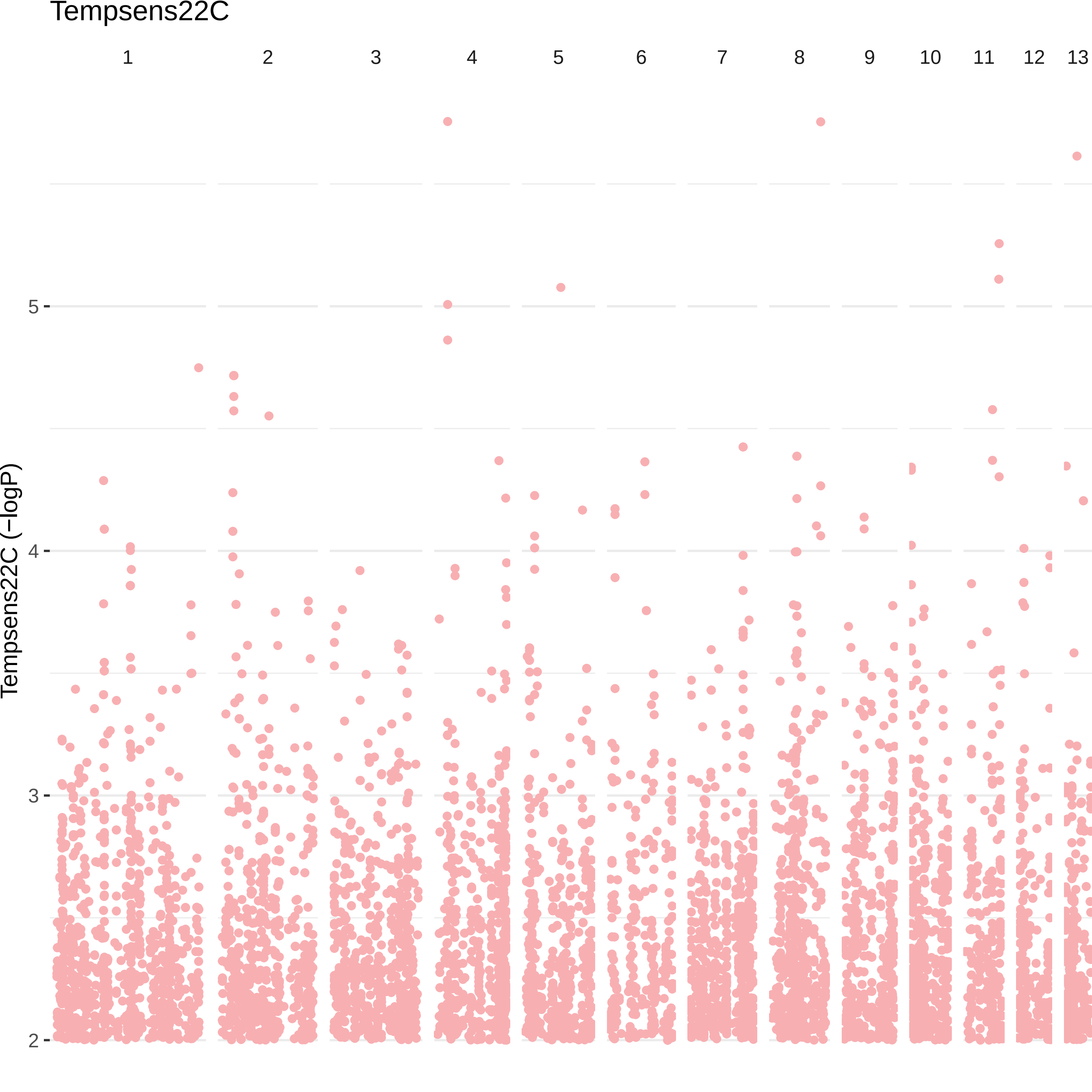

Tempsens22C
1
2
3
4
5
6
7
8
9
10
11
12
13
Tempsens22C (−logP)
5
4
3
2

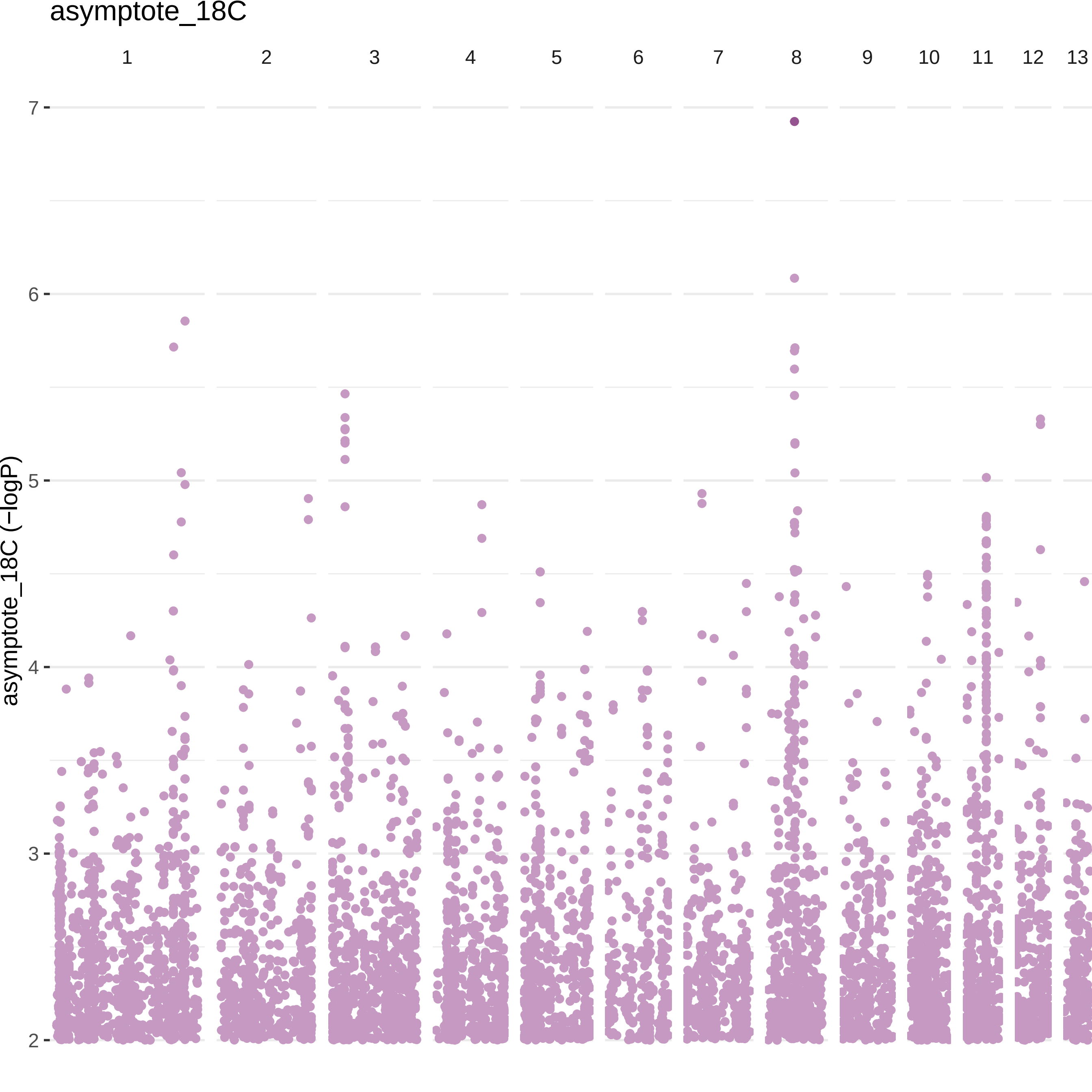

asymptote_18C
1
2
3
4
5
6
7
8
9
10
11
12
13
7
6
5
4
3
2
asymptote_18C (−logP)

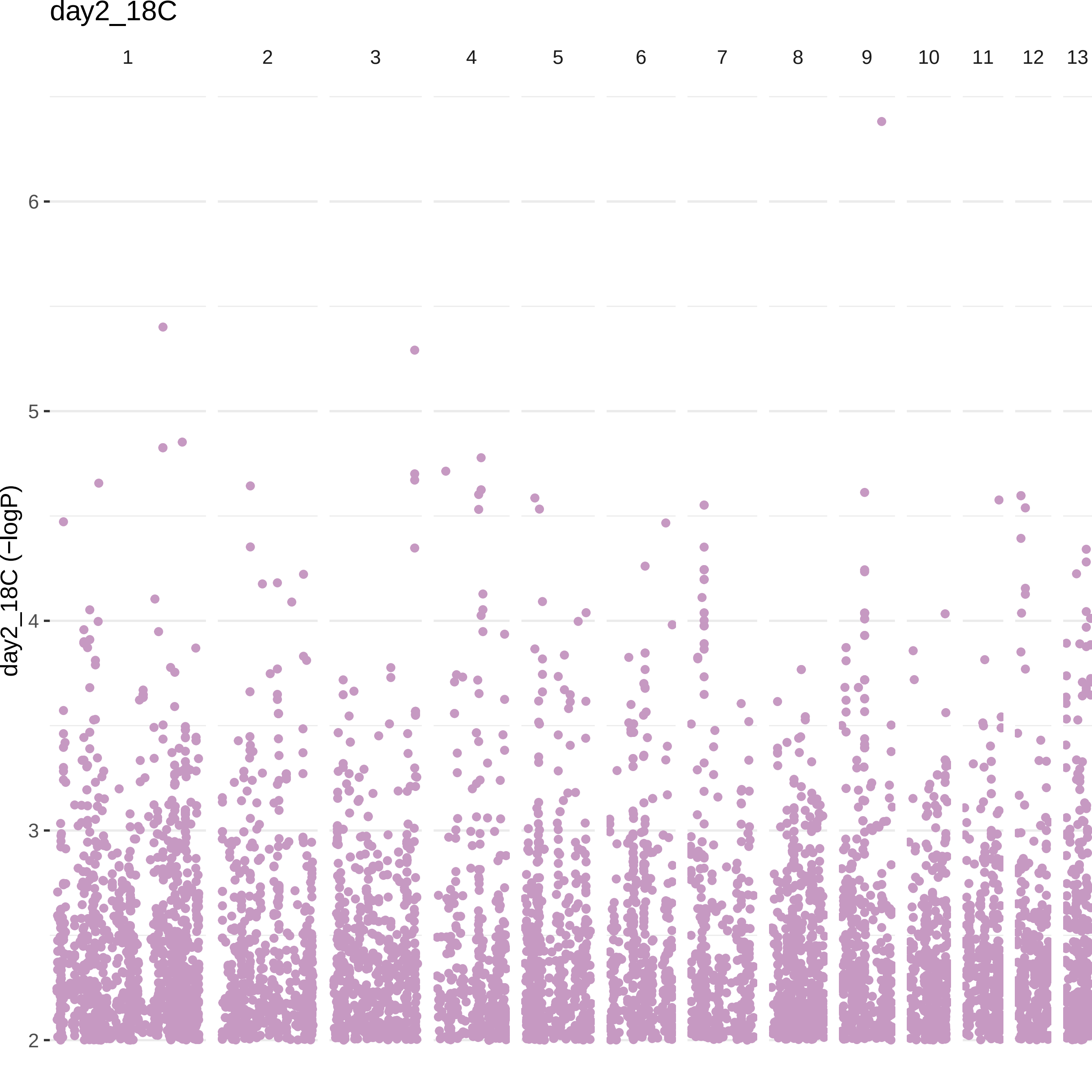
day2_18C
1
2
3
4
5
6
7
8
9
10
11
12
13
day2_18C (−logP)
6
5
4
3
2

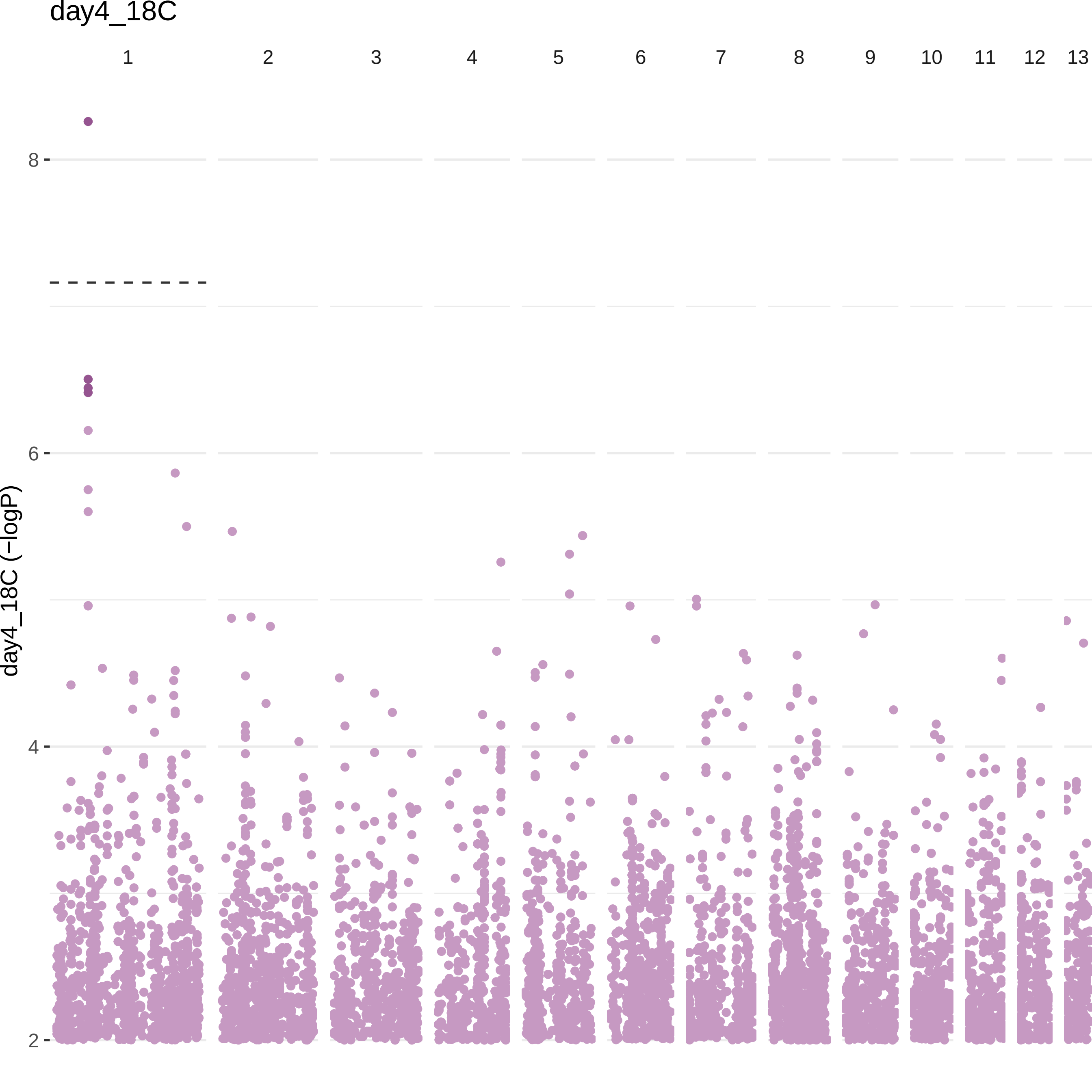

day4_18C
1
2
3
4
5
6
7
8
9
10
11
12
13
day4_18C (−logP)
8
6
4
2

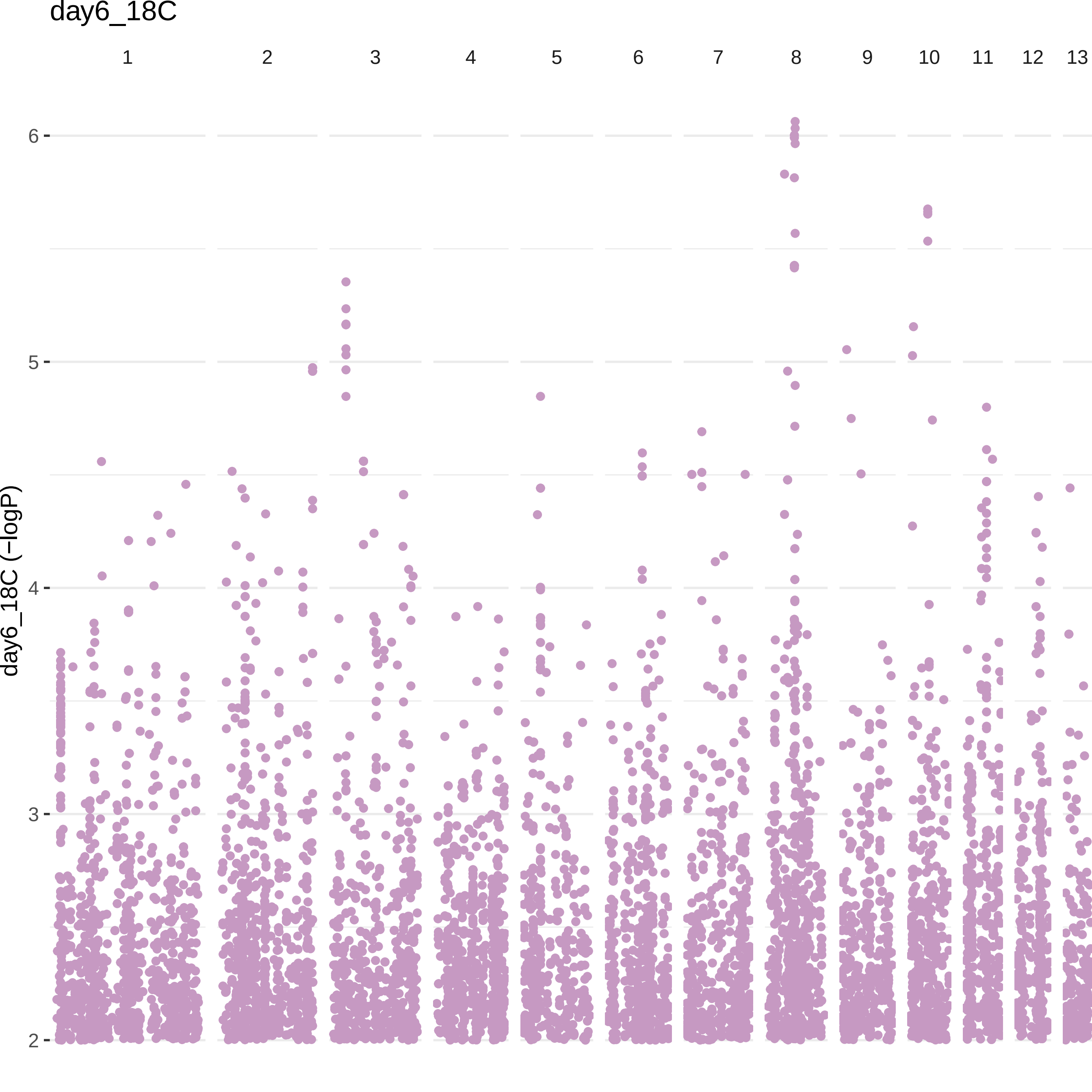

day6_18C
1
2
3
4
5
6
7
8
9
10
11
12
13
day6_18C (−logP)
6
5
4
3
2

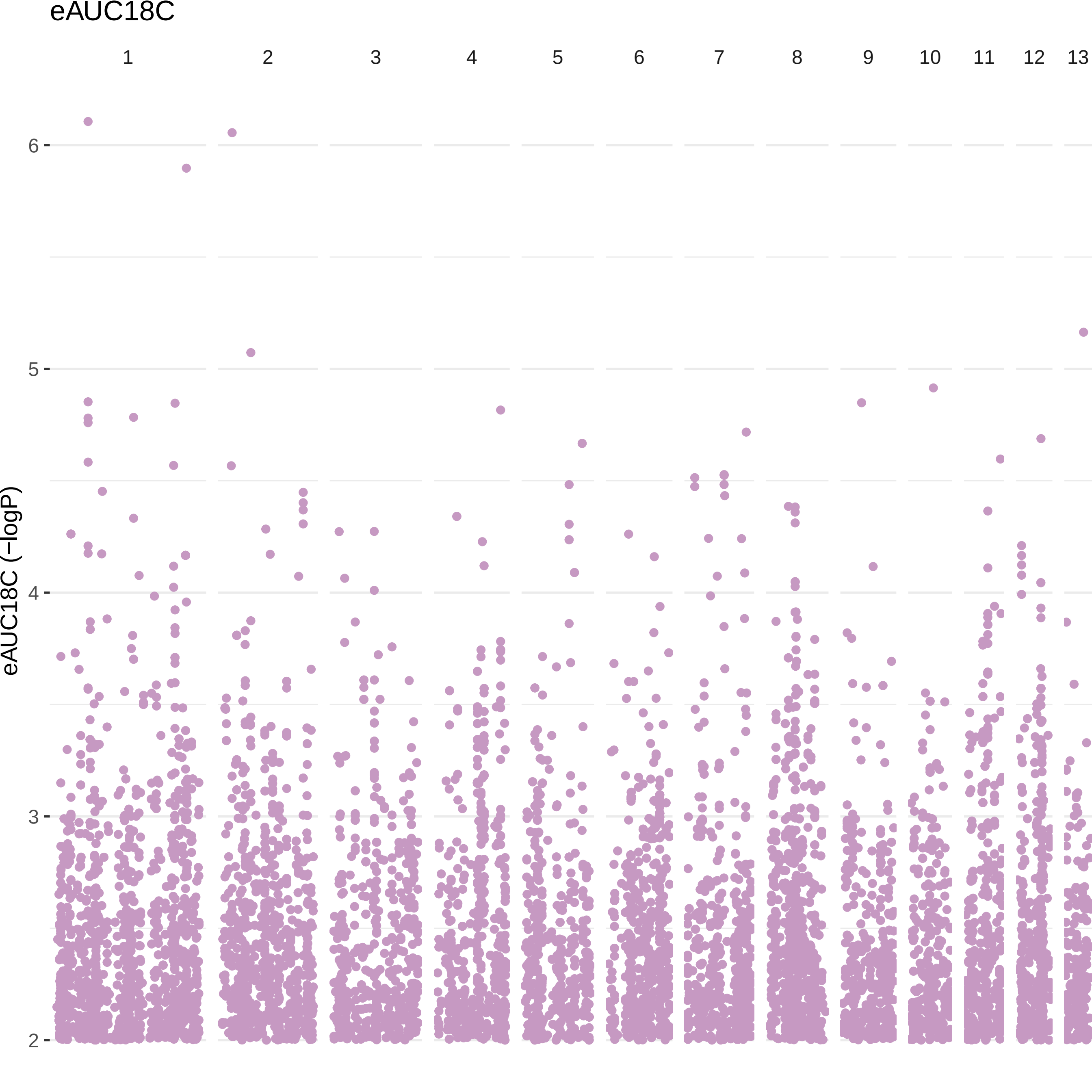
eAUC18C
1
2
3
4
5
6
7
8
9
10
11
12
13
6
5
4
3
2
eAUC18C (−logP)

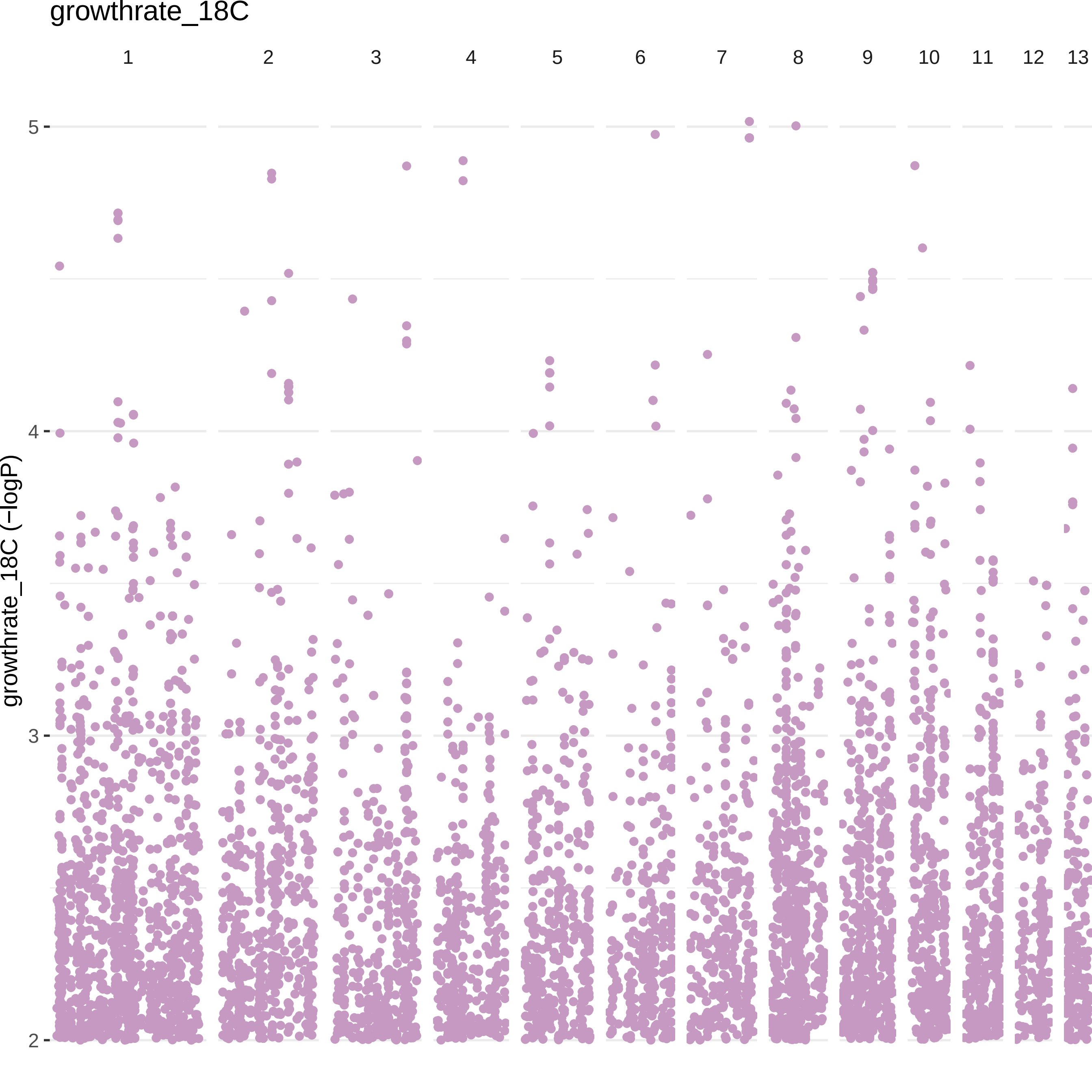

growthrate_18C
1
2
3
4
5
6
7
8
9
10
11
12
13
growthrate_18C (−logP)
5
4
3
2

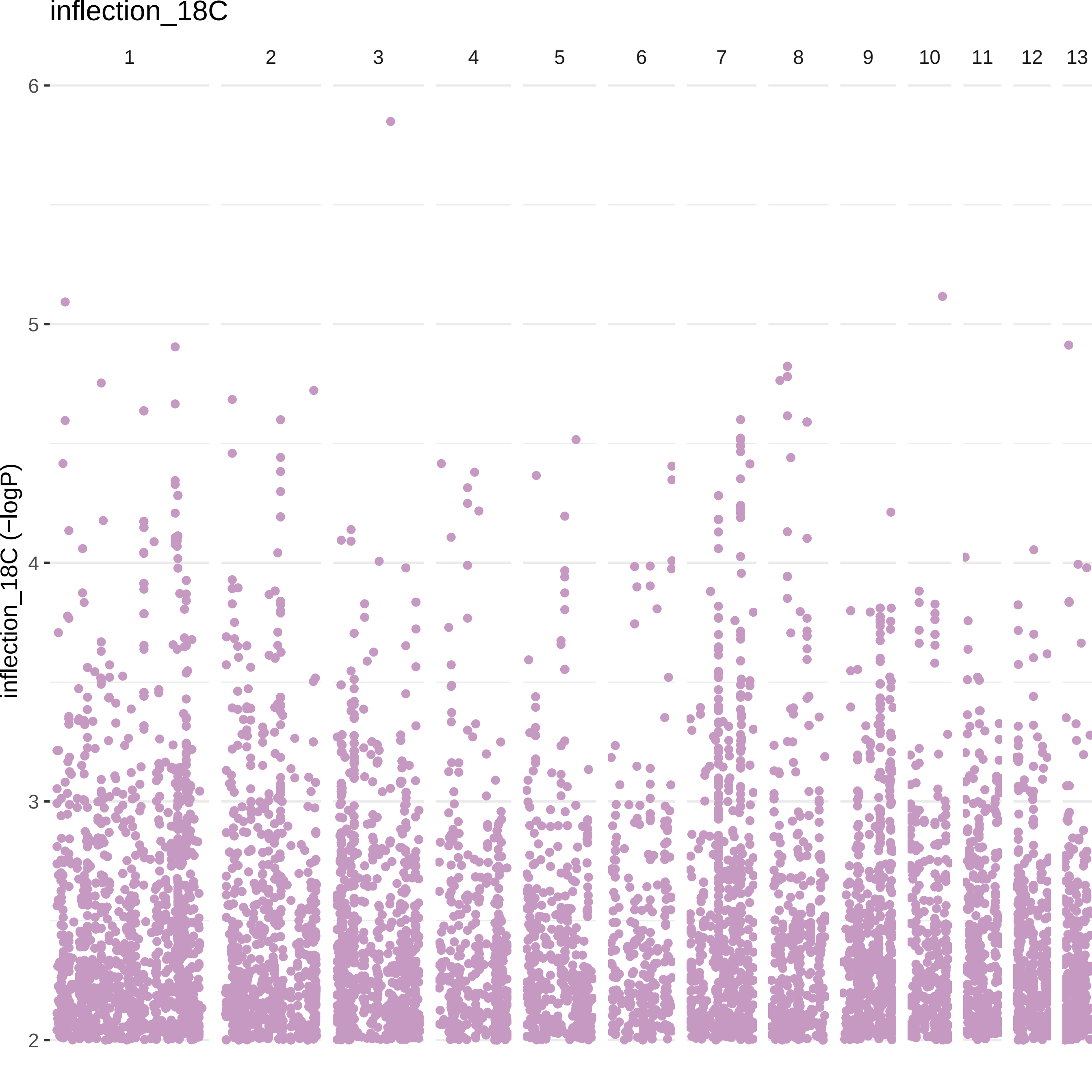

inflection_18C
inflection_18C (−logP)
1
2
3
4
5
6
7
8
9
10
11
12
13
6
5
4
3
2

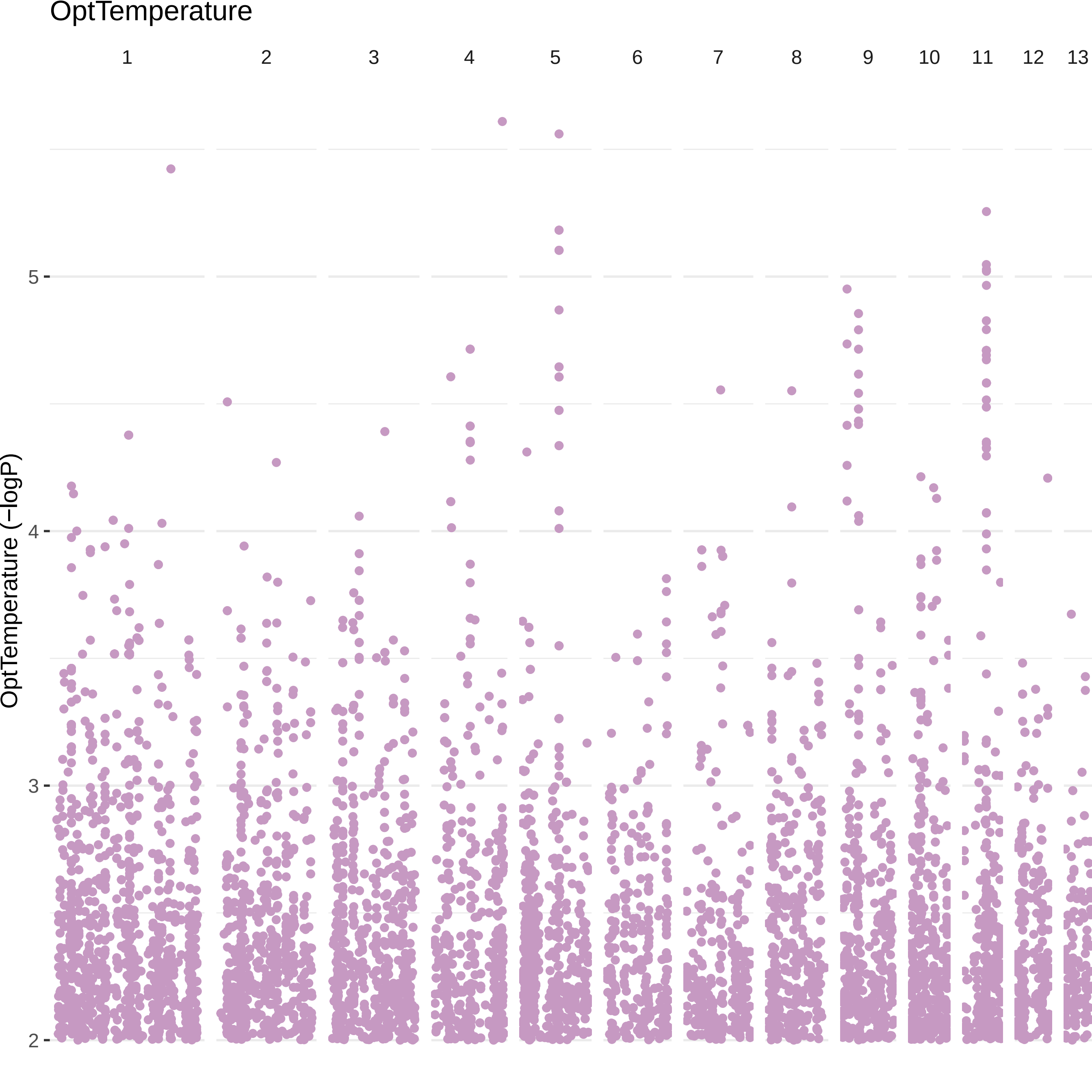
OptTemperature
1
2
3
4
5
6
7
8
9
10
11
12
13
OptTemperature (−logP)
5
4
3
2

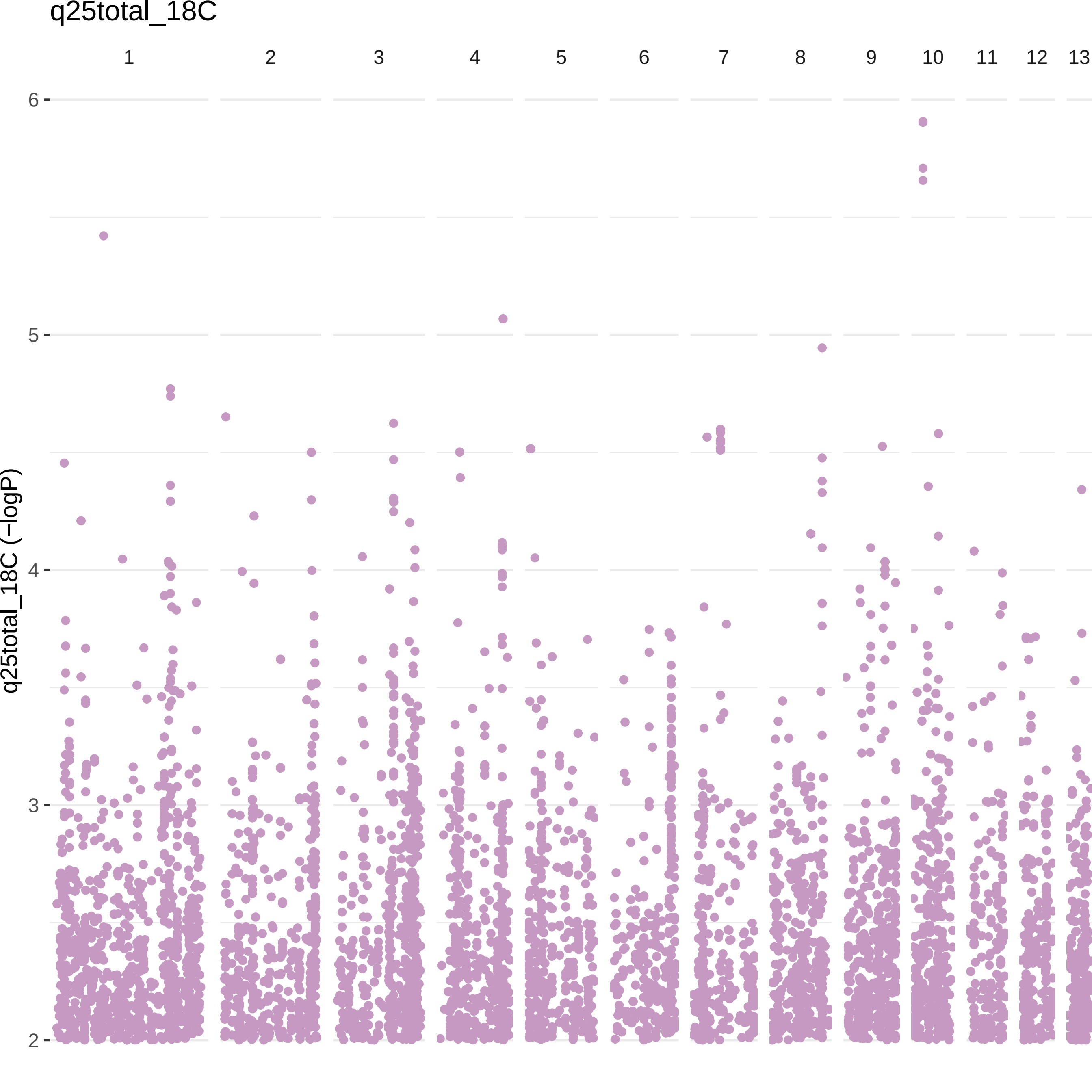
q25total_18C
1
2
3
4
5
6
7
8
9
10
11
12
13
6
5
4
3
2
q25total_18C (−logP)

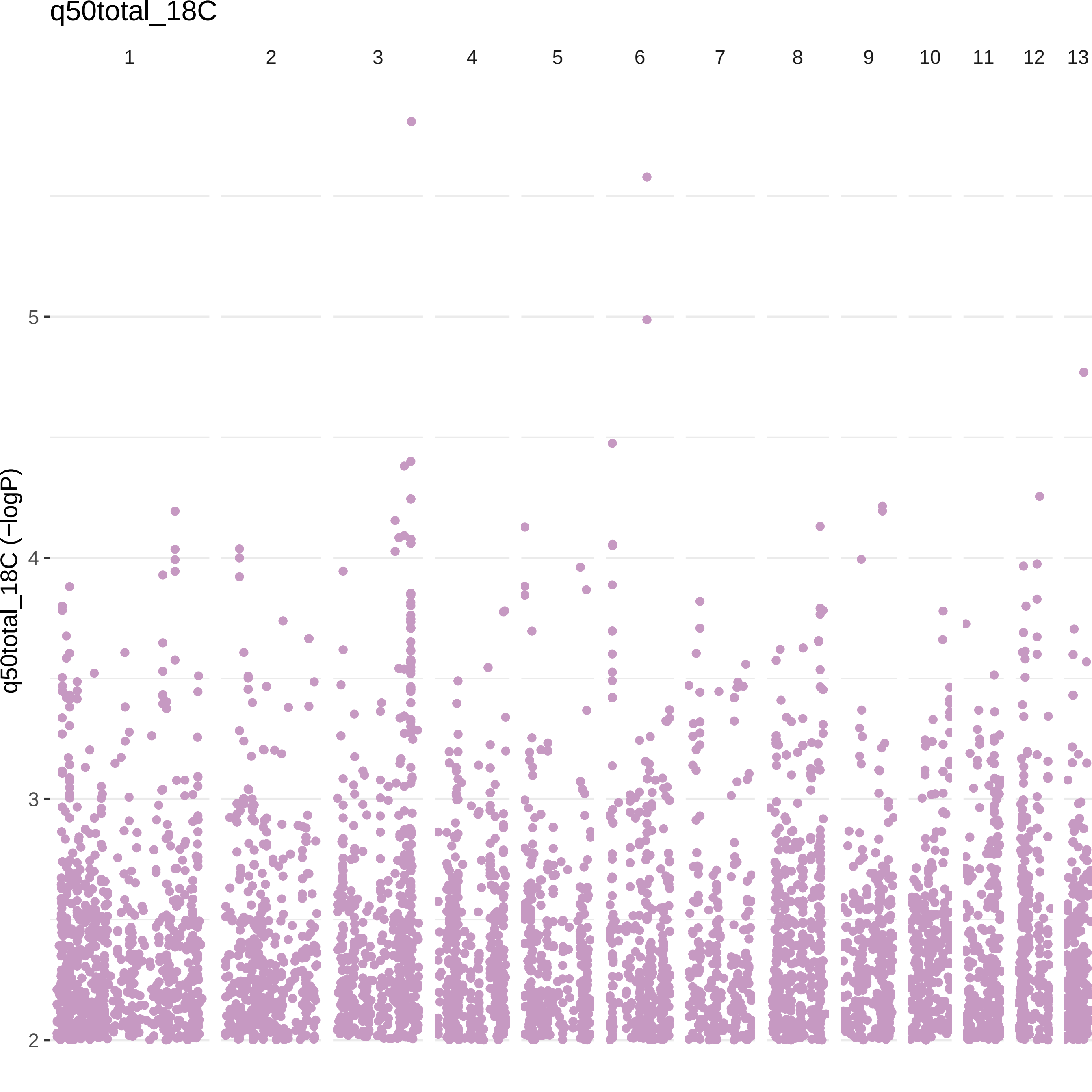

q50total_18C
1
2
3
4
5
6
7
8
9
10
11
12
13
q50total_18C (−logP)
5
4
3
2

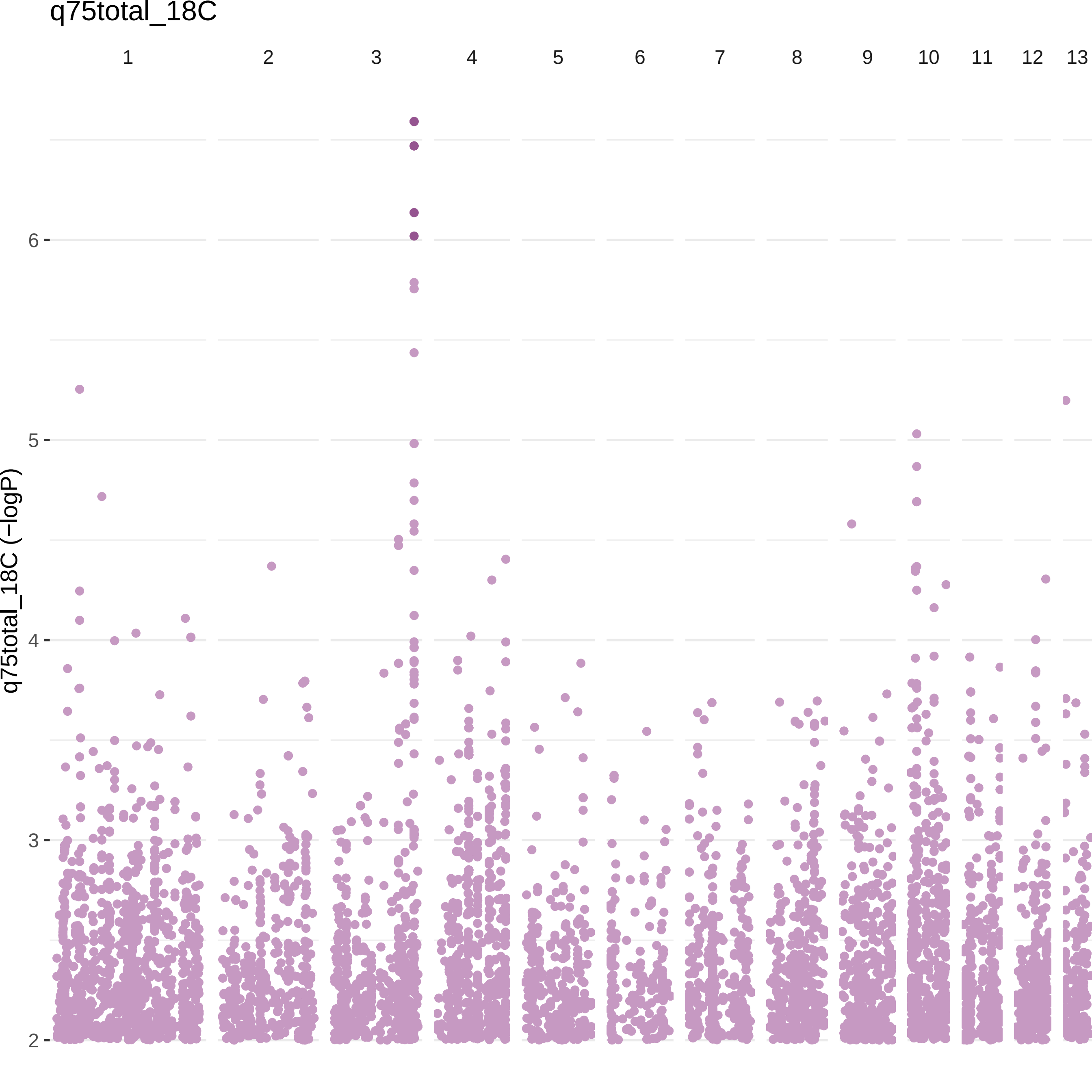

q75total_18C
1
2
3
4
5
6
7
8
9
10
11
12
13
q75total_18C (−logP)
6
5
4
3
2

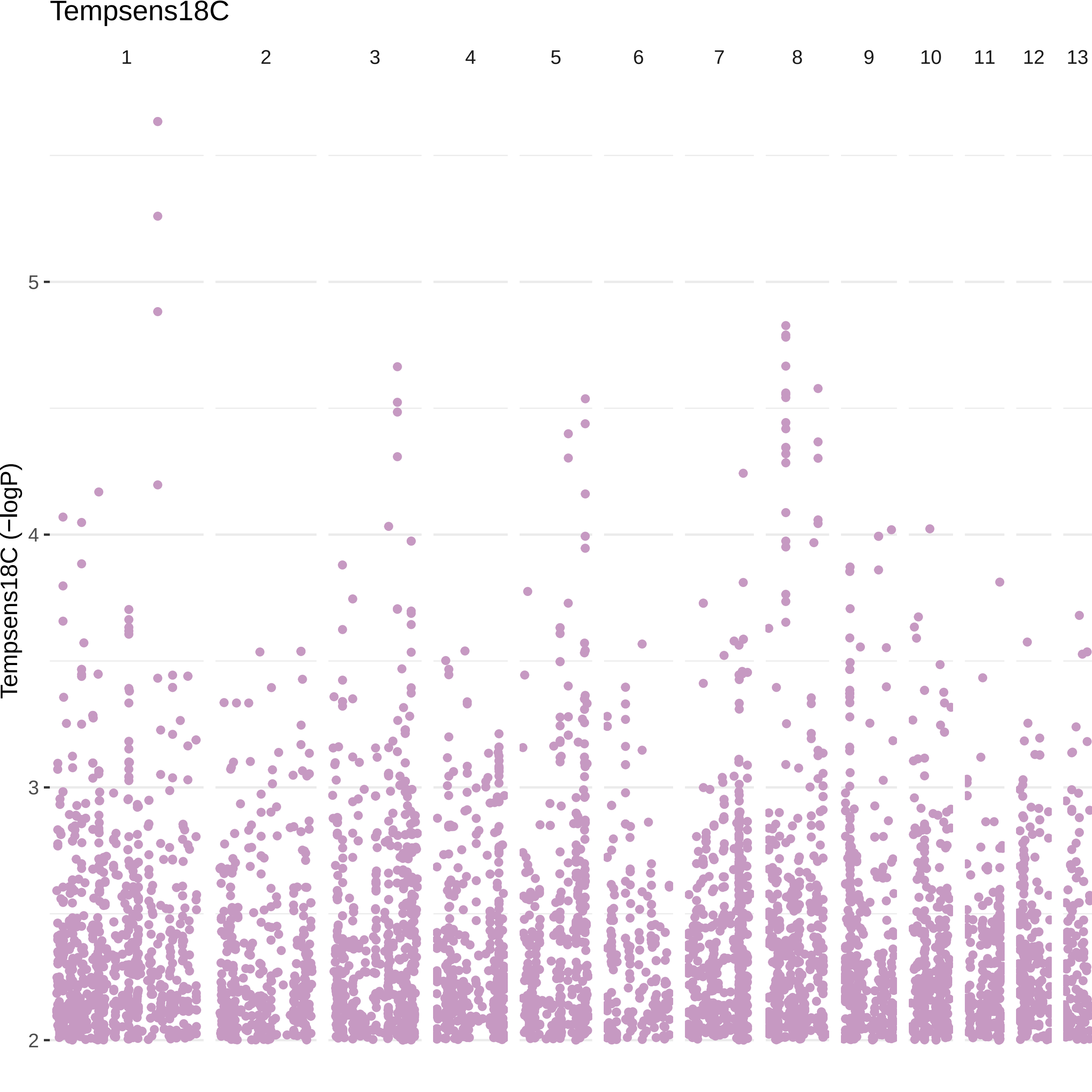

Tempsens18C
1
2
3
4
5
6
7
8
9
10
11
12
13
Tempsens18C (−logP)
5
4
3
2

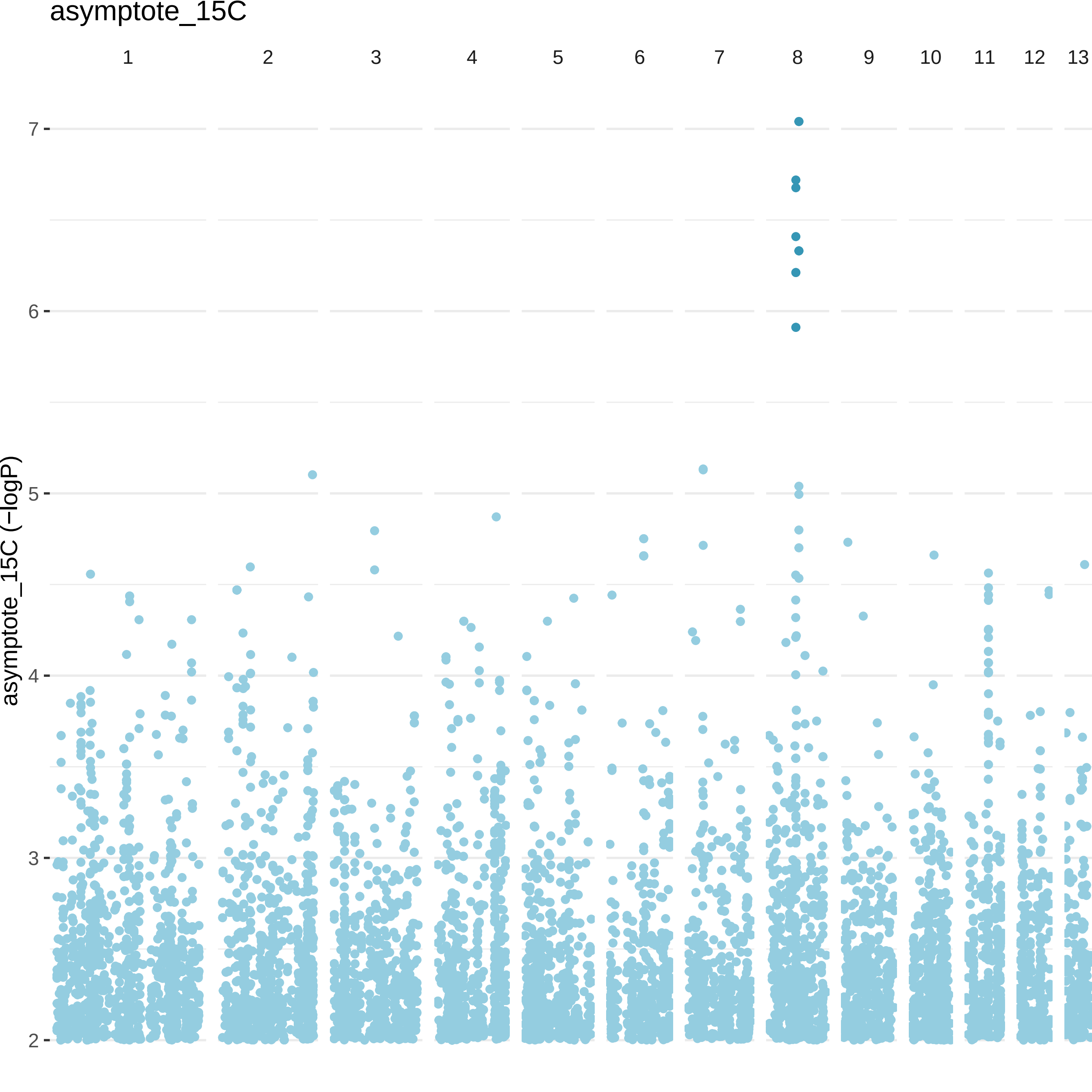

asymptote_15C
1
2
3
4
5
6
7
8
9
10
11
12
13
asymptote_15C (−logP)
7
6
5
4
3
2

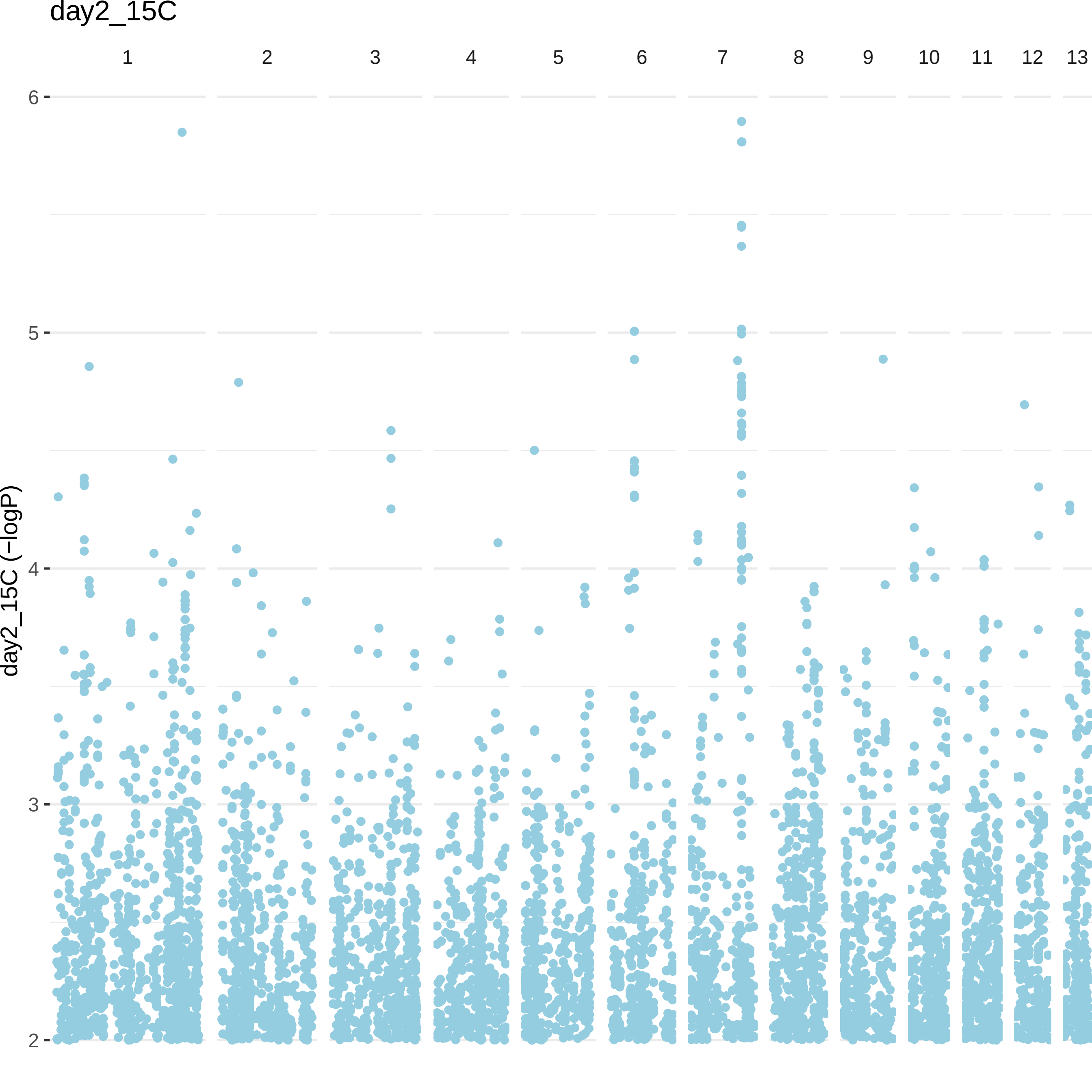
day2_15C
1
2
3
4
5
6
7
8
9
10
11
12
13
6
5
4
3
2
day2_15C (−logP)

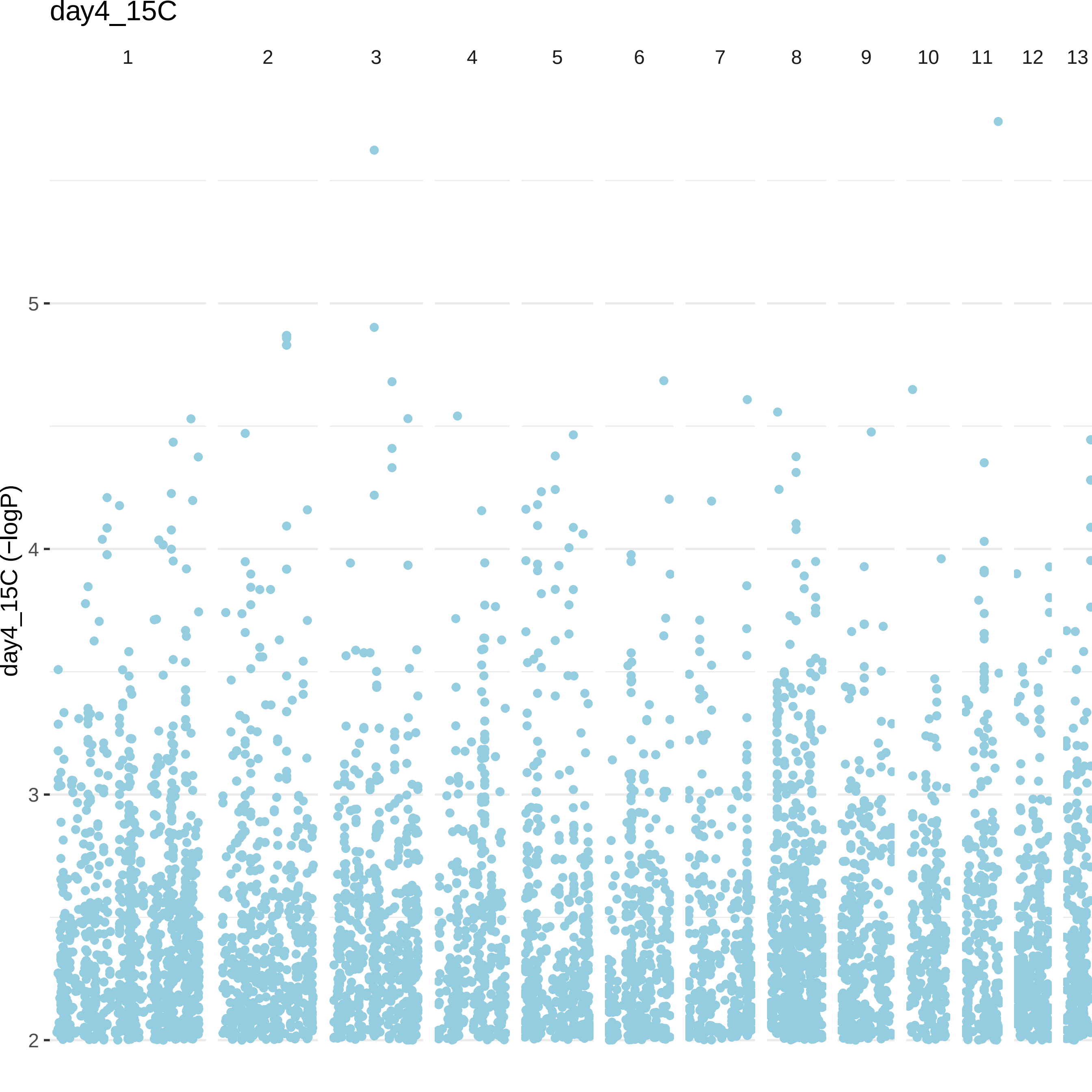

day4_15C
1
2
3
4
5
6
7
8
9
10
11
12
13
day4_15C (−logP)
5
4
3
2

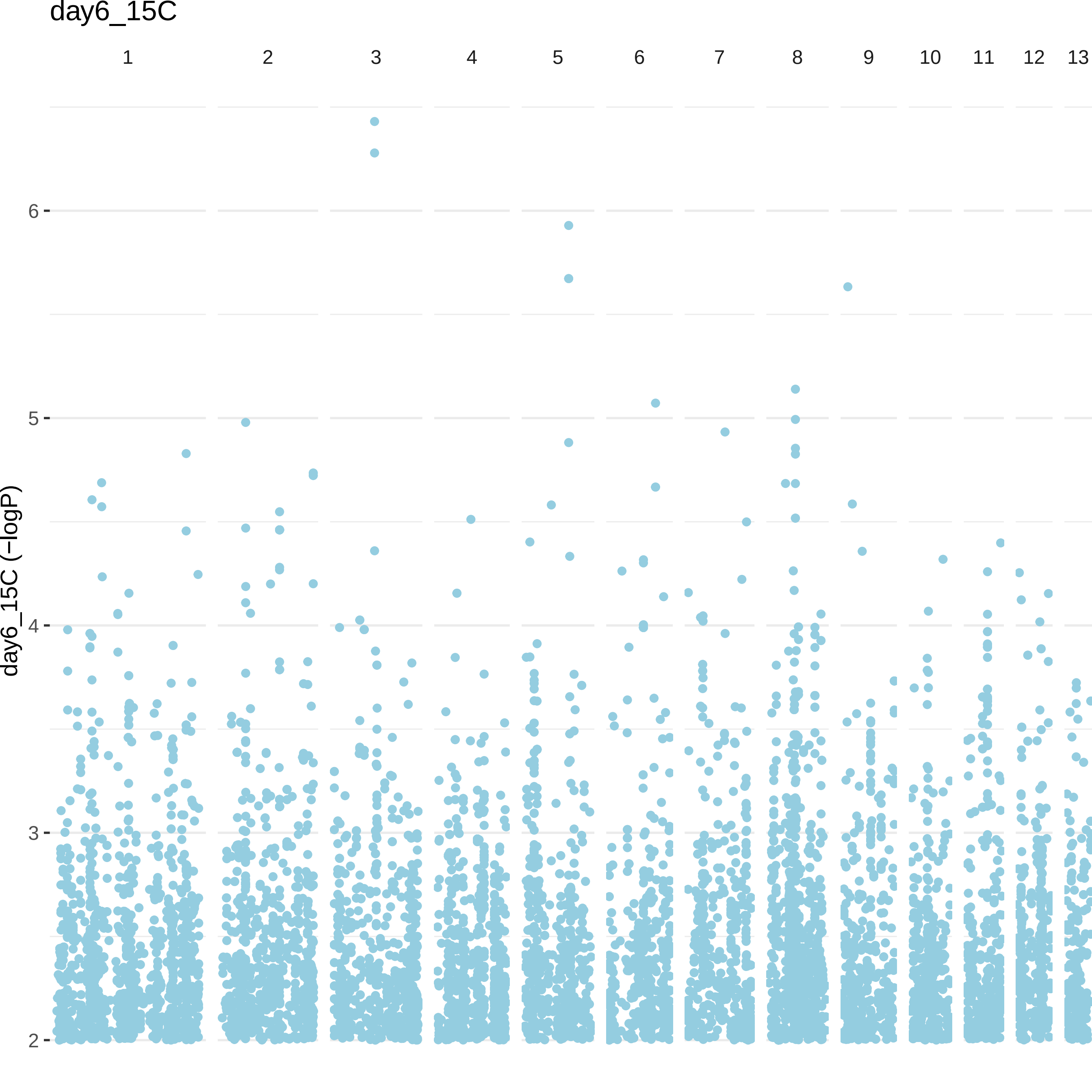
day6_15C
1
2
3
4
5
6
7
8
9
10
11
12
13
6
5
4
3
2
day6_15C (−logP)

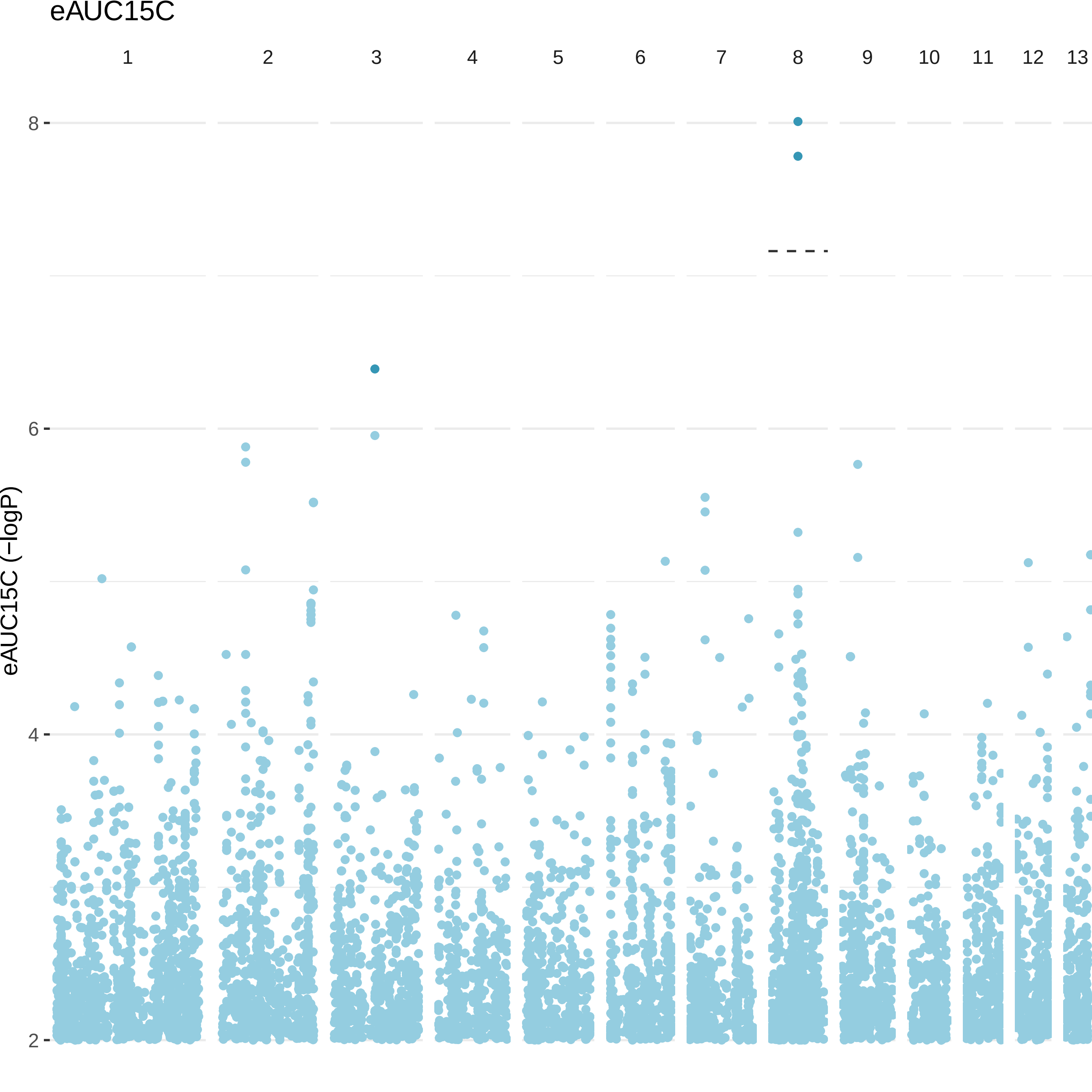

eAUC15C
1
2
3
4
5
6
7
8
9
10
11
12
13
8
6
4
2
eAUC15C (−logP)

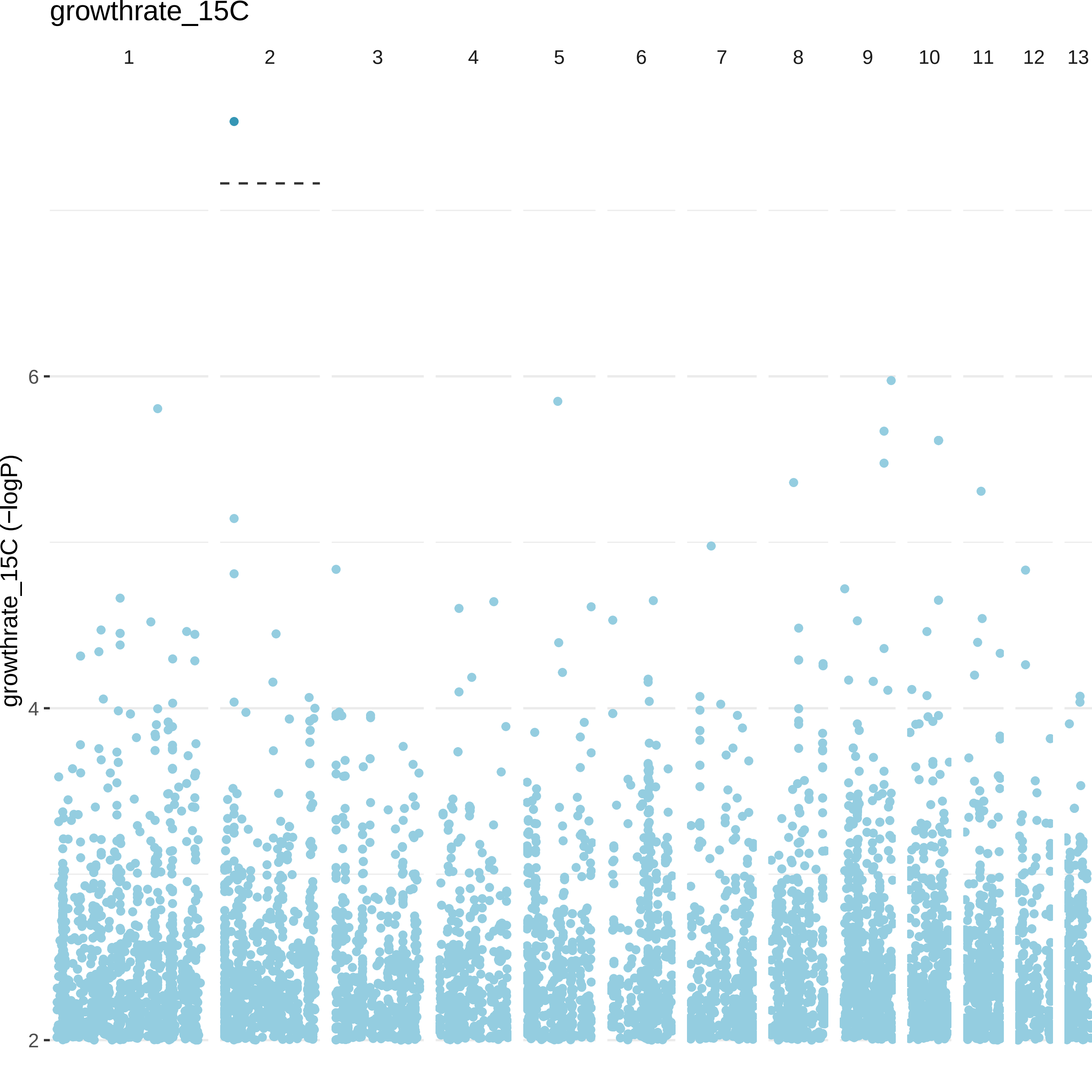

growthrate_15C
1
2
3
4
5
6
7
8
9
10
11
12
13
growthrate_15C (−logP)
6
4
2

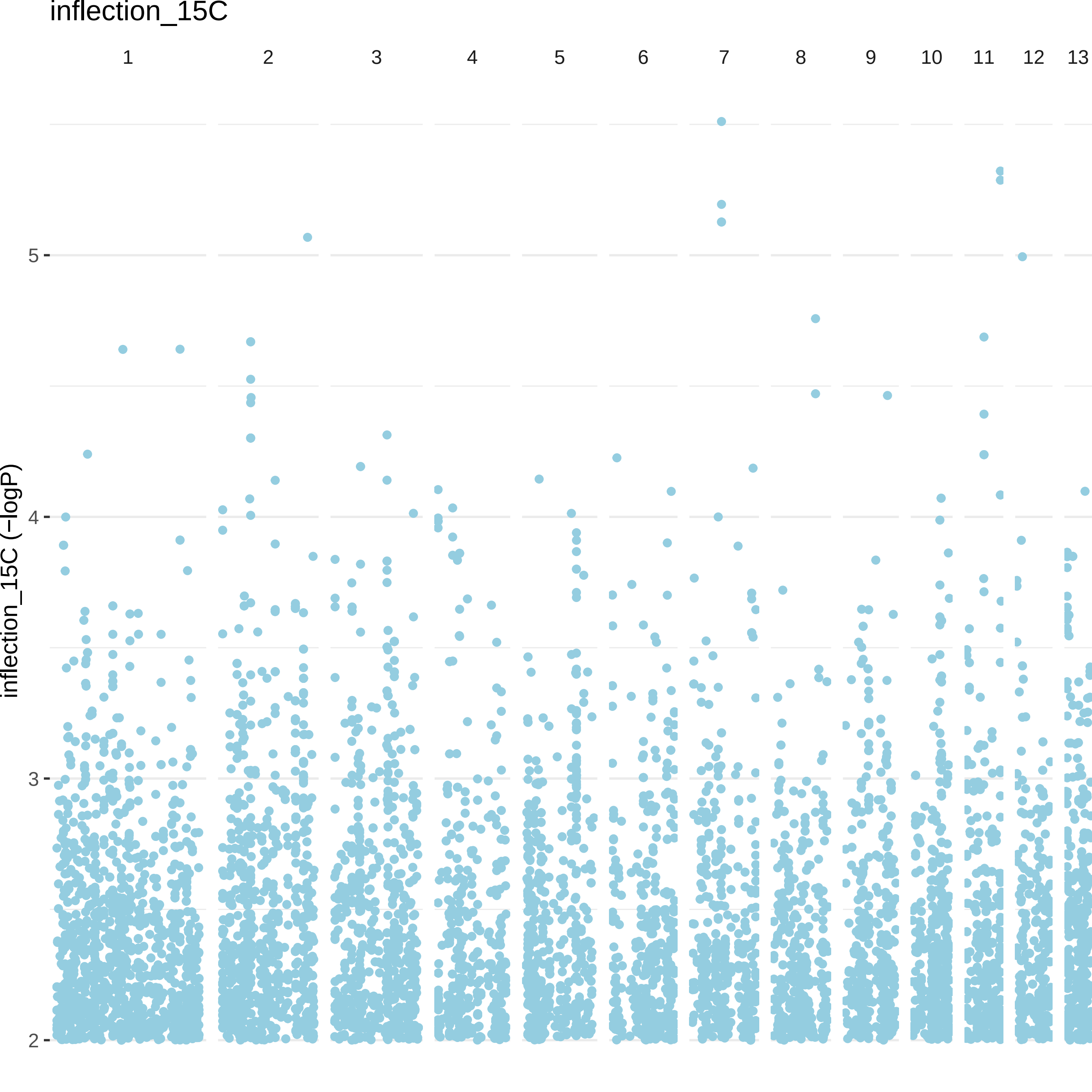

inflection_15C
1
2
3
4
5
6
7
8
9
10
11
12
13
inflection_15C (−logP)
5
4
3
2

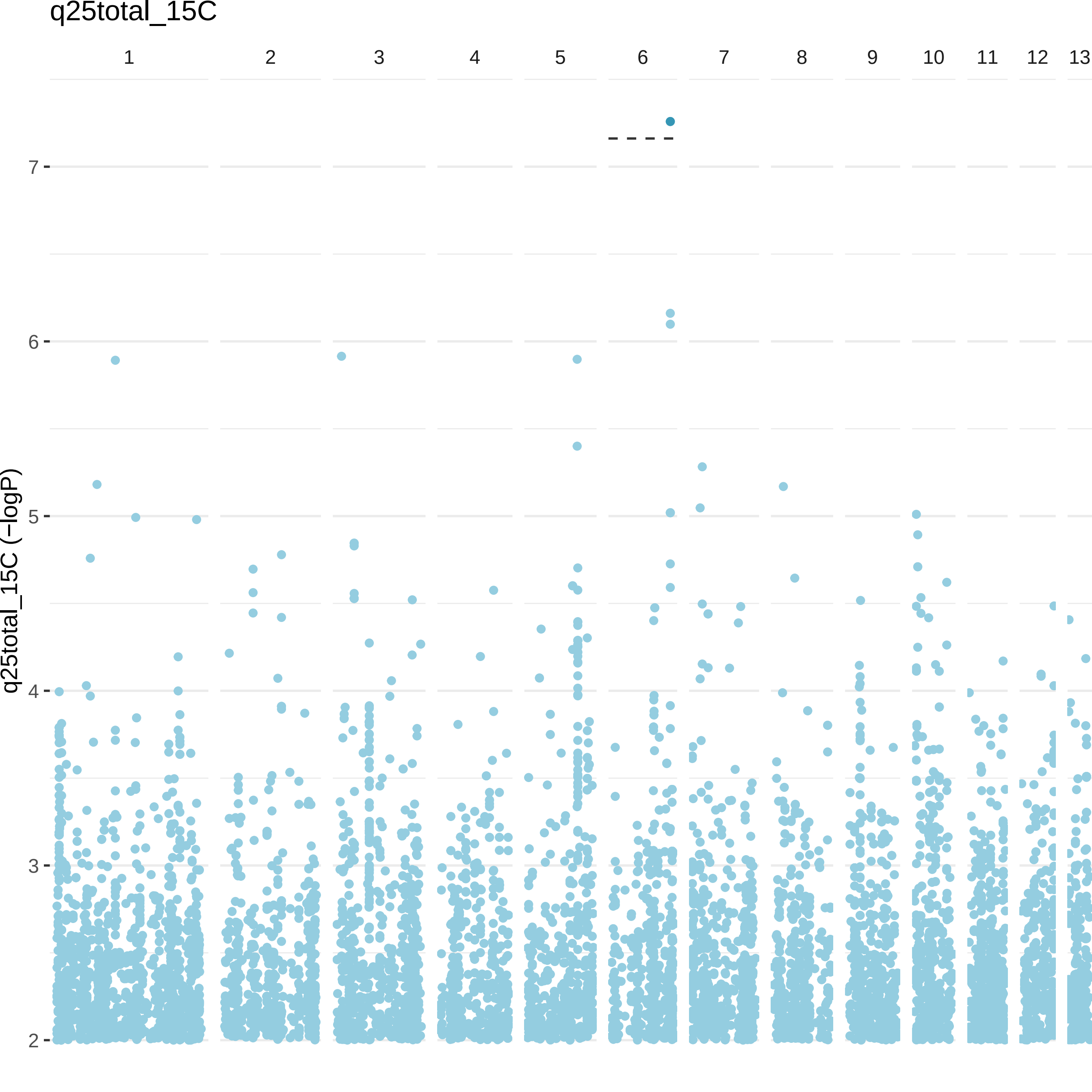
q25total_15C
1
2
3
4
5
6
7
8
9
10
11
12
13
q25total_15C (−logP)
7
6
5
4
3
2

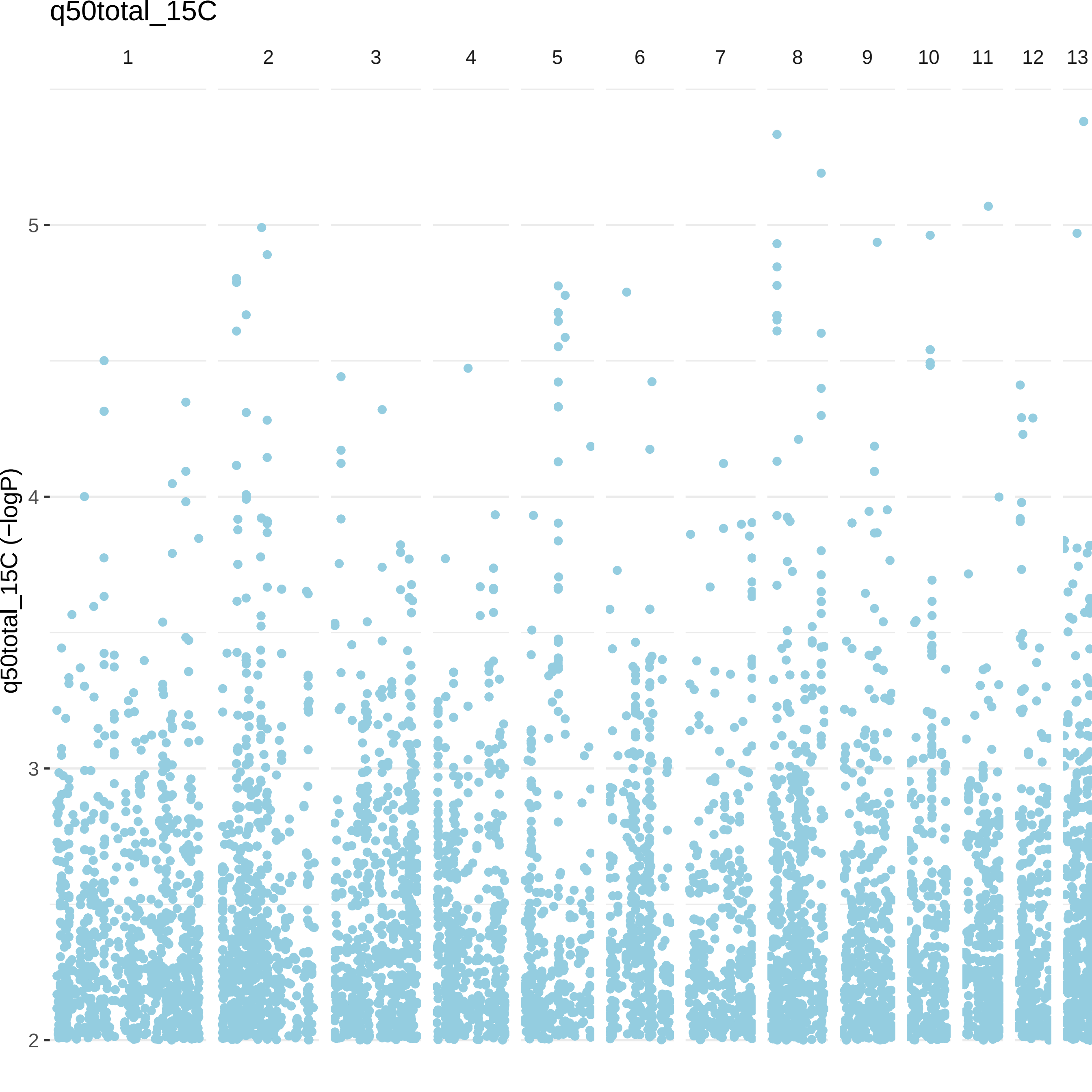

q50total_15C
1
2
3
4
5
6
7
8
9
10
11
12
13
q50total_15C (−logP)
5
4
3
2

q75total_15C

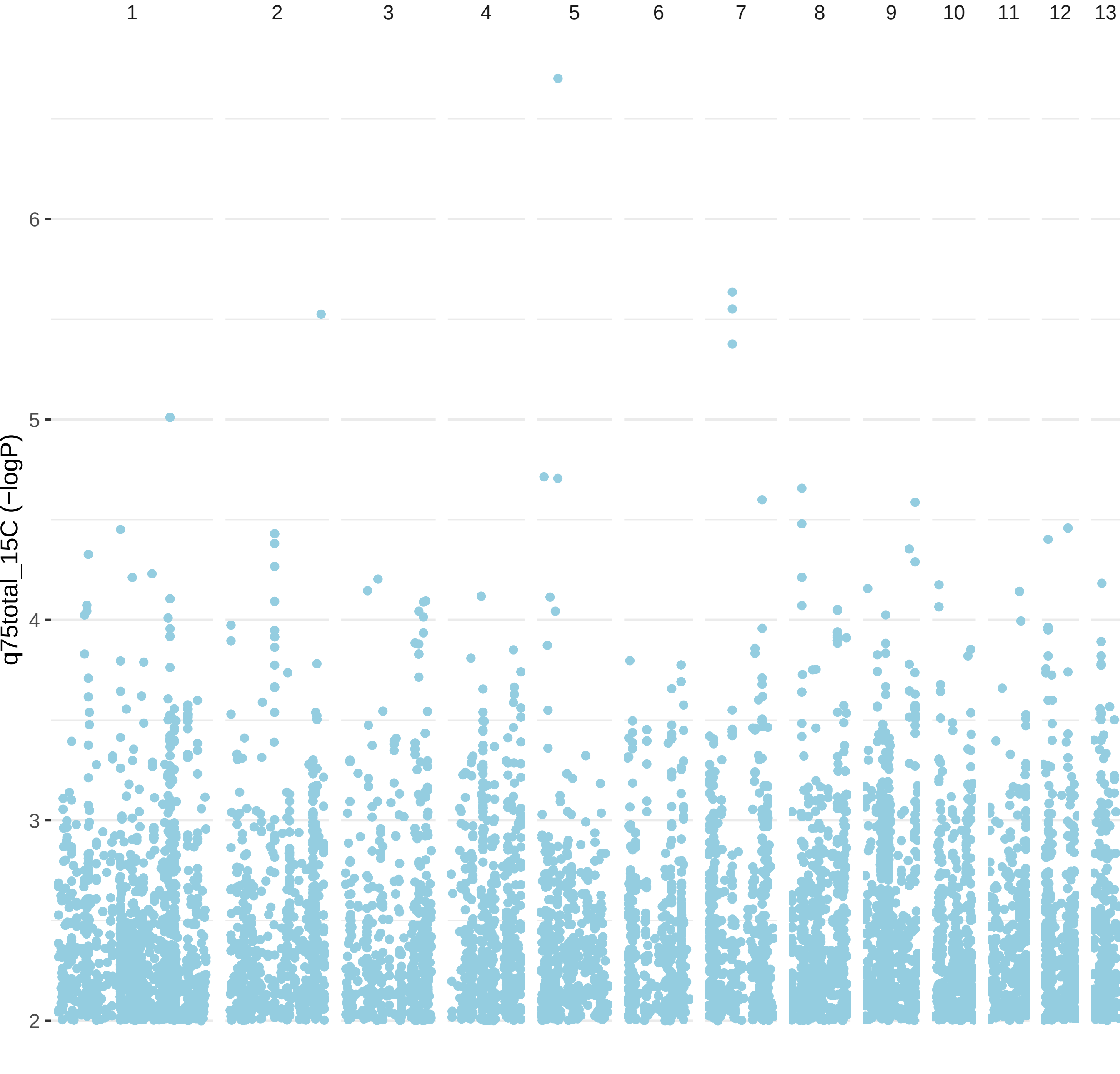

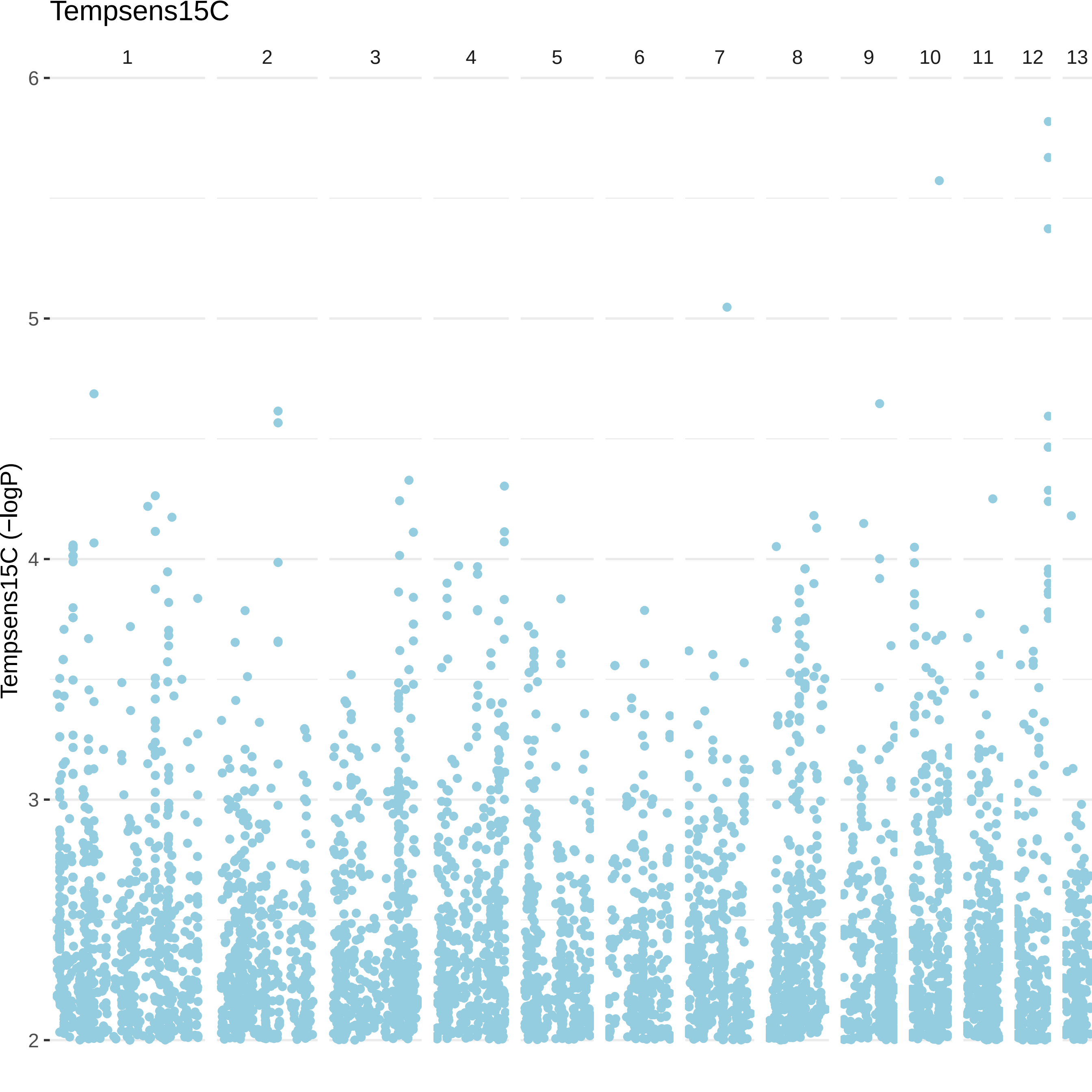
Tempsens15C
1
2
3
4
5
6
7
8
9
10
11
12
13
Tempsens15C (−logP)
6
5
4
3
2

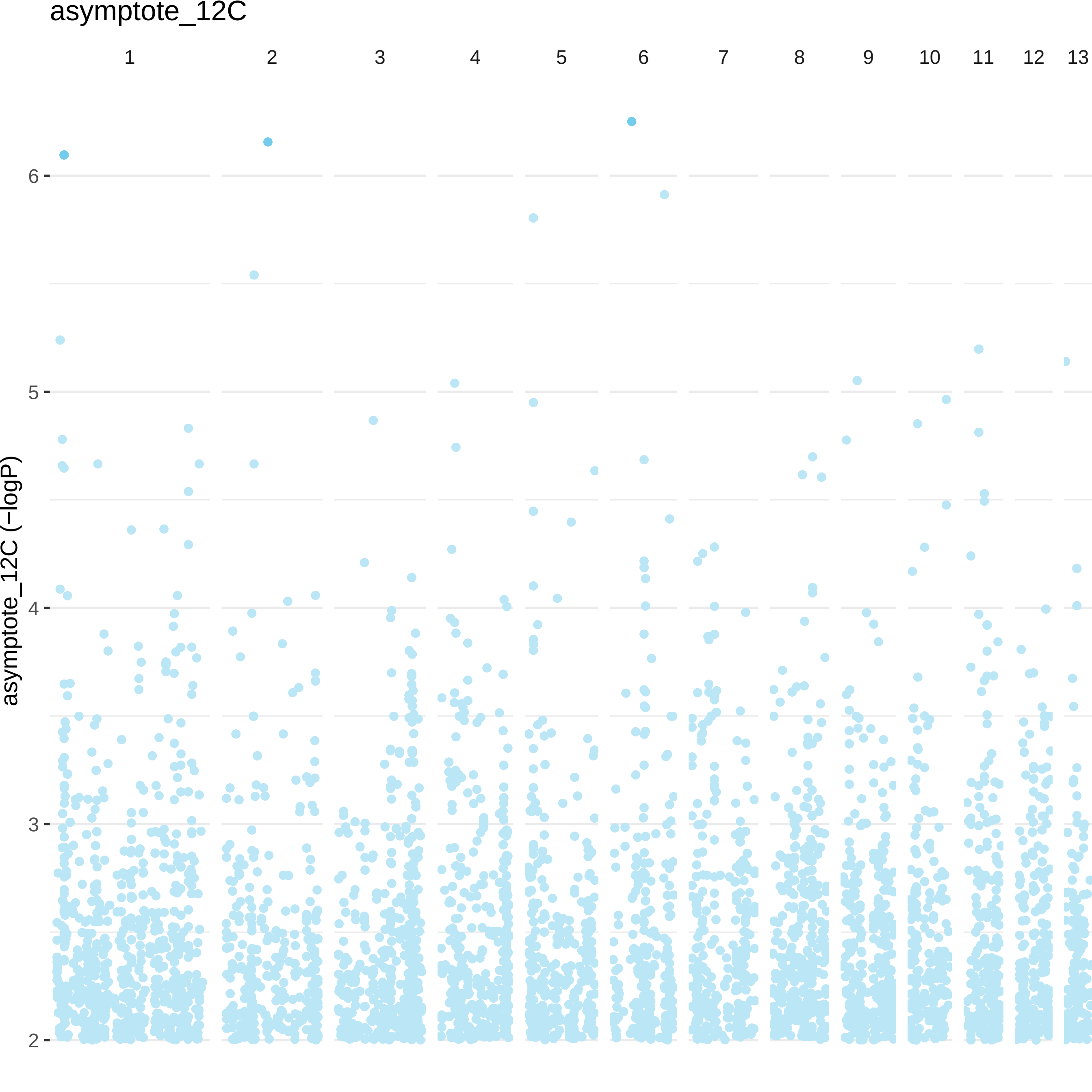

asymptote_12C
1
2
3
4
5
6
7
8
9
10
11
12
13
6
5
4
3
2
asymptote_12C (−logP)

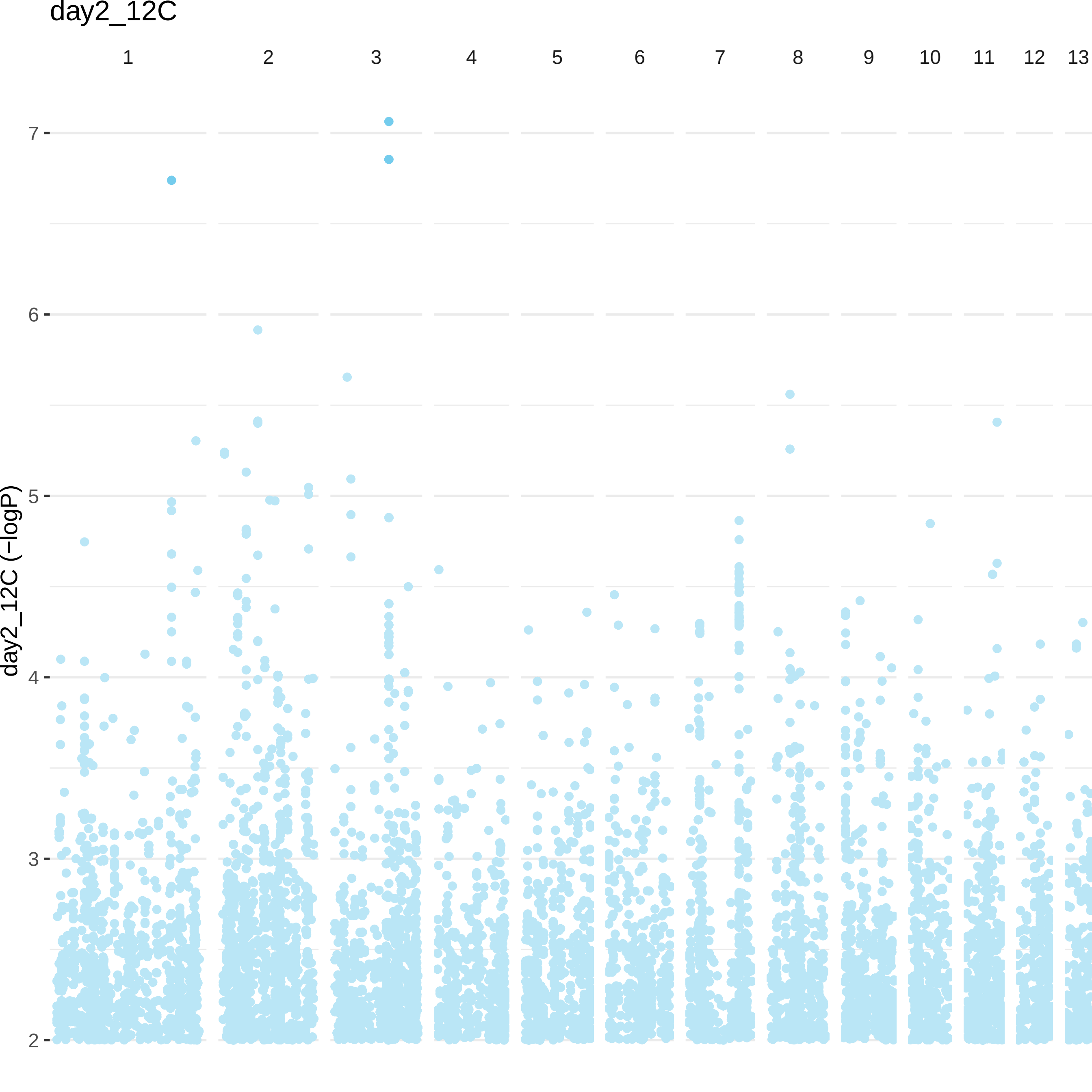

day2_12C
1
2
3
4
5
6
7
8
9
10
11
12
13
day2_12C (−logP)
7
6
5
4
3
2

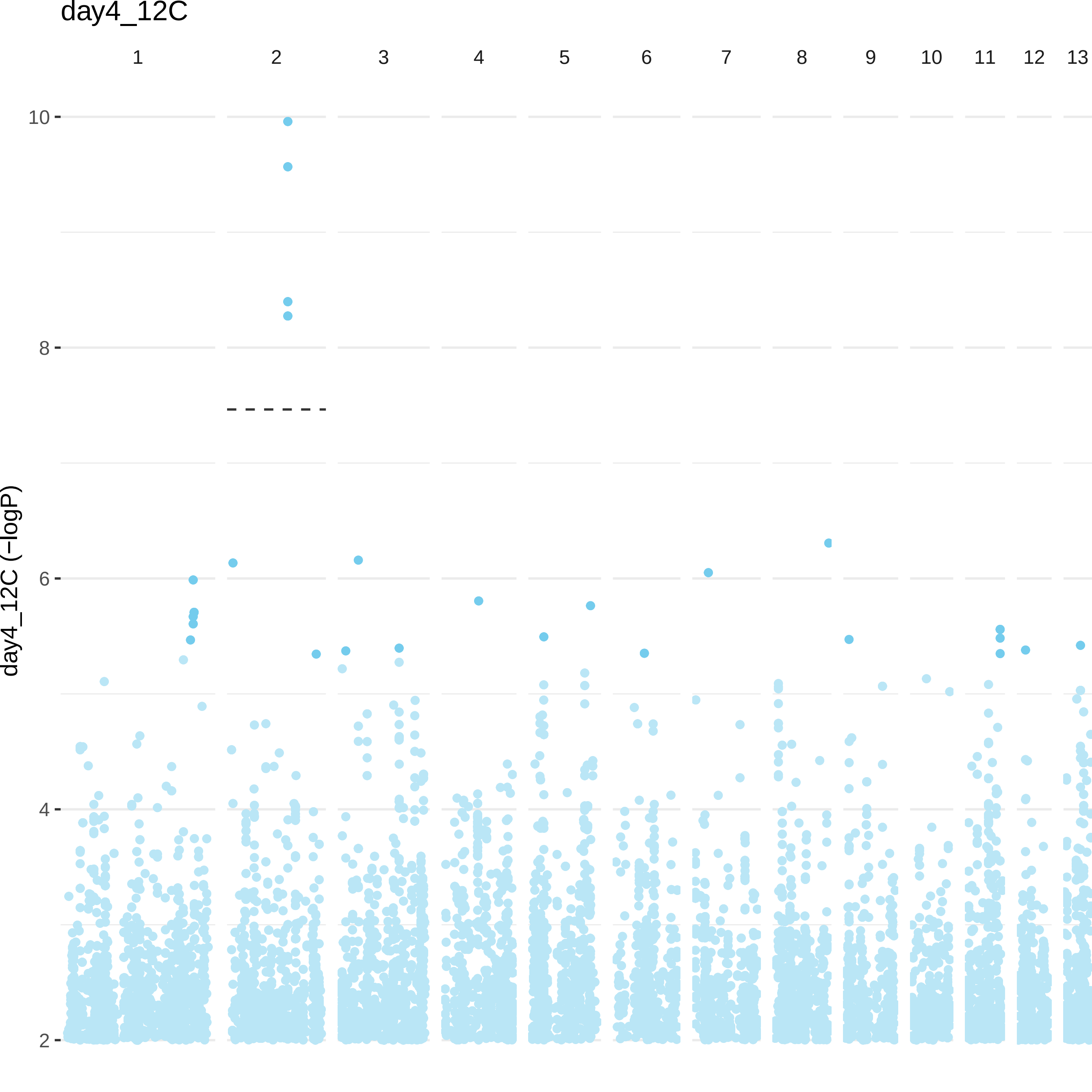

day4_12C
1
2
3
4
5
6
7
8
9
10
11
12
13
10
8
6
4
2
day4_12C (−logP)

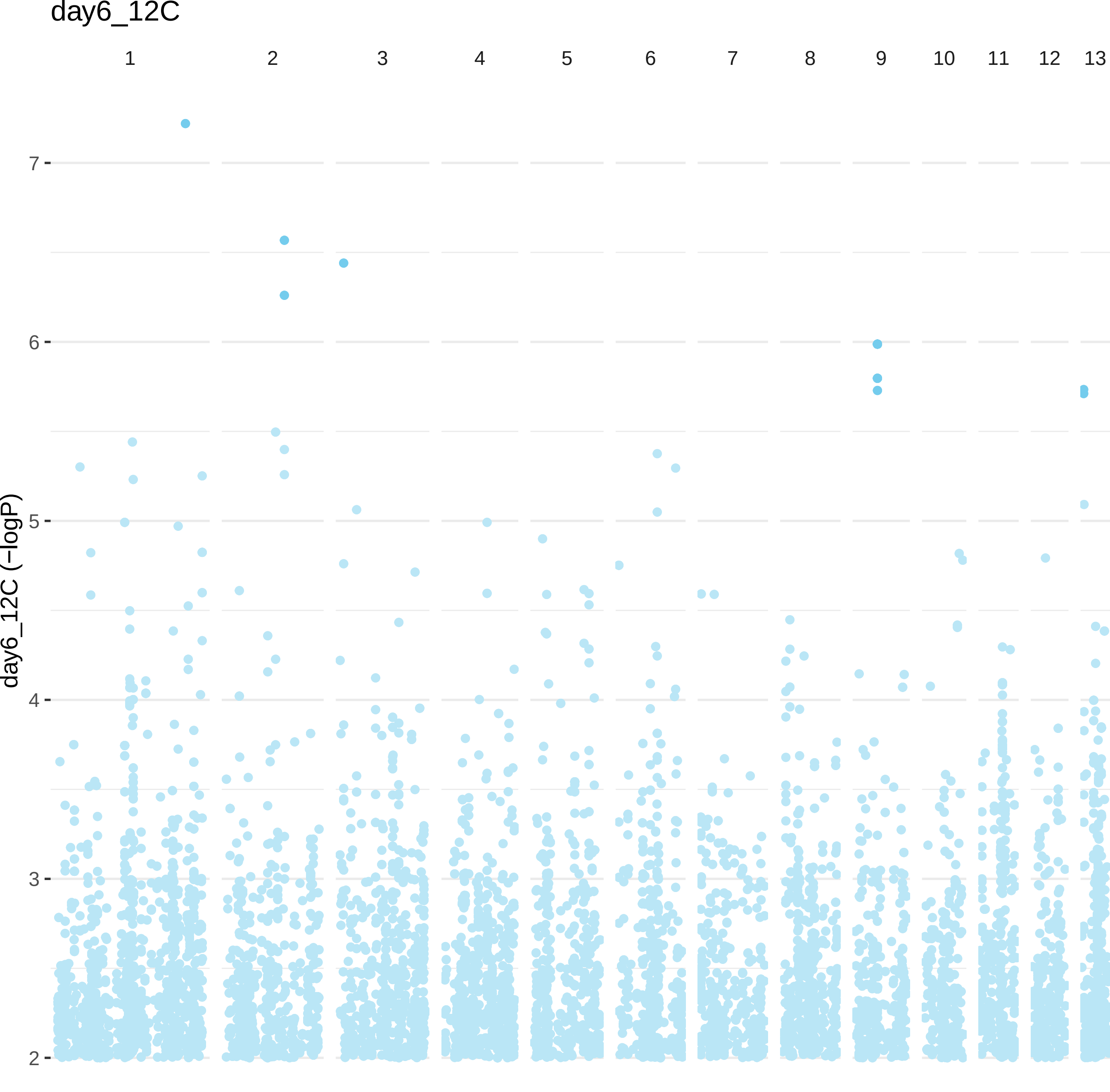
day6_12C
1
2
3
4
5
6
7
8
9
10
11
12
13
7
6
5
4
3
2
day6_12C (−logP)

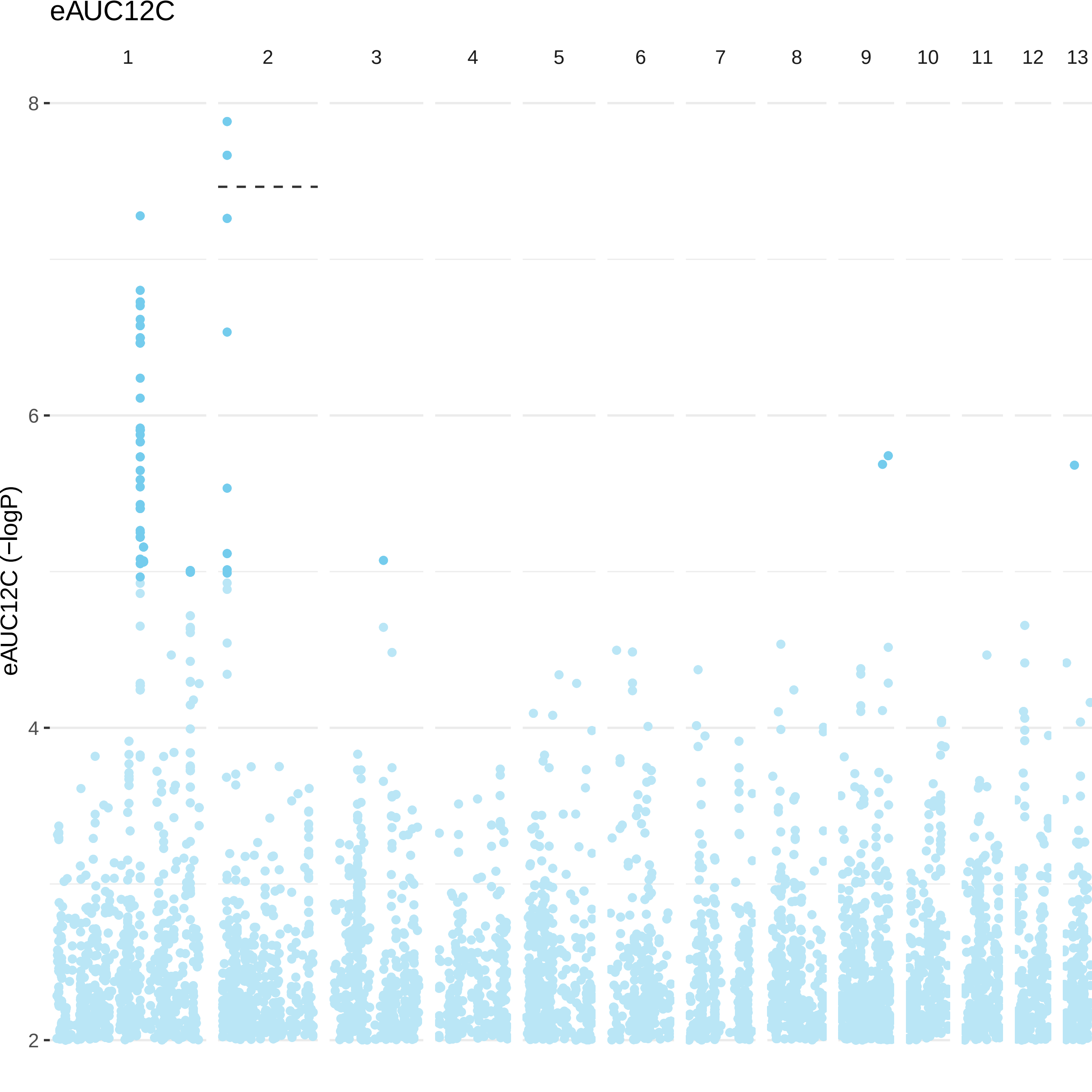
eAUC12C
1
2
3
4
5
6
7
8
9
10
11
12
13
8
6
4
2
eAUC12C (−logP)

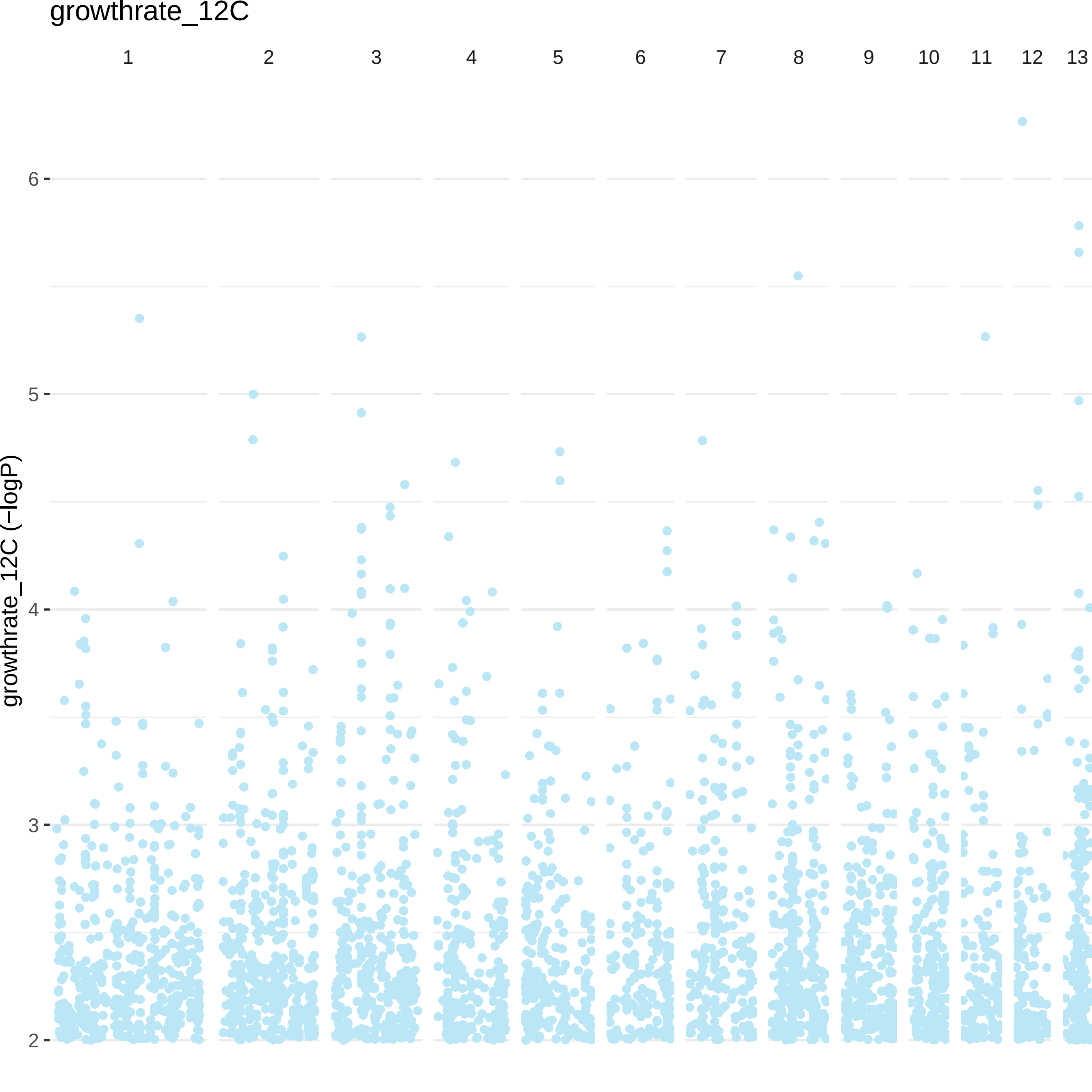

growthrate_12C
1
2
3
4
5
6
7
8
9
10
11
12
13
6
5
4
3
2
growthrate_12C (−logP)

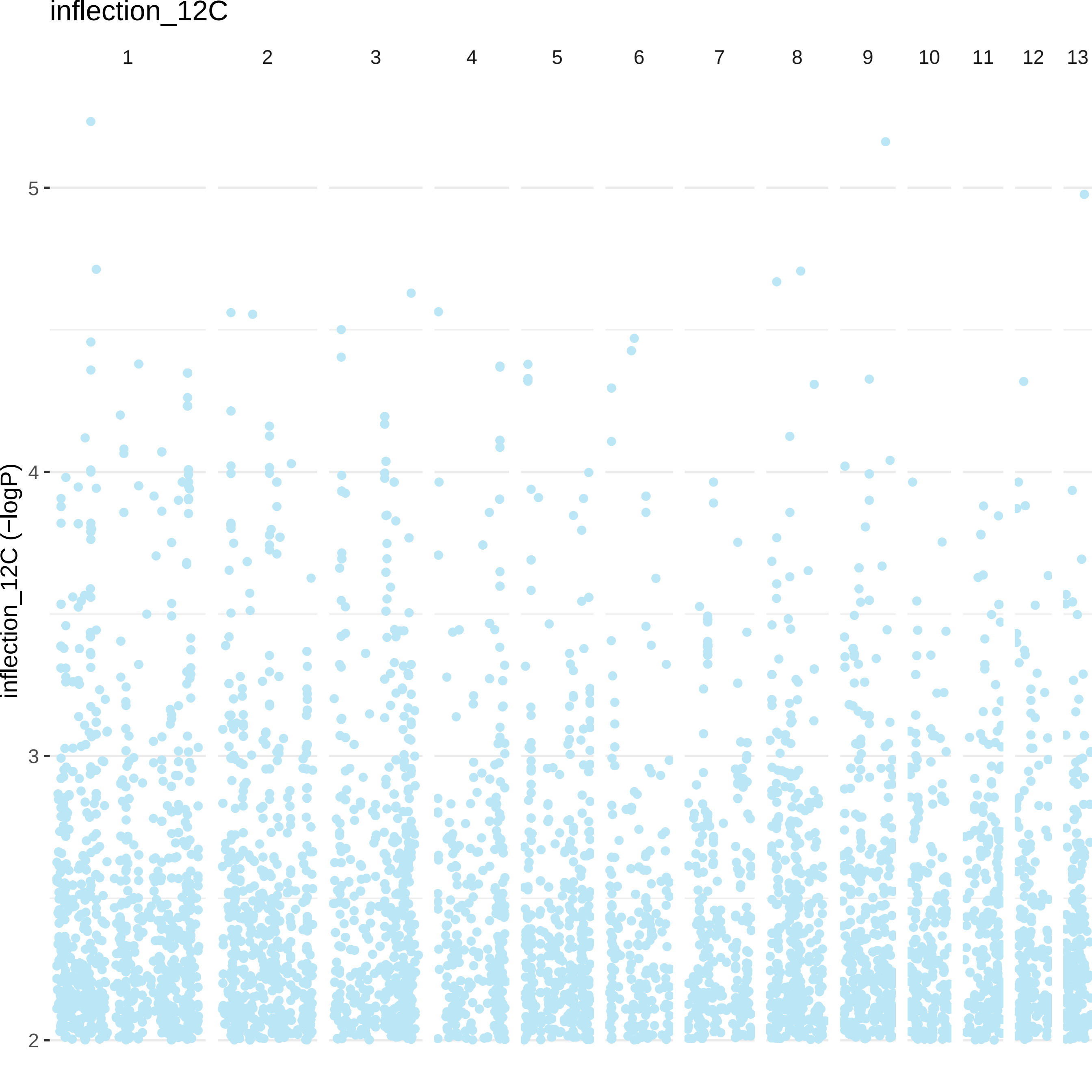

inflection_12C
1
2
3
4
5
6
7
8
9
10
11
12
13
inflection_12C (−logP)
5
4
3
2

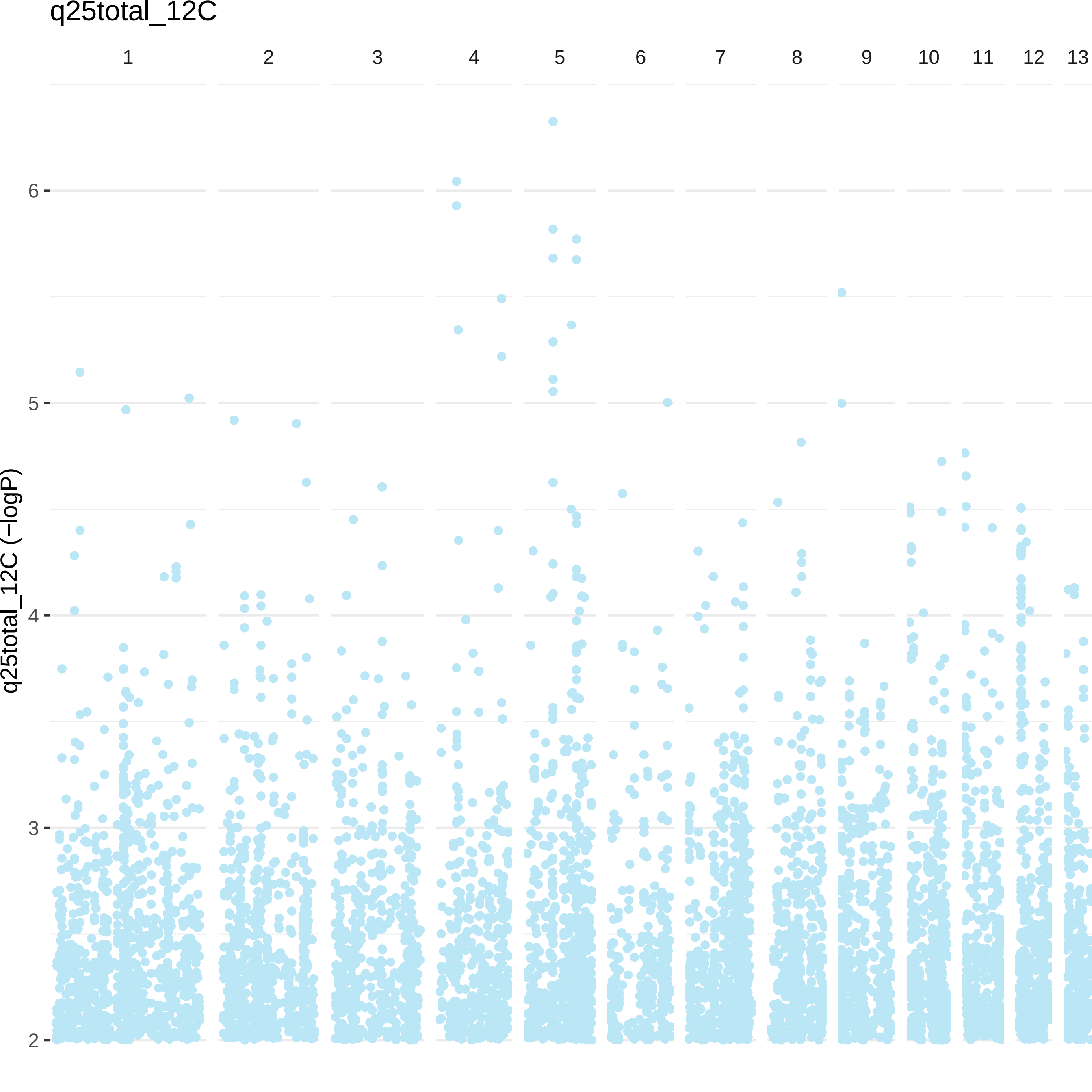

q25total_12C
1
2
3
4
5
6
7
8
9
10
11
12
13
q25total_12C (−logP)
6
5
4
3
2

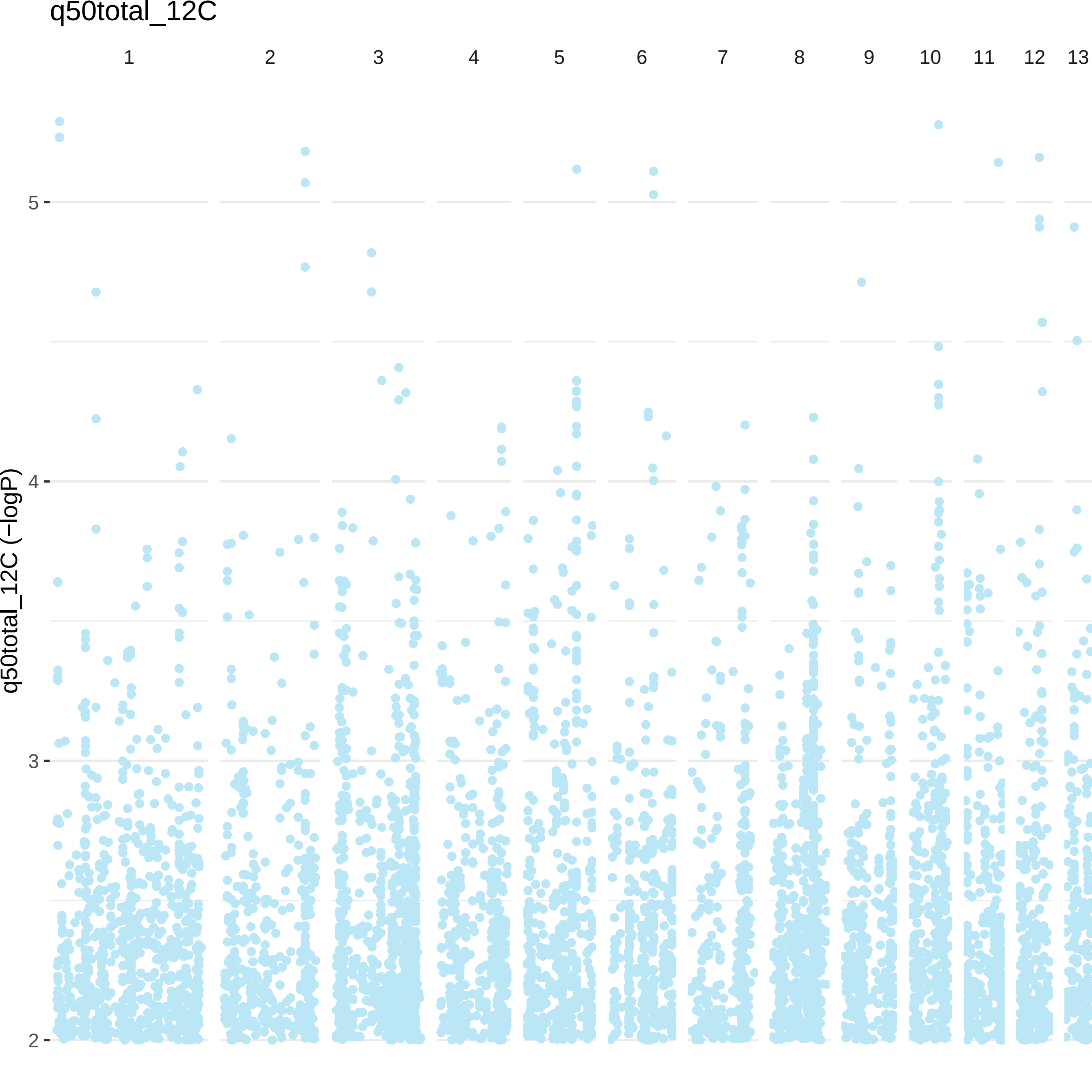

q50total_12C
1
2
3
4
5
6
7
8
9
10
11
12
13
5
4
3
2
q50total_12C (−logP)

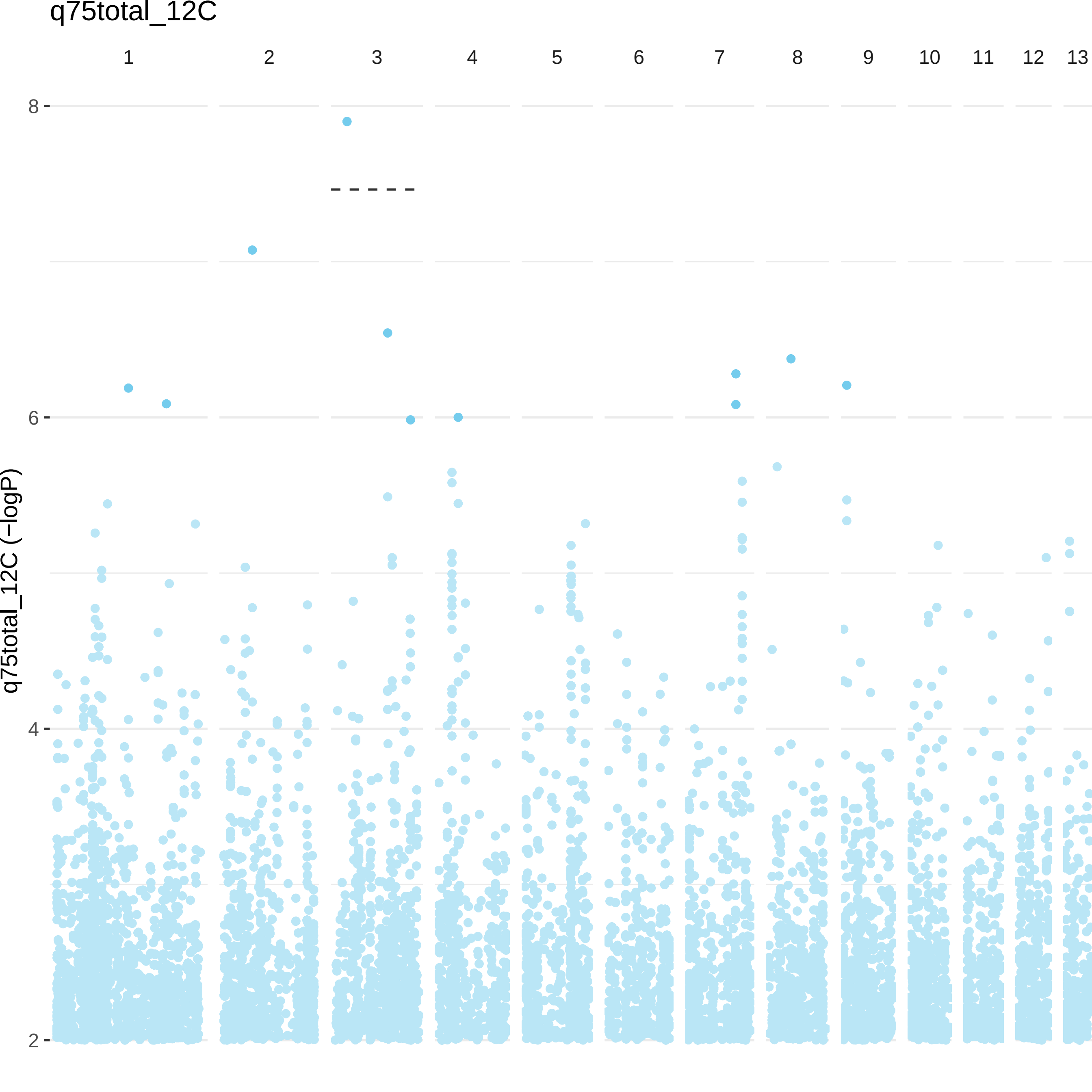
q75total_12C
1
2
3
4
5
6
7
8
9
10
11
12
13
8
6
4
2
q75total_12C (−logP)

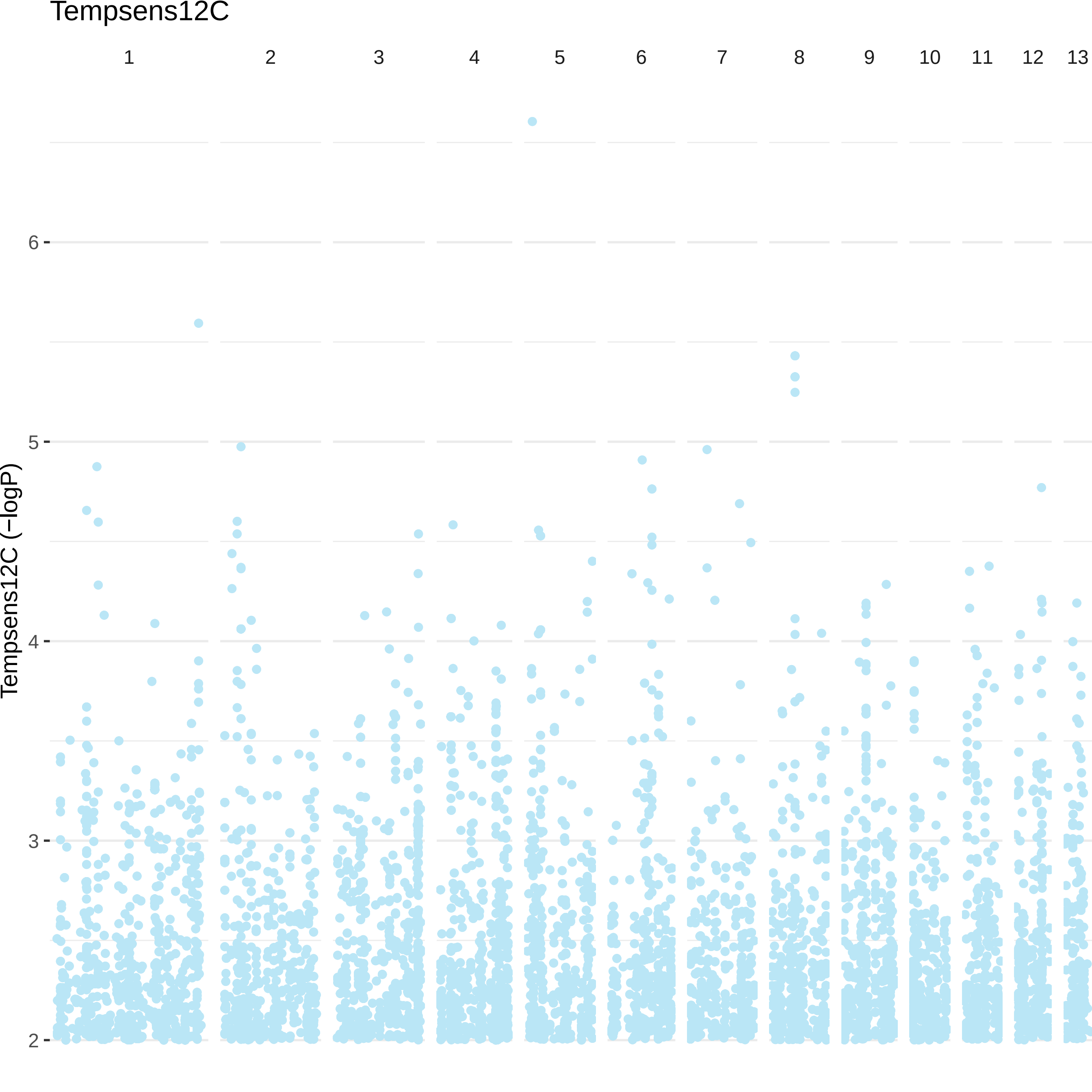
Tempsens12C
1
2
3
4
5
6
7
8
9
10
11
12
13
Tempsens12C (−logP)
6
5
4
3
2
