## Supplementary File S1 for "Thermal adaptation in worldwide collections of a major fungal pathogen"

**Supplementary File S1.** Detailed description of the Supplementary Tables.

For all the growth variables names there are short names in the Supplementary Material. As a reference here there is an explanation of these short names:

- approx_area : eAUC, empirical area under the curve
- growth_rate: maximum growth rate
- inflection: inflection time
- asymptote: asymptote of the logistic model, carrying capacity
- q25total: time to the first quantile of the concentration
- q50total: time to the second quantile of the concentration
- q75total: time to the third quantile of the concentration
- day2: spore concentration after 48 hours
- day4: spore concentration after 96 hours
- day6: spore concentration after 144 hours
- OptTemperature: optimal temperature
- Tempoptsens: temperature sensitivity
- Date: it is the date and time that the measurements were taken in the Tecan machine. It is in the format MM/DD/YYYY HH:MM:SS.

-Date: it is the date and time that the measurements were taken in the Tecan machine. It is in the format MM/DD/YYYY HH:MM:SS.

- Plate: it is the code given to keep track the position of the strains, replicates and dilutions in the microtiter plates. The names are coded by Plate###. Calibration plates (ODLines/ODLine) have two plate code numbers since there are two plates per each experimental week that were created on the same day and that harbour the 20 strains in 8 serial dilutions (look the methods section). Each column of the microtiter plate represents the 8 dilutions for each strain (except columns 1 and 12 which are filled with just media). The phenotyping plates have 4 cells of the microtiter plates with 2.5*10^5 spores/mL per each strain from columns 2-11 (columns 1 and 12 which are filled with just media). Therefore, the phenotyping plates have each 20 strains and there is a plate for each of the temperatures (12C, 15C, 18C, 22C, 25C). For example, the same batch of 20 strains, have 3 plate code names attatched to them: Plate020 for the 5 phenotyping microtiter plates (one per each temperature condition), Plate021 with 10 of the strains and the 8 dilutions, and Plate022 with the 10 other strains and the 8 dilutions.

- Timepoint: it is a code of the timepoint of each measurement in the fashion of t##. The t00 is the first measurement just right after having created the calibration plates and phenotyping plates. The t01-t12 are the following measurements and they were done only for the phenotyping plates (2 times per day for a total of 144 hours).

- Temperature: the temperature at which the phenotyping plates were placed right after the t00. The calibration plates, therefore, have "NO" in this column, since they were only measured at t00.

- Condition: represents the temperature condition that the phenotyping plates were placed after t00 and for the calibration plates it says either "ODLine" or "ODLines" which signifies that these are the calibration plates.

- Repet: it is numbered by 1-3. Represent the run of the machine, since for each timepoint we took 3 replicate measurements from the Tecan machine to address possible machine errors.

- Row: represents the row of the 96-well place microtiter plate. There are 8 rows represented by a letter from A-H.

- Col: represents the column of the 96-well place microtiter plate. There are 12 columns represented by a number from 1-12.

- Absorbance: it is the optical density at 405nm value for that particular cell. Values range from 0-2.

- Cell: is the exact microtiterplate cell (combination of row and column).

- Reduced_strainname: it is a shorter name for the strain names that normally does not include the year and location the strain is from. Some values of this column are named

"Negative_control" and this means that this cell in the microtiter plate did not contain any strain, but only had media.

- Strain: it is the name of the strain, which normally includes the reduced strain name and the year and location that the strain is from. Some values of this column are named "Negative_control" and this means that this cell in the microtiter plate did not contain any strain, but only had media.

- Continent: represents the large scale geographic region that the strain is from. There are 6 geographic regions: Africa, Middle_East, Europe, North_America, South_America and Oceania. Some values of this column are named "Negative_control" and this means that this cell in the microtiter plate did not contain any strain, but only had media.

- ID_genome_file: it is the name that the strains are found in SRA NCBI.

- Cluster: represents the results from snmf analysis by representing the strains that belong to the 8 genetic clusters (here represented by V# with # from 1-8) that were found and it is a category only present for strains that had an admixture coefficient higher than 0.8.

- Admix_coef: it is the admixture coefficient estimated by the snmf analysis and it is only represented for strains with a coefficient higher than 0.75.

- K: represents the number of genetic clusters that were found to be most suitable by the snmf analysis which was 8 (only present for strains with admixture coefficient higher than 0.8).

- Isolate_full_name_phenotyping: it is the same column as the "Strain" column but it only shows the names of the strains that had an admixture coefficient higher than 0.75.

- fordisplay: it is the "Continent" column combined with the "Location_year" column. It is a column for graphic display of the data.

- Adm_Cluster: it is a common name given according to which of the "Cluster" column categories are related to their locations. The categories are Argentina_Uruguay, Chile, Australia, Middle_East, Tunisia, USA, Europe and Canada.

- Latitude: the decimal value for the latitude of the location from which the strains' population is from.

- Longitude: the decimal value for the latitude of the longitude from which the strains' population is from.

- Variety: the wheat type or variety (when the information was available) that the strains' population was isolated from.

- Location: country or state (for the USA) that the strains' populations came from.

- Country: country from which the strains' populations came from.

- Geographic_region: exactly the same as the "Continent" column.

- Year: year in which the strains' populations were collected.

- Location_year: a combination of columns "Location" and "Year".

- KGnum: Koppen-Geiger climate number (established by the website World Maps of Koppen-Geiger Climate Classification, https://koeppen-geiger.vu-wien.ac.at/present.htm). They give the script Map_KG-Global.R and the raster files to recreate the KG-climate map in R.

- climate: it is the "KGnum" but now instead of numbers per climate class, it is the actual code for each climate class. The code consists of 3 letters, the first letter represents the main climate group (A:Tropical; B:Arid; C:Temperate; D:Continental; E:Polar). The second letter refers to the seasonal precipitation type and the third letter indicates the level of heat.

- Reason_removed: these are cell values to be removed and were filtered out of the analyses due to multiple reasons. The reason "No_genome_file" are cell values for strains that were phenotyped but that were not sequenced. The reason "Negative_control_contamination" are cell values for the media only cells that have a higher absorbance value than 6 times the standard deviation of the media only wells at the last timepoint measured. The reason "Wrong_genetic_cluster_or_missing_data" are cell values for strains that were identified through the snmf analysis to be misslabelled when sequencing or that were found to have more than 80% of missing genomic data. The reason "IQR_run_artifact" are cell values that were found to have through upper Inter Quartile Range filtering that the differences between this machine run and the other two (column "Repet") were too different, therefore, most likely a machine error. The reason "IQR_median_well" are cell values of cells that when compared to the other 3 cells for the same strain are outliers through upper Inter Quartile Range filtering, which most likely represents pipetting errors. More detailed information on all these filtered data can be found in the Materials and Methods of the research article.

- Absorbance_control: represents the mean absorbance values for all the only media wells per each "Plate", "Timepoint", "Date" and "Condition". This was only calculated for phenotyping microtiter plates.

- Norm_absorbance: it is the normalized absorbance, which are the cell values of the "Absorbance" per each "Cell", "Strain", "Plate", "Timepoint", "Date" and "Condition", minus the "Absorbance_control". This was only calculated for phenotyping microtiter plates.

- Slope_calcurve: represents the slope of the linear regression of the 8 dilutions and the respective absorbance values of the calibration microtiter plate results for the strain in that row of the file.

- intercept: represents the value of y for when x=0 of the linear regression of the 8 dilutions and the respective absorbance values of the calibration microtiter plate results for the strain in that row of the file.

- Adj_rsquared_calcurve: represents the adjusted R-squared of the linear regression of the 8 dilutions and the respective absorbance values of the calibration microtiter plate results for the strain in that row of the file. The closest it is to 1, better the fit of the linear regression.

- Concentration: it is the converted absorbance to spore concentration value through the linear regression of the 8 dilutions of the respective absorbance values of the calibration plates results of the strain that corresponds.

- RemoveIQR: the value "Yes" for this column is where in the column "Reason_removed" have the value "IQR_run_artifact" that were found to have through upper Inter Quartile Range filtering that the differences between this machine run and the other two (column "Repet") were too different, therefore, most likely a machine error. The value "No" for this column means that these cells should not be filtered out.

- RemoveIQRmed: the value "Yes" for this column is where in the column "Reason_removed" have the value "IQR_median_well" are cell values of cells that when compared to the other 3 cells for the same strain are outliers through upper Inter Quartile Range filtering, which most likely represents pipetting errors. The value "No" for this column means that these cells should not be filtered out.
- New_Location_year: It includes at least two different terms to signify the population it comes from. The first term is the Country of origin and the second term is the city or the most precise location information available. The populations from the USA have an extra term, which is the code for the State that they come from. Then, sometimes the month of collection is included as an extra term if the populations were obtained from the same field at different time points.
